# A Platform for Controlled Template-Independent Enzymatic Synthesis of RNA Oligonucleotides and Therapeutics

**DOI:** 10.1101/2023.06.29.547106

**Authors:** Daniel J. Wiegand, Jonathan Rittichier, Ella Meyer, Howon Lee, Nicholas J. Conway, Daniel Ahlstedt, Zeynep Yurtsever, Dominic Rainone, Erkin Kuru, George M. Church

**Affiliations:** Department of Genetics, Harvard Medical School; Boston, Massachusetts 02115, USA; Wyss Institute for Biologically Inspired Engineering; Boston, Massachusetts 02115, USA; EnPlusOne Biosciences, Inc.; Watertown, Massachusetts 02472, USA

## Abstract

Therapeutic RNA oligonucleotides have shown tremendous potential to manage and treat disease, yet current manufacturing methods cannot deliver on this promise. Here, we report the development and optimization of a novel, aqueous-based, template-independent enzymatic RNA oligonucleotide synthesis platform as an alternative to traditional chemical methodologies. Our platform is made possible by reversible terminator nucleoside triphosphates and an enzyme capable of their incorporation. We show that many common therapeutic RNA modifications are compatible with our process and demonstrate the enzymatic synthesis of natural and modified oligonucleotides in both liquid and solid phases. Our platform offers many unique advantages over chemical synthesis, including the realization of a more sustainable process to produce therapeutic RNA oligonucleotides.

**One-Sentence Summary:** An enzyme and novel nucleotide building blocks are used to synthesize RNA oligonucleotides template independently under aqueous conditions.

## Main Text

The synthesis of RNA oligonucleotides by the phosphoramidite chemical method has enabled many valuable discoveries and novel ways to treat disease throughout the past 50 years (*1*–*8*). This has culminated in an immense demand for large-scale manufacturing of therapeutically relevant oligonucleotides; however, the future promise of RNA is limited by the production techniques of the past (*9*–*11*). Enzymatically synthesizing oligonucleotides over traditional chemical methods holds tremendous potential to deliver high-quality RNA to the world (*12*–*15*). In addition to improving oligonucleotide synthesis yield and purity, adoption of enzymatic methods can also simplify downstream purification, eliminate large-scale consumption of organic solvents, and prevent the accumulation of toxic by-products, thus reducing the environmental impact of RNA production (*16, 17*). Here, we describe the development of a novel, aqueous-based enzymatic synthesis platform with the capacity to write natural and modified RNA oligonucleotides one base at a time without the need of a template sequence.

Our novel platform synthesizes RNA oligonucleotides over a series of iterative reaction cycles in the liquid bulk phase or on solid supports in a controlled, template-independent manner. Synthesis occurs in the 5’- to 3’-direction and three primary components are necessary for synthesis: (1) reversible terminator nucleoside triphosphate (NTP) building blocks, (2) an enzyme capable of their efficient incorporation, and (3) a pre-existing oligonucleotide to initiate controlled synthesis **(Fig. 1A)**. We utilize mutant variants of CID1 Poly(U) Polymerase (PUP) derived from the fission yeast *Schizosaccharomyces pombe* to write RNA oligonucleotides (*18*–*20*). Our mutants have an increased ability to incorporate NTP bases indiscriminately in comparison to their wild-type counterpart **(Fig. S1)**. The initiator oligonucleotide, which is essential for enzymatic functionality and controlled, template-independent synthesis, should be at least 10 nucleotides (nt) or greater in length and can be either a homopolymeric string of bases or a rationally designed sequence (**Fig. S2, S3)**.

**Fig. 1.**
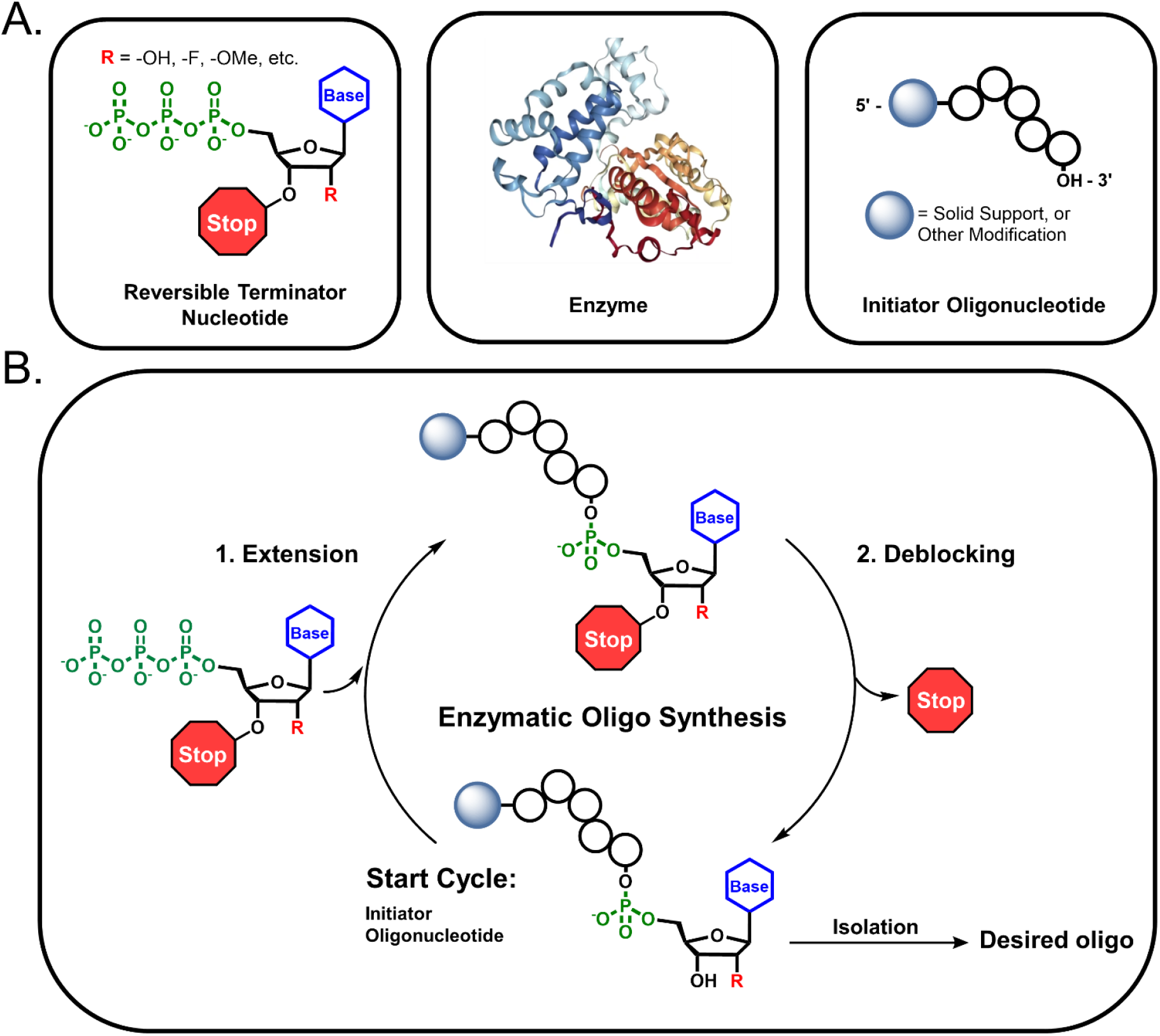
General overview of the controlled, template-independent enzymatic RNA oligonucleotide synthesis process. **(A)** Three primary components are required for carrying out an enzymatic extension: 3’-blocked reversible terminator nucleotides, an enzyme capable of their robust and indiscriminate incorporation, and an initiator oligonucleotide. The reversible terminator group stops uncontrolled polymerization by the enzyme and limits extension to a single incorporation event. The initiator oligonucleotide may be variable in terms of sequence and length. It can also be bound to a solid support or feature other modifications such as 5’-fluorophore or functional handle. **(B)** A typical cycle of enzymatic synthesis begins with (1) extension of the initiator oligonucleotide in the presence of a reversible terminator NTP and enzyme. A deblocking step (2) then occurs to remove the reversible terminator group from the extended oligonucleotide allowing the next cycle of synthesis to commence. After reaching the desired length and composition, the final oligonucleotide product is isolated.

A typical reaction cycle is similar to that of chemical phosphoramidite synthesis (*21*), except that there are only two main steps per cycle: extension and deblocking **(Fig. 1B)**. During an extension step, the enzyme incorporates the desired reversible terminator NTP onto the 3’-terminus of the initiator oligonucleotide. A successful extension step results in the generation of an N+1* product, where N and the asterisk represent the length of the initiator oligonucleotide and presence of a reversible terminator blocking group, respectively. Next, the N+1* oligonucleotide undergoes deblocking, where the reversible terminator is removed with a mild hydrolysis reagent, yielding an unblocked N+1 product and allows for the subsequent cycle of controlled, enzymatic synthesis to commence. The process of iterative extension and deblocking is repeated until the desired full-length RNA oligonucleotide is synthesized. The product can then be released enzymatically from the initiator or solid support for isolation. Unlike chemical-based RNA oligonucleotide synthesis, a final global deprotection is not required with our enzymatic method (*22*).

### Development and Evaluation of Reversible Terminator Nucleotide Building Blocks

The key to building our enzymatic RNA synthesis platform was the development of reversible terminator nucleotides with a blocking group that can be efficiently removed without the formation of reactive side products. An ideal blocking group is one that is stable during enzymatic extension reactions and under long-term storage conditions (*23*). It should also be a small moiety to limit perturbation of enzymatic substrate binding to ensure efficient nucleotide incorporation onto the growing oligonucleotide during synthesis. Although several viable options were considered (*24*–*26*), we ultimately decided that an allyl ether (-OCH_2_CHCH_2_) blocking group was the most ideal option for meeting the aforementioned criteria. This choice was further supported by previous work demonstrating quantitative allyl ether deblocking using palladium (Pd) as a catalyst and triphenylphosphine trisulfonate (TPPTS) in buffered aqueous solutions (*27*–*29*). The versatility and selectivity of Pd as a catalyst has enabled the manufacturing of many pharmaceuticals and fine chemicals at the kilogram scale in general (*30*).

Next, we needed to decide where to install the allyl ether blocking group on the NTP. Because PUP catalyzes the uncontrolled polymerization of long homo- and heteropolymers in the presence of NTPs with a free 3’-hydroxyl (-OH), installing the allyl ether group at this position was crucial to limiting extension to a single incorporation. Blocking at the 3’-sugar position would also enable the use of 2’-modifications such as 2’-fluoro (-F), 2’-methoxy (-OMe), and 2’-methoxyethyl (- MOE), which are important to the functionality of many therapeutic and antisense oligonucleotides (*31*). Thus, we accessed the complete set (A, U, G, C) of 3’-O-allyl ether reversible terminator NTPs using established methods for nucleoside preparation and triphosphorylation (**Sch. I-IV**). Since our enzymatic reaction conditions are significantly milder compared to chemical phosphoramidite synthesis, we did not need to protect the base or phosphate of the NTP with acetyl, benzoyl, or 2-cyanoethyl groups (*32*). Chemical synthesis especially struggles with RNA because of the need to protect the 2’-OH with t-butyldimethylsilyl (TBDMS) or triisopropylsilyloxymethyl (TOM) (*33*–*35*). Due to our mild conditions, we also leave the 2’-OH unprotected.

Following a liquid bulk phase reaction scheme (**Fig. 2A**), we evaluated the capacity of our 3’-O-allyl ether NTPs to undergo a single cycle of enzymatic RNA synthesis. Post-reaction analysis of enzymatic extension using matrix-assisted laser desorption/ionization-time of flight (MALDI-TOF) mass spectrometry showed the absence of initiator oligonucleotide and presence of the desired N+1* product for each reversible terminator NTP (**Fig. 2B**). These results were supported by liquid chromatography mass spectrometry (LC/MS), which showed coupling efficiencies > 95% for each blocked NTP **(Fig. S4**). MALDI-TOF and LC/MS also confirmed the removal of the allyl ether group upon deblocking for each reversible terminator NTP (**Fig. 2B, S5**). A time-course analysis showed that extension reactions are completed within the first minutes of incubation (**Fig. 2C**); however, enzymatic processing was found to be dependent on the initiator concentration and nucleobase (**Fig. S6**). In assessing the product impurity profiles of our initial extension and deblocking reactions, we found that the majority to be buffer components and additives used in extension and isolation steps, respectively. In general, these impurities eluted well before extended or deblocked oligonucleotide product as shown by LC, indicating an overall good isolated purity in all cases (**Fig. S4, S5**).

**Fig. 2.**
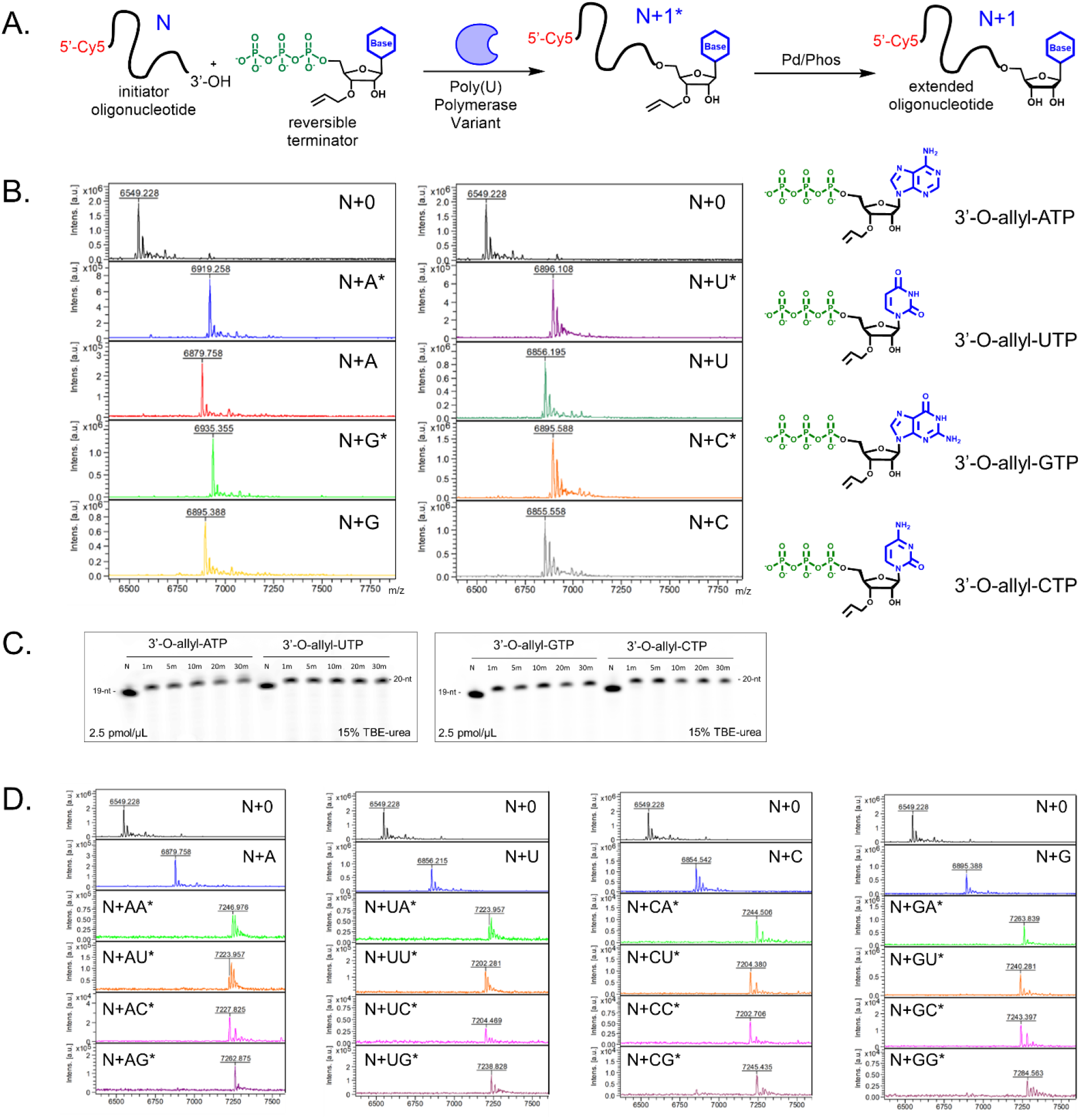
Initial evaluation of 3’-O-allyl ether reversible terminator NTPs as building blocks for controlled, enzymatic RNA oligonucleotide synthesis. **(A)** A complete set (A, U, G, C) of 3’-O-allyl ether NTPs were tested for enzymatic incorporation and deblocking using a liquid bulk phase reaction scheme, where N is the length of the initiator, N+1* is the extension intermediate with the 3’-O-allyl ether group as represented by the asterisk, and N+1 is the deblocked product for each base. **(B)** MALDI-TOF mass spectrometry was used to verify NTP extension to N+1* by the Poly(U) mutant variant and subsequent deblocking of the allyl ether group to N+1; the masses of all resultant oligonucleotides are given and compared to the 19-nt initiator. (**C**) A kinetic profile for each 3’-O-allyl ether NTP was obtained and analyzed with denaturing gel electrophoresis;reaction samples were taken at 1-, 5-, 10-, 20-, and 30-minutes. Control reactions (N) included all reaction components except NTP. **(D)** MALDI-TOF was used to assess the efficiency of two controlled, enzymatic synthesis cycles in which all N+2* combinations of base extensions were produced; the masses of all resultant oligonucleotides are given and compared to the 19-nt initiator.

### Multi-Cycle Enzymatic Synthesis of Natural RNA Oligonucleotides

With our reversible terminator NTP building blocks characterized, we next sought to prove that multi-cycle enzymatic synthesis of longer RNA oligonucleotides was possible with our platform. To do this, we first generated oligonucleotide extension products that constituted all possible 16 N+2* base transitions (e.g., A to A, G to U, *etc*.) under standard extension and deblocking reaction conditions. We previously found that certain base transitions can be troublesome for template-independent polymerases (*36*); however, analysis with MALDI-TOF confirmed the formation of all intended products as evidenced by the total consumption of the initiator oligonucleotide and deblocked N+1 during the first and second cycles of synthesis, respectively **(Fig. 2D)**. All N+2* base transitions were achieved at high efficiency without needing to alter any reaction components or increase incubation times for extension or deblocking steps.

We next turned to performing 5x cycles of controlled, enzymatic synthesis to produce an N+5* oligonucleotide with the natural RNA sequence N+U-U-U-C-G* in the liquid bulk phase using a Cy5 labeled initiator (**Fig. 3A**). To achieve longer synthesis lengths, we increased the initial scale to approximately 20 nmol in a volume of 8 mL and adjusted the extension and deblocking volumes accordingly after each cycle to maximize the efficiency of enzymatic coupling by maintaining standard reaction conditions (e.g., 2.5 pmol/μL oligonucleotide). MALDI-TOF analysis after each cycle showed the successful formation of all extended and deblocked products, indicating a high coupling efficiency over the course of enzymatic synthesis (**Fig. 3B**). This was confirmed with LC analysis, where we found excellent isolated purity of the oligonucleotide intermediates and final product (as measured by 649 nm for Cy5) (**Fig. 3C, Fig. S7A, B**).

**Fig. 3.**
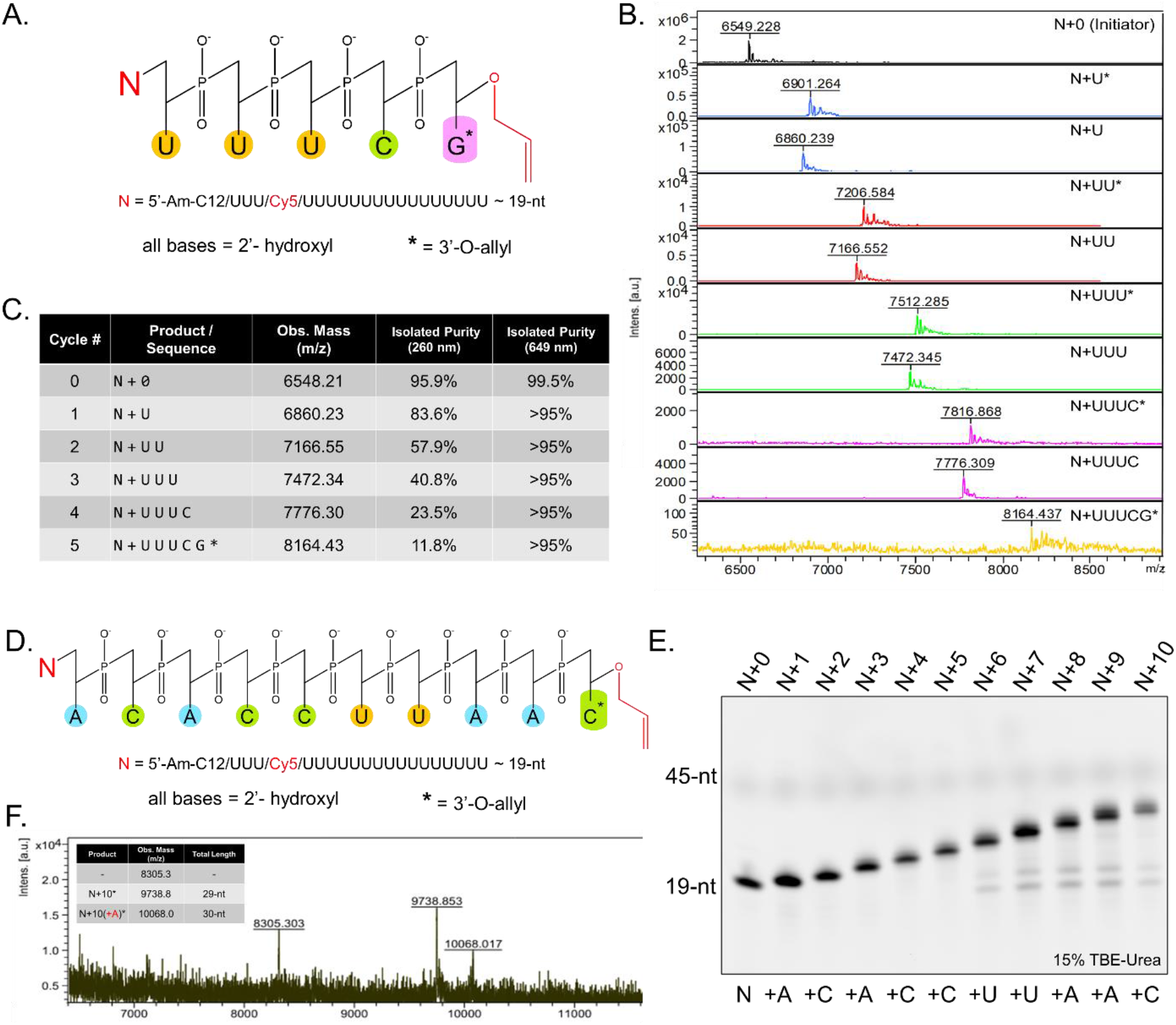
Results of multi-cycle enzymatic synthesis to produce natural RNA oligonucleotides using the 3’-O-allyl ether NTP set. **(A)** An N+5 RNA oligonucleotide with the sequence N+U-U-U-C-G* was produced in the liquid bulk phase, where the asterisk represents a 3’-O-allyl ether group. **(B)** MALDI-TOF mass spectrometry was used to track the outcome of the extension and deblocking steps during each cycle of enzymatic synthesis. **(C)** The isolated purity of the growing oligonucleotide and final product was determined after each cycle using LC/MS at 260 nm and 649 nm, which are summarized in the table. **(D)** An N+10 RNA oligonucleotide with the sequence N+A-C-A-C-C-U-U-A-A-C* was also produced in the liquid bulk phase. **(E)** High resolution gel electrophoresis was used to analyze the success of each cycle after the sequence was enzymatically synthesized with an imager set to collect Cy5 signal. **(F)** The final oligonucleotide N+10* product was assessed with MALDI-TOF and summarized along with any major impurities detected.

Following a successful N+5 synthesis, we next attempted to enzymatically synthesize an N+10* oligonucleotide with the natural RNA sequence N+A-C-A-C-C-U-U-A-A-C* (**Fig. 3D**). Tracking synthesis with high resolution gel electrophoresis showed formation of all extension intermediates and the N+10* final product (**Fig. 3E**), which had an isolated purity of 67% (as determined by LC at 649 nm) (**Fig. S8**). MALDI-TOF analysis showed the expected mass of the N+10* (9738.8 m/z) in addition to an N+11* impurity (10068.0 m/z) with an extra “A” in the sequence (**Fig. 4F**). A potential explanation is ATP carry-over during the later cycles of synthesis. Another impurity with a mass of 8305.30 m/z was observed with MALDI-TOF; additional characterization is required to determine its exact composition.

**Fig. 4.**
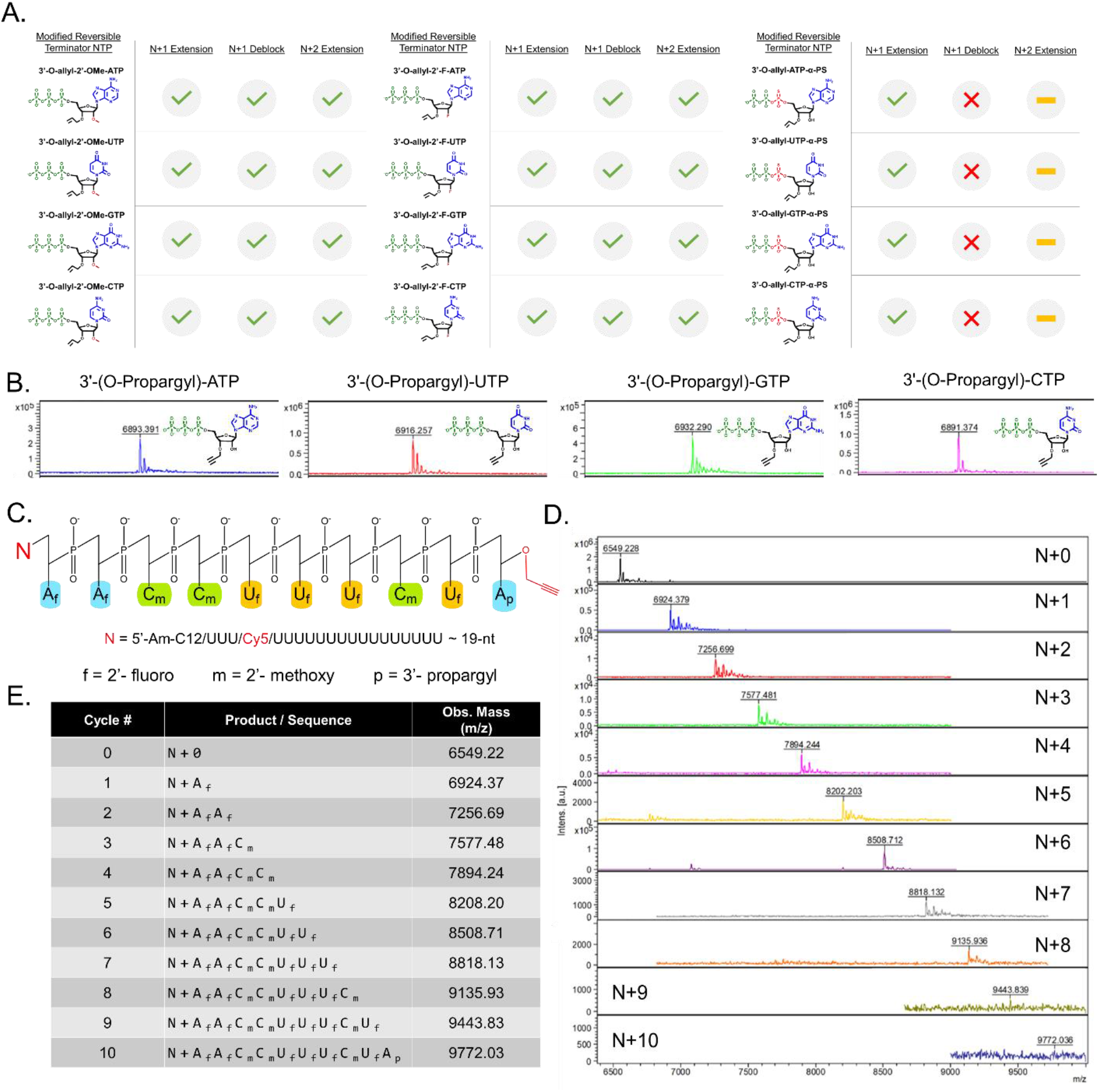
Compatibility summary of modified 3’-O-allyl ether reversible terminator NTP sets and results of multi-cycle synthesis to produce a fully modified RNA oligonucleotide. **(A)** Modified 3’-O-allyl ether reversible terminator NTPs were evaluated by performing an initial N+1* extension, deblocking reaction, and an N+2* extension. A green checkmark indicates a successful reaction and a red cross-out indicates an unsuccessful reaction. Reactions that were not attempted are indicated by a yellow bar. Evaluation of each individual cycle step was determined by MALDI-TOF mass spectrometry (**Fig. S11-13**). (**B**) MALDI-TOF assessment of enzymatic extension reactions using a set of 3’-O-propargyl ether NTPs (A, U, G, C) to install a functional handle onto oligonucleotides. **(C)** A fully modified N+10 oligonucleotide with the sequence N+A_f_-A_f_-C_m_-C_m_-U_f_-U_f_-C_m_-U_f_-A_p_ was synthesized using modified reversible terminator NTPs, where f = 2’-fluoro, m = 2’-methoxy and p = 3’-O-propargyl. **(D)** MALDI-TOF mass spectrometry was used to verify extension using the modified reversible terminators NTPs during each cycle of enzymatic synthesis. (**E**) The expected oligonucleotide sequences and observed masses from MALDI-TOF are summarized in the table.

### Enzymatic Incorporation of NTPs with Therapeutic Modifications

While the benefits of template independent enzymatic synthesis of natural RNA oligonucleotides are numerous, all commercial RNA-based therapeutics are partially or fully modified (*37*). We therefore accessed sets of modified 3’-O-allyl ether reversible terminator NTP sets with either a 2’-F, 2’-OMe, or alpha-phosphorothioate (α-PS) modification. We evaluated the capacity of each modified reversible terminator NTP to control enzymatic synthesis by generating all single base transition (e.g., A_m_ to A_m_, C_f_ to C_f_, *etc*. where **f** is a 2’-F modification, **m** is a 2’-OMe) N+2* extension products for each set (**Fig. 4A**). The formation of all expected oligonucleotide products with 2’-F and 2’-OMe modifications was observed with MALDI-TOF analysis using standard reaction conditions (**Fig. S9, S10**). Interestingly, we found that α-PS modified NTPs can be enzymatically incorporated by our enzyme; however, deblocking with the allyl ether Pd/TPPTS chemistry resulted in the formation of reduction side products (**Fig. S11**) (*38*). Despite efforts to prevent degradation of the PS during deblocking, we were unsuccessful and therefore could not move on to obtain the desired N+2* product. Further investigation into alternative deblocking conditions is warranted for our synthesis platform to better handle PS modifications.

Strong and indiscriminate incorporation of all modified reversible terminator NTPs by our enzyme was further exemplified by the generation of long homopolymer sequences in the presence of their unblocked counterparts **(Fig. S12**). Similar results were found when we tested various propargyl modified nucleotides with the intention of installing functional handles onto our oligonucleotide products. These handles provide a way to conjugate enzymatically synthesized oligonucleotides with important ligands such as N-Acetylgalactosamine (GalNAc), which is commonly used to deliver therapeutic oligonucleotides to the liver (*39, 40*). Uncontrolled polymerization using unblocked N^6^-propargyl-ATP and 2-Ethynyl-ATP resulted in the generation of long homopolymer sequences >100-nt, whereas a set of 3’-propargyl ether modified NTPs yielded N+1 single extension products for each base (**Fig. 4B, S13**). Since a single terminal propargyl group was the preferred result of this activity, we labeled the functionalized N+1 oligonucleotides with α-GalNAc-PEG3-Azide using a standard click chemistry protocol. MALDI-TOF analysis indicated complete conjugation, marked by the total consumption of the unlabeled material (**Fig. S14**). We did not label the homopolymer sequences generated by N^6^-propargyl-ATP and 2-Ethynyl-ATP with the GalNAc ligand. However, our capacity to readily generate these sequences enables further exploratory opportunities in nucleic acid-based materials (*41*) as well as the modulation of mRNA stability with modifications to the polyA tail (*42*).

Now that we had a full palette of modified reversible terminator NTPs at our disposal and proved out that our platform can accommodate nearly all of them, we set out to synthesize a fully modified RNA oligonucleotide of longer length as a final synthesis capstone. Starting with a 60 mL, 200-nmol liquid bulk phase reaction and using the standard Cy5-labeled initiator oligonucleotide, we performed 10x cycles of enzymatic synthesis to produce the sequence N+A_**f**_**-**A_**f**_**-**U_**m**_**-**U_**m**_**-**C_**f-**_C_**f**_**-**C_**f**_**-**U_**m**_**-**C_**f**_**-**A_**p**_, where **p** is a terminal 3’-propargyl (**Fig. 4C**). MALDI-TOF analysis, performed after each cycle, indicated the formation of all expected extension intermediates and final product (**Fig. 4D, E)**. The overall yield of synthesis was low with ≤50 pmol of final product, which may have impeded observation of any arising impurities on the MALDI-TOF during the last few cycles. Similarly, we were able to verify that the terminal propargyl group was enzymatically installed, but insufficient material made it challenging to label with a ligand like α-GalNAc-PEG3-Azide and to fully characterize the final product. Nonetheless, this capstone represents, to our knowledge, the first-ever controlled synthesis of a fully modified oligonucleotide with reversible terminator NTPs and a template-independent polymerase.

### Initiator Oligonucleotide Cleavage and Development of a Solid Support System

After successful synthesis of a fully modified RNA oligonucleotide with our platform, we found it crucial to develop a method to remove the undesired initiator sequence from the final product. To do this, we exploited the ability of the enzyme Endonuclease V from *E. coli* to cleave single-stranded oligonucleotides in a site-specific manner (*43*). Endonuclease V cleavage is mediated by the recognition of an inosine placed rationally within the initiator sequence (*44*). Cleavage occurs two bases downstream of the inosine and products are then left with a 5’-phosphate (PO) group, which can be removed with a phosphatase (**Fig. S15, S16**). An Endonuclease V cleavage site can be either pre-installed during initiator production or added by the enzymatic incorporation of an inosine reversible terminator nucleotide. To demonstrate this, we accessed a 3’-O-allyl ether inosine building block and validated its general functionality as a reversible terminator similarly to that of our 2’-sugar modified NTP variants (**Fig. S17A**). We then performed uncontrolled polymerization off of an N+I deblocked oligonucleotide product to generate a long homopolymer and verified that the initiator sequence, which featured a 5’-Cy5 modification, could be cleaved from homopolymer product (**Fig. S17B)**. The desired oligonucleotide product can then be isolated from the initiator sequence through further enrichment or purification.

In addition to providing a method to remove the initiator sequence from the enzymatically synthesized oligonucleotide product, Endonuclease V-mediated cleavage formed the basis for the development of a solid-support system for our platform (**Fig. 5A**). A robust and economical solid support should enable the synthesis of longer oligonucleotides at commercially relevant scales in comparison to synthesis in the liquid bulk phase. To access such solid support, we added a Bis(NHS)PEG5 linker to a long chain alkylamine (LCAA) controlled pore glass (CPG) and then labeled it with a 5’-amine initiator oligonucleotide using NHS conjugation chemistry. We used an initiator oligonucleotide that had a pre-installed inosine and demonstrated that incubating labeled CPG in a stir reactor format with Endonuclease V cleaved the initiator at the desired location, releasing the oligonucleotide downstream of the inosine (**Fig. S18**). We next tested a single cycle of controlled extension and deblocking on the CPG solid-support using each 3’-O-allyl ether NTPs and found successful formation of the desired **N+1** products in all cases after Endonuclease V cleavage (**Fig. S19**).

**Fig. 5.**
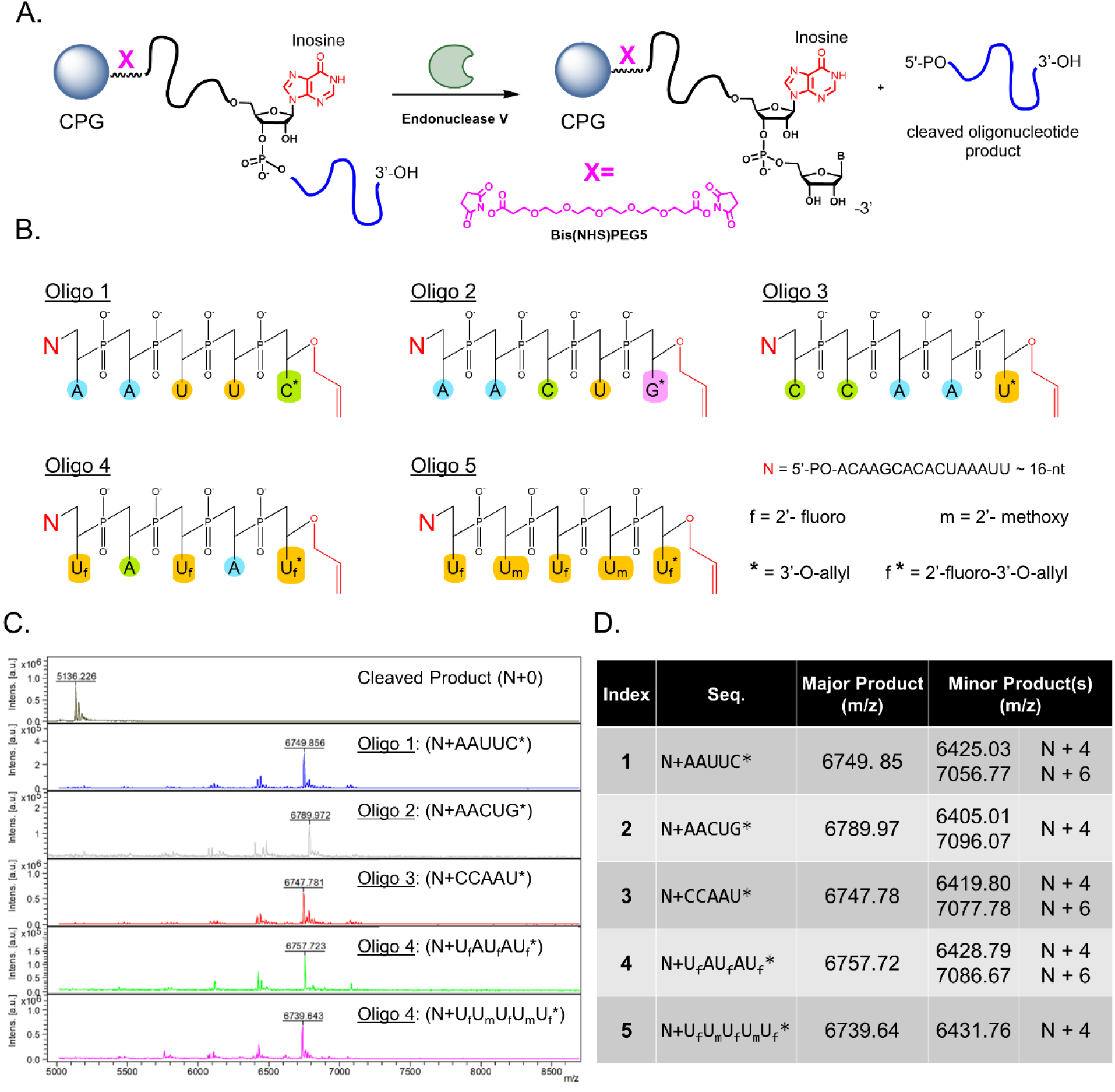
Overview and demonstration of a solid support system for controlled, enzymatic RNA oligonucleotide synthesis. Outlined in a general scheme **(A)**, an initiator oligonucleotide (black) is bound to LCAA-CPG using a Bis(NHS)PEG5 linker (pink) using NHS conjugation chemistry. The initiator harbors a deoxy- or riboinosine base (red) for recognition by *E. coli* Endonuclease V, which cleaves the desired oligonucleotide product (blue) immediately downstream of the inosine base. The oligonucleotide product can then be isolated from the CPG solid support. (**B**) To demonstrate the viability of the CPG solid support, 5x N+5 oligonucleotides were enzymatically synthesized in a stir reactor format. Their sequences were comprised of natural and modified bases, with one being partially modified with 2’-fluoro groups and another fully modified with both 2’-fluoro and 2’-methoxy. **(C)** MALDI-TOF mass spectrometry was used to evaluate the oligonucleotide material cleaved from the solid support. **(D)** A summary of enzymatic synthesis is given, including a high-level description of the major and minor products found.

Building on these experiments, we performed controlled enzymatic synthesis of 5x unique **N+5** oligonucleotides with unmodified, partially modified, and fully modified sequences on our solid support (**Fig. 5B**). For these syntheses, the inosine base was present in the surface bound initiator oligonucleotide. In general, it was observed that all 5 RNA oligonucleotides were successfully synthesized upon cleavage from the solid support. MALDI-TOF analysis showed the major peak corresponded to the expected masses of the intended sequences (**Fig. 5C**). The primary impurities were N+4 and N+6 products, which may have arisen from incomplete access of the enzyme to surface bound oligonucleotide or carry-over of NTPs from inadequate washing of the solid support (**Fig. 5D**). To better address and prevent such impurities, future work should involve packing the CPG solid support into a closed column with precise flow control.

## Discussion

In this work, we report the development and optimization of a novel platform for the controlled, enzymatic synthesis of natural and modified RNA oligonucleotide sequences in an aqueous based-format using 3’-O-allyl ether reversible terminators and an enzyme capable of their incorporation. Our efforts primarily focused on accessing and evaluating the performance of 24 unique NTP building blocks. We demonstrated several RNA syntheses, including a fully modified oligonucleotide with therapeutically important moieties. We also show the development of a CPG solid-support system capable of multi-cycle synthesis and enzymatic cleavage of natural RNA and modified oligonucleotides from the surface.

We found that our enzymatic platform is comparable to chemical synthesis methods in terms of building block coupling efficiencies and cycle times; however, our overall yields were lower than what is generally expected for RNA oligonucleotide synthesis using phosphoramidite chemistry (*45*). The primary culprit for observing lower than expected yields could be traced to inefficiencies in the retention of short oligonucleotides during liquid phase reaction purifications after enzymatic extension and deblocking. Our solid support system, which is similar to that of chemical synthesis, will be critical to retaining more desired oligonucleotide product from cycle to cycle and scaling of the enzymatic synthesis process to meet the demands of the oligonucleotide therapeutic market.

Although only a few enzymatic oligonucleotide synthesis technologies have emerged (*46, 47*), it is abundantly clear that there are many advantages for its use as an alternative to chemical-based methods. Not only will wide-ranging implementation of enzymatic synthesis greatly reduce environmental impact by furnishing a more sustainable and ecologically friendly process, but it will also remove the limitations that plague phosphoramidite chemistry. This will enable opportunities to explore modifications and structures that can be used in the discovery of novel, next-generation oligonucleotide therapeutics to treat rare diseases with personalized medicine (*48*) or the rapid, far-reaching deployment of RNA on a global scale (*49*).

## Acknowledgements

The authors thank R. Kohman, J. Tam, and members of the Church lab for their helpful discussions.

## Author Contributions

- Conceptualization: DJW, JR, EK, NC, GMC
- Methodology: DJW, JR, EM, HL
- Investigation: DJW, JR, EM, HL, DA, ZY, DR
- Writing – original draft: DJW, JR
- Writing – review & editing: DJW, JR, EM, DA, ZY
- Project Supervision: DJW, EK, GMC

## Supplementary Materials

Materials & Methods

Figures S1-S19

Preparation of 3’-O-allyl ether NTPs (A, U, G, C)

NMR Analysis of 3’-O-allyl ether NTPs (A, U, G, C)

References

## References

1. A. Khvorova, J. K. Watts, The chemical evolution of oligonucleotide therapies of clinical utility. Nat. Biotechnol. 35, 238–248 (2017).

2. S. T. Crooke, B. F. Baker, R. M. Crooke, X.-H. Liang, Antisense technology: an overview and prospectus. Nat. Rev. Drug Discov. 20, 427–453 (2021).

3. S. L. Beaucage, Solid-phase synthesis of siRNA oligonucleotides. Curr. Opin. Drug Discov. Devel. 11, 203–216 (2008).

4. A. A. Levin, Treating disease at the RNA level with oligonucleotides. N. Engl. J. Med. 380, 57–70 (2019).

5. R. L. Setten, J. J. Rossi, S.-P. Han, The current state and future directions of RNAi-based therapeutics. Nat. Rev. Drug Discov. 18, 421–446 (2019).

6. F. Wang, T. Zuroske, J. K. Watts, RNA therapeutics on the rise. Nat. Rev. Drug Discov. 19, 441–442 (2020).

7. K. K. Ray, U. Landmesser, L. A. Leiter, D. Kallend, R. Dufour, M. Karakas, T. Hall, R. P. T. Troquay, T. Turner, F. L. J. Visseren, P. Wijngaard, R. S. Wright, J. J. P. Kastelein, Inclisiran in Patients at High Cardiovascular Risk with Elevated LDL Cholesterol. N. Engl. J. Med. 376, 1430–1440 (2017).

8. D. R. Corey, Nusinersen, an antisense oligonucleotide drug for spinal muscular atrophy. Nat. Neurosci. 20, 497–499 (2017).

9. M. Catani, C. De Luca, J. Medeiros Garcia Alcântara, N. Manfredini, D. Perrone, E. Marchesi, R. Weldon, T. Müller-Späth, A. Cavazzini, M. Morbidelli, M. Sponchioni, Oligonucleotides: Current Trends and Innovative Applications in the Synthesis, Characterization, and Purification. Biotechnol. J. 15, e1900226 (2020).

10. A. G. Molina, Y. S. Sanghvi, Liquid-phase oligonucleotide synthesis: Past, present, and future predictions. Curr. Protoc. Nucleic Acid Chem. 77, e82 (2019).

11. Y. Huang, K. W. Knouse, S. Qiu, W. Hao, N. M. Padial, J. C. Vantourout, B. Zheng, S. E. Mercer, J. Lopez-Ogalla, R. Narayan, R. E. Olson, D. G. Blackmond, M. D. Eastgate, M. A. Schmidt, I. M. McDonald, P. S. Baran, A P(V) platform for oligonucleotide synthesis. Science. 373, 1265–1270 (2021).

12. M. A. Jensen, R. W. Davis, Template-Independent Enzymatic Oligonucleotide Synthesis (TiEOS): Its History, Prospects, and Challenges. Biochemistry. 57, 1821–1832 (2018).

13. K. J. D. Van Giesen, M. J. Thompson, Q. Meng, S. L. Lovelock, Biocatalytic synthesis of antiviral nucleosides, cyclic dinucleotides, and oligonucleotide therapies. JACS Au. 3, 13–24 (2023).

14. C. Schmitz, M. T. Reetz, Solid-phase enzymatic synthesis of oligonucleotides. Org. Lett. 1, 1729–1731 (1999).

15. N. Sabat, D. Katkevica, K. Pajuste, M. Flamme, A. Stämpfli, M. Katkevics, S. Hanlon, S. Bisagni, K. Püntener, F. Sladojevich, M. Hollenstein, Towards the controlled enzymatic synthesis of LNA containing oligonucleotides. Front Chem. 11, 1161462 (2023).

16. B. I. Andrews, F. D. Antia, S. B. Brueggemeier, L. J. Diorazio, S. G. Koenig, M. E. Kopach, H. Lee, M. Olbrich, A. L. Watson, Sustainability challenges and opportunities in oligonucleotide manufacturing. J. Org. Chem. 86, 49–61 (2021).

17. L. Ferrazzano, D. Corbisiero, A. Tolomelli, W. Cabri, From green innovations in oligopeptide to oligonucleotide sustainable synthesis: differences and synergies in TIDES chemistry. Green Chem. 25, 1217–1236 (2023).

18. O. S. Rissland, C. J. Norbury, The Cid1 poly(U) polymerase. Biochim. Biophys. Acta. 1779, 286–294 (2008).

19. P. Munoz-Tello, C. Gabus, S. Thore, Functional implications from the Cid1 poly(U) polymerase crystal structure. Structure. 20, 977–986 (2012).

20. B. M. Lunde, I. Magler, A. Meinhart, Crystal structures of the Cid1 poly (U) polymerase reveal the mechanism for UTP selectivity. Nucleic Acids Res. 40, 9815–9824 (2012).

21. M. H. Caruthers, Gene synthesis machines: DNA chemistry and its uses. Science. 230, 281–285 (1985).

22. F. Wincott, A. DiRenzo, C. Shaffer, S. Grimm, D. Tracz, C. Workman, D. Sweedler, C. Gonzalez, S. Scaringe, N. Usman, Synthesis, deprotection, analysis and purification of RNA and ribozymes. Nucleic Acids Res. 23, 2677–2684 (1995).

23. P. G. M. Wuts, T. W. Greene, “The Role of Protective Groups in Organic Synthesis” in Greene’s Protective Groups in Organic Synthesis (2006), pp. 1–15.

24. B. P. Stupi, H. Li, J. Wang, W. Wu, S. E. Morris, V. A. Litosh, J. Muniz, M. N. Hersh, M. L. Metzker, Stereochemistry of benzylic carbon substitution coupled with ring modification of 2-nitrobenzyl groups as key determinants for fast-cleaving reversible terminators. Angew. Chem. Int. Ed Engl. 51, 1724–1727 (2012).

25. F. Chen, M. Dong, M. Ge, L. Zhu, L. Ren, G. Liu, R. Mu, The history and advances of reversible terminators used in new generations of sequencing technology. Genomics Proteomics Bioinformatics. 11, 34–40 (2013).

26. M. Flamme, S. Hanlon, I. Marzuoli, K. Püntener, F. Sladojevich, M. Hollenstein, Evaluation of 3′-phosphate as a transient protecting group for controlled enzymatic synthesis of DNA and XNA oligonucleotides. Communications Chemistry. 5, 1–12 (2022).

27. H. Ruparel, L. Bi, Z. Li, X. Bai, D. H. Kim, N. J. Turro, J. Ju, Design and synthesis of a 3’-O-allyl photocleavable fluorescent nucleotide as a reversible terminator for DNA sequencing by synthesis. Proc. Natl. Acad. Sci. U. S. A. 102, 5932–5937 (2005).

28. J. Ju, D. H. Kim, L. Bi, Q. Meng, X. Bai, Z. Li, X. Li, M. S. Marma, S. Shi, J. Wu, J. R. Edwards, A. Romu, N. J. Turro, Four-color DNA sequencing by synthesis using cleavable fluorescent nucleotide reversible terminators. Proc. Natl. Acad. Sci. U. S. A. 103, 19635–19640 (2006).

29. J. Wu, S. Zhang, Q. Meng, H. Cao, Z. Li, X. Li, S. Shi, D. H. Kim, L. Bi, N. J. Turro, J. Ju, 3′-O-modified nucleotides as reversible terminators for pyrosequencing. Proceedings of the National Academy of Sciences. 104, 16462–16467 (2007).

30. C. Torborg, M. Beller, Recent applications of palladium-catalyzed coupling reactions in the pharmaceutical, agrochemical, and fine chemical industries. Adv. Synth. Catal. (2009) (available at https://onlinelibrary.wiley.com/doi/abs/10.1002/adsc.200900587).

31. G. F. Deleavey, M. J. Damha, Designing chemically modified oligonucleotides for targeted gene silencing. Chem. Biol. 19, 937–954 (2012).

32. A. Somoza, Protecting groups for RNA synthesis: an increasing need for selective preparative methods. Chem. Soc. Rev. 37, 2668–2675 (2008).

33. N. Usman, K. K. Ogilvie, M. Y. Jiang, R. J. Cedergren, The automated chemical synthesis of long oligoribuncleotides using 2’-O-silylated ribonucleoside 3’-O-phosphoramidites on a controlled-pore glass support: synthesis of a 43-nucleotide sequence similar to the 3’-half molecule of an Escherichia coli formylmethionine tRNA. J. Am. Chem. Soc. 109, 7845–7854 (1987).

34. S. Pitsch, P. A. Weiss, L. Jenny, A. Stutz, X. Wu, Reliable chemical synthesis of oligoribonucleotides (RNA) with 2′-O-[(triisopropylsilyl)oxy]methyl(2′-O-tom)-protected phosphoramidites. Helv. Chim. Acta. 84, 3773–3795 (2001).

35. M. H. Caruthers, A brief review of DNA and RNA chemical synthesis. Biochem. Soc. Trans. 39, 575–580 (2011).

36. H. Lee, D. J. Wiegand, K. Griswold, S. Punthambaker, H. Chun, R. E. Kohman, G. M. Church, Photon-directed multiplexed enzymatic DNA synthesis for molecular digital data storage. Nat. Commun. 11, 5246 (2020).

37. J. A. Kulkarni, D. Witzigmann, S. B. Thomson, S. Chen, B. R. Leavitt, P. R. Cullis, R. van der Meel, The current landscape of nucleic acid therapeutics. Nat. Nanotechnol. 16, 630–643 (2021).

38. D. R. Edwards, R. S. Brown, Development of metal-ion containing catalysts for the decomposition of phosphorothioate esters. Biochim. Biophys. Acta. 1834, 433–442 (2013).

39. J. K. Nair, J. L. S. Willoughby, A. Chan, K. Charisse, M. R. Alam, Q. Wang, M. Hoekstra, P. Kandasamy, A. V. Kel’in, S. Milstein, N. Taneja, J. O’Shea, S. Shaikh, L. Zhang, R. J. van der Sluis, M. E. Jung, A. Akinc, R. Hutabarat, S. Kuchimanchi, K. Fitzgerald, T. Zimmermann, T. J. C. van Berkel, M. A. Maier, K. G. Rajeev, M. Manoharan, Multivalent N-acetylgalactosamine-conjugated siRNA localizes in hepatocytes and elicits robust RNAi-mediated gene silencing. J. Am. Chem. Soc. 136, 16958–16961 (2014).

40. A. J. Debacker, J. Voutila, M. Catley, D. Blakey, N. Habib, Delivery of Oligonucleotides to the Liver with GalNAc: From Research to Registered Therapeutic Drug. Mol. Ther. 28, 1759–1771 (2020).

41. H. Li, T. Lee, T. Dziubla, F. Pi, S. Guo, J. Xu, C. Li, F. Haque, X.-J. Liang, P. Guo, RNA as a stable polymer to build controllable and defined nanostructures for material and biomedical applications. Nano Today. 10, 631–655 (2015).

42. A. Aditham, H. Shi, J. Guo, H. Zeng, Y. Zhou, S. D. Wade, J. Huang, J. Liu, X. Wang, Chemically Modified mocRNAs for Highly Efficient Protein Expression in Mammalian Cells. ACS Chem. Biol. 17, 3352–3366 (2022).

43. E. Sebastian Vik, M. Sameen Nawaz, P. Strøm Andersen, C. Fladeby, M. Bjørås, B. Dalhus, I. Alseth, Endonuclease V cleaves at inosines in RNA. Nat. Commun. 4, 1–7 (2013).

44. I. Alseth, B. Dalhus, M. Bjørås, Inosine in DNA and RNA. Curr. Opin. Genet. Dev. 26, 116–123 (2014).

45. S. A. Scaringe, RNA oligonucleotide synthesis via 5’-silyl-2’-orthoester chemistry. Methods. 23, 206–217 (2001).

46. S. Palluk, D. H. Arlow, T. de Rond, S. Barthel, J. S. Kang, R. Bector, H. M. Baghdassarian, A. N. Truong, P. W. Kim, A. K. Singh, N. J. Hillson, J. D. Keasling, De novo DNA synthesis using polymerase-nucleotide conjugates. Nat. Biotechnol. 36, 645–650 (2018).

47. E. R. Moody, R. Obexer, F. Nickl, R. Spiess, S. L. Lovelock, An enzyme cascade enables production of therapeutic oligonucleotides in a single operation. Science. 380, 1150–1154 (2023).

48. J. Kim, C. Hu, C. Moufawad El Achkar, L. E. Black, J. Douville, A. Larson, M. K. Pendergast, S. F. Goldkind, E. A. Lee, A. Kuniholm, A. Soucy, J. Vaze, N. R. Belur, K. Fredriksen, I. Stojkovska, A. Tsytsykova, M. Armant, R. L. DiDonato, J. Choi, L. Cornelissen, L. M. Pereira, E. F. Augustine, C. A. Genetti, K. Dies, B. Barton, L. Williams, B. D. Goodlett, B. L. Riley, A. Pasternak, E. R. Berry, K. A. Pflock, S. Chu, C. Reed, K. Tyndall, P. B. Agrawal, A. H. Beggs, P. E. Grant, D. K. Urion, R. O. Snyder, S. E. Waisbren, A. Poduri, P. J. Park, A. Patterson, A. Biffi, J. R. Mazzulli, O. Bodamer, C. B. Berde, T. W. Yu, Patient-Customized Oligonucleotide Therapy for a Rare Genetic Disease. N. Engl. J. Med. 381, 1644–1652 (2019).

49. R. M. Meganck, R. S. Baric, Developing therapeutic approaches for twenty-first-century emerging infectious viral diseases. Nat. Med. 27, 401–410 (2021).

50. A. R. Kore, M. Shanmugasundaram, A. Senthilvelan, B. Srinivasan, An improved protection-free one-pot chemical synthesis of 2’-deoxynucleoside-5’-triphosphates. Nucleosides Nucleotides Nucleic Acids. 31, 423–431 (2012).

