## Supplementary Information for "A Platform for Controlled Template-Independent Enzymatic Synthesis of RNA Oligonucleotides and Therapeutics"

#### Title:

### Materials & Methods

#### Poly(U) Polymerase Expression & Purification

The DNA sequences for the wild-type CID1 *S. pombe* Poly(U) polymerase (PUP) (SEQ1) and mutant variants (H336R (SEQ2) and H336R-N171A-T172S (SEQ3)) were codon optimized, ordered as a gBlocks® (IDT), and inserted into the pET-28-a-(+) expression vector (EMD Millipore) through Gibson assembly. T7 Express chemically competent *E. coli* (New England Biolabs) cells were transformed with the fully assembled plasmid as per manufacturer's instructions and positive transformants were selected for on LB-kanamycin plates. Bacterial colonies were sequenced and those with correct sequences were grown in liquid LB-kanamycin media overnight at 37 °C, diluted the next morning 1:400 in fresh liquid LB supplemented with 50 µg/mL kanamycin, and induced with 1 mM IPTG (Sigma) at OD600 = ~0.6. The induced liquid cultures were incubated overnight at 15 °C, shaking at 250 RPM. Cultures were then pelleted at 3500 x g for 10 minutes and His-Tag purified using a HisTalon Resin Kit as per manufacturer's instructions (Takara). The eluted enzyme samples were concentrated and buffer-exchanged into 1X PUP storage buffer (10 mM Tris-HCl, 250 mM NaCl, 1 mM DTT, 0.1 mM EDTA, pH 7.5 at 25 °C) using 30K MWCO 15-mL filter columns (Millipore) and then flash frozen using liquid nitrogen and stored at -80 °C until needed.

#### Endonuclease V Expression & Purification

Wild-type *E. coli* Endonuclease V (SEQ4) and Endonuclease V fused to a maltose binding protein (MBP) at the N-terminus (SEQ 5) were expressed and purified as similarly described for Poly(U) Polymerase with the exception of the 1X Endo V storage buffer being comprised of 10 mM Tris-HCl, 250 mM NaCl, 0.1 mM EDTA, and 1 mM DTT, pH 8.0 at 25 °C. Expressed enzyme was flash frozen using liquid nitrogen and stored at -80 °C until needed.

#### Standard Liquid Bulk Phase Reactions

##### *Controlled Enzymatic Extension Reactions with Poly(U) Polymerase*

A standard master mix for controlled oligonucleotide enzymatic extension in the bulk liquid phase was comprised of 1x Extension Buffer (50 mM NaCl, 10 mM Tris-HCl, 8 mM $MgCl_2$, 2 mM $MnCl_2$, 1 mM DTT at pH 7.9), 0.1 mg/mL enzyme, 1 mM 3'-O-allyl-ether reversible terminator NTP, and 2.5 pmol/µL initiator oligonucleotide. All extension reactions were carried out at 37 °C for 30 minutes unless otherwise specified. Following incubation, 2 µL Proteinase K (NEB) was added to the samples, mixed gently, and then incubated for 5 more minutes at 37 °C. Extension products were then isolated using Zymo Oligonucleotide Clean and Concentrator spin-columns following manufacturer's instructions and eluted in MilliQ water. All standard liquid bulk phase extension reactions used an internally Cy5-labeled, 19-nt initiator oligonucleotide with a 5'- amino C12 group (sequence: /5AmMC12/-rU-rU-rU-/iCy5/-rU-rU-rU-rU-rU-rU-rU-rU-rU-rU-rU-rU-rU-rU-rU-rU) (IDT) and PUP mutant variant H336R unless otherwise specified.

*Allyl Ether Deblocking Reactions*

A standard deblocking reaction was comprised of degassed 10 mM Tris-HCl (pH 6.7), 1.15 nmol/µL sodium tetrachloropalladate(II) ($Na_2PdCl_4$) (Sigma), 8.80 nmol/µL triphenylphosphine-3,3′,3″-trisulfonic acid trisodium salt ($P(PhSO_3Na)_3$) (Sigma) and 2.5 pmol/µL blocked RNA oligonucleotide. All deblocking reactions were carried out at 62 °C for 12 minutes unless otherwise specified. Deblocked oligonucleotide was then purified using Zymo Oligonucleotide Clean and Concentrator spin-columns and eluted in MilliQ water.

*Endonuclease V Mediated Oligonucleotide Cleavage Reactions*

A standard Endonuclease V mediated cleavage reaction in the liquid bulk phase was carried out by incubating 2.5 pmol/µL initiator oligonucleotide containing a deoxy- or riboinsoine base in 1x cleavage buffer (50 mM potassium acetate, 20 mM Tris-acetate, 10 mM magnesium acetate, 1 mM DTT, pH 7.9 at 25 °C) and 0.05 mg/mL Endonuclease V at 37 °C for 30 minutes. For commercially sourced Endonuclease V (NEB), 20U of enzyme was added to reactions. Cleaved oligonucleotide was purified Zymo Oligonucleotide Clean and Concentrator spin-columns and eluted in MilliQ water for downstream analysis. Wild-type Endonuclease V prepared in-house was used for all standard cleavage reactions unless otherwise noted.

*Phosphatase-Mediated Oligonucleotide Dephosphorylation Reactions*

A standard dephosphorylation reaction in the liquid bulk phase was carried out by incubating an 5'-phosphate modified oligonucleotide with 100 U of Antarctic Phosphatase (NEB) in 1x reaction buffer (50 mM Bis-Tris-Propane-HCl, 1 mM $MgCl_2$, and 0.1 mM $ZnCl_2$ pH 6 at 25 °C) at a concentration of 2.5 pmol/µL for 30 minutes at 37 °C. Dephosphorylated oligonucleotide was purified using Zymo Oligonucleotide Clean and Concentrator spin-columns and eluted in MilliQ water.

*Preparation of CPG Solid Support Derivatized with Initiator Oligonucleotide*

Initiator oligonucleotide labeled solid support was prepared by covalently attaching a 5'- amine modified oligonucleotide to the surface of long-chain alkylamine controlled pore glass (LCAA-CPG) using a Bis-N-hydroxysuccinimide ester linker. To a 20 mL scintillation vial, 2 g of dry LCAA-CPG with a pore size of 1000Å (ChemGenes) was added and washed 3x with 15 mL of anhydrous dimethylformamide (DMF) (Sigma). The vial containing LCAA-CPG was rotated for 15 minutes during each DMF wash and liquid waste was discarded. A 100 mg/mL solution of Bis-PEG5-NHS ester linker (BroadPharm) was prepared in anhydrous DMF. After the final wash of the LCAA-CPG, 5 mL of the 100 mg/mL Bis-PEG5-NHS ester linker solution was added. Additional anhydrous DMF was added (~4-5 mL) to bring the solution to volume and then the vial was incubated at room temperature for 2 hours while rotating. After incubation, the Bis-PEG5-NHS solution was discarded, and the LCAA-CPG was washed 3x with anhydrous DMF. To link the initiator oligonucleotide to the now derivatized LCAA-CPG, 500 µL of a 1 mM solution containing the 5'- amine modified oligonucleotide (sequence: 5'-$NH_2$-C12-rU-rC-rU-rA-rC-rC-rA-rU-rA-rU-rA-rU-dI-rA-rA-rC-rA-rA-rG-rC-rA-rC-rA-rCr-U-rA-rA-rA-rU-rU) (IDT), where dI is deoxyinosine) was prepared in MilliQ water and directly added to the LCAA-CPG in the vial along with an additional 10 mL of fresh anhydrous DMF. This solution was incubated at room temperature for at least 4 hours while rotating. After incubation, the initiator oligonucleotide labeled CPG solid support was washed 3x with anhydrous DMF, then a 0.1 M solution of succinimide anhydride (Sigma) to cap any remaining primary amine sites on the surface of the LCAA-CPG. The solid support was then transferred to a 20 mL solid-phase extraction (SPE) column/filter (Sigma) and washed in excess with a 10 mM Tris-HCl solution using a vacuum manifold. The resultant labeled CPG solid support was then stored at 4 °C until needed for enzymatic RNA oligonucleotide synthesis.

#### Standard Solid Phase Reactions

##### *Controlled Enzymatic Extension Reactions on CPG Solid Support*

A standard solid phase enzymatic extension reaction was conducted by incubating 150 mg of initiator oligonucleotide labeled CPG solid support with 1x Extension Buffer (50 mM NaCl, 10 mM Tris-HCl, 8 mM $MgCl_2$, 2 mM $MnCl_2$, 1 mM DTT at pH 7.9), 0.1 mg/mL enzyme, 1 mM allyl-ether reversible terminator NTP terminator in a total volume of 1.5 mL. Reactions were carried out in a "stir format" where the CPG solid support and extension reaction master mix was combined in a capped 3 mL solid phase extraction (SPE) column (Agilent) containing a small flea-sized magnetic stir bar and placed on a custom-made heat block/magnetic stir plate set to 37 °C and 1500 RPM, respectively, for 30 minutes. Following incubation, the SPE column was uncapped and placed on a vacuum manifold where the extension master mix was discarded. The solid support was then washed 2x with 3 mL of DNA wash buffer (Zymo) and 5x with 3 mL of 10 mM Tris-HCl (pH 6.7). During each wash, the CPG solid support was gently agitated with a 1 mL pipette to ensure complete washing. The SPE column was then removed from the vacuum manifold, capped, and placed on ice or stored at 4 °C until needed.

##### *Allyl Ether Deblocking Reactions on CPG Solid Support*

To remove the 3'-O-allyl ether blocking group from the growing oligonucleotide on the surface of the CPG solid support, 1 mL of deblocking solution (degassed, 10 mM Tris-HCl (pH 6.7), 1.15 nmol/µL $Na_2PdCl_4$ (Sigma), 8.80 nmol/µL $P(PhSO_3Na)_3$, Sigma) was prepared and added directly to SPE column. The SPE column was then placed on the combination heat block/magnetic stir plate and incubated at 62 °C for 12 minutes without stirring. After incubation, the SPE column was uncapped and placed on a vacuum manifold where the deblocking solution was immediately discarded. The solid support was then washed 1x with 3 mL of 3% ammonium hydroxide (Sigma), 2x with 3 mL of DNA wash buffer (Zymo) and 5x with 3 mL of a 10 mM Tris-HCl solution (pH 6.7). During each wash, the CPG solid support was gently agitated with a 1 mL pipette to ensure complete washing. The SPE column was then removed from the vacuum manifold, capped, and placed on ice until the subsequent enzymatic extension step or cleavage from the surface.

*Enzymatic Cleavage from the CPG Solid Support*

Once enzymatic RNA synthesis was completed, the oligonucleotide product was cleaved and collected by incubating 150 mg of CPG solid support with 1x cleavage buffer (50 mM potassium acetate, 20 mM Tris-acetate, 10 mM magnesium acetate, 1 mM DTT, pH 7.9 at 25 °C) and 0.05 mg/mL Endonuclease V at 37 °C for 30 minutes in a total volume of 0.750 mL. Cleavage reactions were carried out as before in a "stir format" (**Fig. S18**) where the same SPE column containing the CPG solid support and magnetic flea was incubated at 37 °C and spun at 1500 RPM for 30 minutes. After incubation, the cleaved oligonucleotide was collected by placing the uncapped SPE column into a 15 mL empty falcon tube and centrifuged for 1 minute at 1000 x g. The RNA oligonucleotide product was then stored at -20 °C until needed for analysis or downstream applications.

**Conjugation of GalNAc Ligand to Propargyl Functional Handles using Click Chemistry**

The following stock solutions were prepared prior to performing the click chemistry protocol: 5 mM ascorbic acid (Sigma) prepared in MilliQ water, 10 mM Copper (II)-TBTA in 55% DMSO (prepared by dissolving 25 mg copper (II) sulfate pentahydrate (Sigma) in 10 mL MilliQ water and mixing with a solution of 58 mg of Tris(benzyltriazolylmethyl)amine (TBTA) ligand (Sigma) in 11 mL of anhydrous DMSO), and 2M Triethylammonium Acetate (TEAA) Buffer, pH 7.0 (prepared by mix 2.78 mL of triethylamine (Chem-Impex) with 1.14 mL of glacial acetic acid (Fisher), bring to volume in 10 mL and adjusting the pH to 7.0.). A stock solution of α-GalNAc-PEG3-Azide ligand (Sigma) was prepared at a final concentration of 10 mM in 100% anhydrous DMSO. Click chemistry reactions took place in a 1.5 mL HPLC glass vial and the standard components are as follows: 200 mM TEAA buffer, 0.5 mM ascorbic acid, 0.5 mM Copper (II)-TBTA complex, 30 µM α-GalNAc-PEG3-Azide, 20 µM 3'-O-propargyl ether modified RNA oligonucleotide (previously dissolved in MilliQ water) in a total volume of 100 µL. A low flow of high purity argon was bubbled through the click reaction for 30 seconds and then the HPLC vial was sealed tightly. Reactions were carried out overnight for 12 hours at room temperature and the α-GalNAc-PEG3 labeled RNA oligonucleotides were then purified using Zymo Oligonucleotide Clean and Concentrator spin-columns and eluted in MilliQ water for downstream analysis.

**Analysis of RNA Oligonucleotide Product Mass, Purity, and Concentration**

Enzymatic RNA oligonucleotide synthesis product profiles were analyzed either by a combination of high-resolution gel electrophoresis, MALDI-TOF mass spectrometry or liquid chromatography paired with mass spectrometry (LC/MS). A NanoDrop spectrophotometer (Thermo) was used to determine the concentration of all oligonucleotide products based on absorbance at 260 nm.

*High-resolution Gel Electrophoresis*

For high-resolution gel electrophoresis, 15% TBE urea denaturing gels were loaded with approximately 10-100 pmol of oligonucleotide material and run for 90 minutes at 185V per manufacturer's instructions. If necessary, gels were then incubated with 1X GelStar nucleic acid stain for 10 minutes on an orbital shaker. Gels were imaged using a GE Typhoon Imager using the appropriate laser and filter settings (SYBR: 497 nm | 520 nm and Cy5 651 nm | 670 nm).

*MALDI-TOF Mass Spectrometry*

Oligonucleotide masses were analyzed using MALDI-TOF by mixing 0.5 µL of prepared MALDI matrix (50 mg/mL 3-Hydroxypicolinic acid (3-HPA) and 10 mg/mL ammonium citrate in solution of 50/50 MS-grade acetonitrile and water) with 0.75 µL purified oligonucleotide directly on a 384-spot polished steel target plate. Samples were dried under vacuum for 5 minutes before analysis on a Bruker Rapid-flex MALDI-TOF using flexControl software. Peak acquisition was performed in positive polarity mode using an in-source decay (ISD) with reflector engaged method. Analysis of acquired data was performed through the Bruker flexAnalysis software, where all peaks were transformed and smoothed using the built-in baseline subtraction feature.

*LC/MS Analysis*

The final mass and purity of oligonucleotide intermediates and final products were assessed with an Agilent 1200 series LC with diode array detection and XBridge Oligonucleotide BEH C18 Column (130Å, 2.5 µm, 4.6 mm x 50 mm) (Waters) using a reversed phase method (Mobile Phase A: 5:95 methanol:water, 400 mM 1,1,1,3,3,3-Hexafluoro-2-propanol (HFIP), 15 mM triethylamine (TEA), Mobile Phase B: 50:50 methanol:water, 400 mM HFIP, 15 mM TEA, Method: 56% isocratic over 60 minutes ). Any mass spectra were obtained by running an Agilent

6400 series single-quadrupole MS module in scanning negative mode. Deconvolution was performed using the Agilent Bioanalysis software package.

**Preparation of Nucleoside Triphosphate Building Blocks from Nucleoside Intermediates**

*Procurement and Preparation of Reaction Components*

Natural and modified 3'-O-allyl ether nucleoside intermediates were prepared in-house and purchased from ChemGenes Corporation. Installation of the alpha-phosphorothioate (α-PS) was outsourced to Jena Bioscience. Our method for the synthesis of nucleoside triphosphates was conducted following previously reported literature (*50*). Triphosphorylation reactions were carried out in dried glassware, under an argon atmosphere, using anhydrous acetonitrile (Sigma) and tributyl amine (Sigma). Additionally, for triphosphorylation reactions, nucleoside and proton sponge were premixed in their reaction flasks and vacuum dried overnight. Nucleosides that would not dissolve readily were gently heated until they were almost completely dispersed in the solution.

*Isolation and Purification of Prepared Nucleoside Triphosphates*

Isolation and purification of nucleoside triphosphates was optimized and generally carried out as follows. Crude, quenched reaction mixture was washed with DCM. The aqueous layer containing triphosphorylated product was washed with hexane and concentrated *in vacuo*. The isolated material was purified via ion-exchange chromatography (DEAE Sepharose resin, Mobile Phase A: MilliQ water, Mobile Phase B: 1M triethyl ammonium bicarbonate buffer, pH 8 ± 0.5). The fractions containing primarily triphosphate (assayed via LC-MS) were combined and concentrated *in vacuo*. The sample was then further purified via preparatory HPLC (1260 Infinity Preparative LC System and a Phenomenex Jupiter C18 reverse-phase column, 10 µ particle size, 300 Å pore size, 250 mm length x 21.2 mm diameter). Generally, a single method provided excellent purities for the various NTP products (Mobile Phase A: 0.1% ammonium acetate in ACN, Mobile Phase B: 0.1% ammonium acetate in 1:667 water:ACN, general method: 5% isocratic over 20 minutes, then to 90% over 30 minutes). Pure fractions were combined, frozen, and lyophilized. The counter-ions on the triphosphate were exchanged by diluting lyophilic in triethyl ammonium bicarbonate (1M) and concentrating on the lyophilizer. Excess triethyl ammonium bicarbonate was removed from the sample by additional dilution with water and freezing, followed by lyophilization until the sample reached constant mass. *NOTE: For this protocol, yields and stock solutions of the nucleoside triphosphates were prepared with the presumption that all final products existed as tetra-triethylammonium salts. However, after additional, scrupulous lyophilization of analytical samples for the NMR analysis, we generally saw two to three triethylammoniums present in the final product.*

*Evaluation and of Prepared Nucleoside Triphosphates*

Analytical high performance liquid chromatography (HPLC) was performed on an Agilent 1260 series LC with diode array detection using a reversed phase method (Mobile Phase A: water, 400 mM HFIP, 15 mM TEA, Mobile Phase B: methanol, 400 mM HFIP, 15 mM TEA, unless otherwise mentioned). The best triphosphate resolution was obtained using a Waters™ XBridge Oligonucleotide BEH C18 Column (130Å, 2.5 µm, 4.6 mm x 50 mm). High resolution mass spectra were obtained via ESI-MS-HiRes on a Thermo q-Exactive Plus spectrometer. $^{1}H$ NMR (400 MHz on a Varian Mercury instrument) and $^{31}P$ NMR (400 MHz on a Varian Mercury instrument) spectra were measured. Chemical shifts are reported relative to the central line of residual solvent.

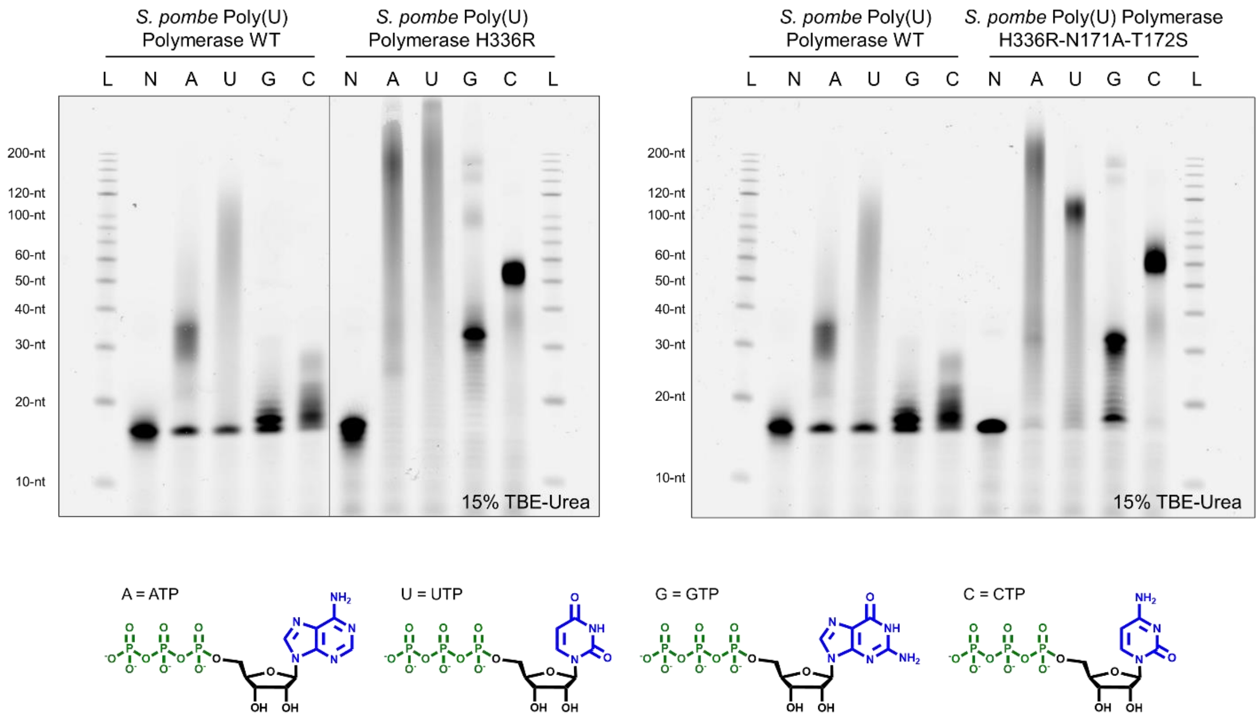


**Fig. S1:** A comparison of uncontrolled polymerization catalyzed by wild-type *S. pombe* Poly(U) Polymerase and two PUP mutant variants (H336R & H336R-N171A-T172S) using the natural unblocked RNA triphosphates (A, U, G, C) under standard extension reaction conditions. Control reactions (N) contained all components except for nucleotide. A 200-nt ssDNA ladder was used to measure the approximate length of uncontrolled polymerization.

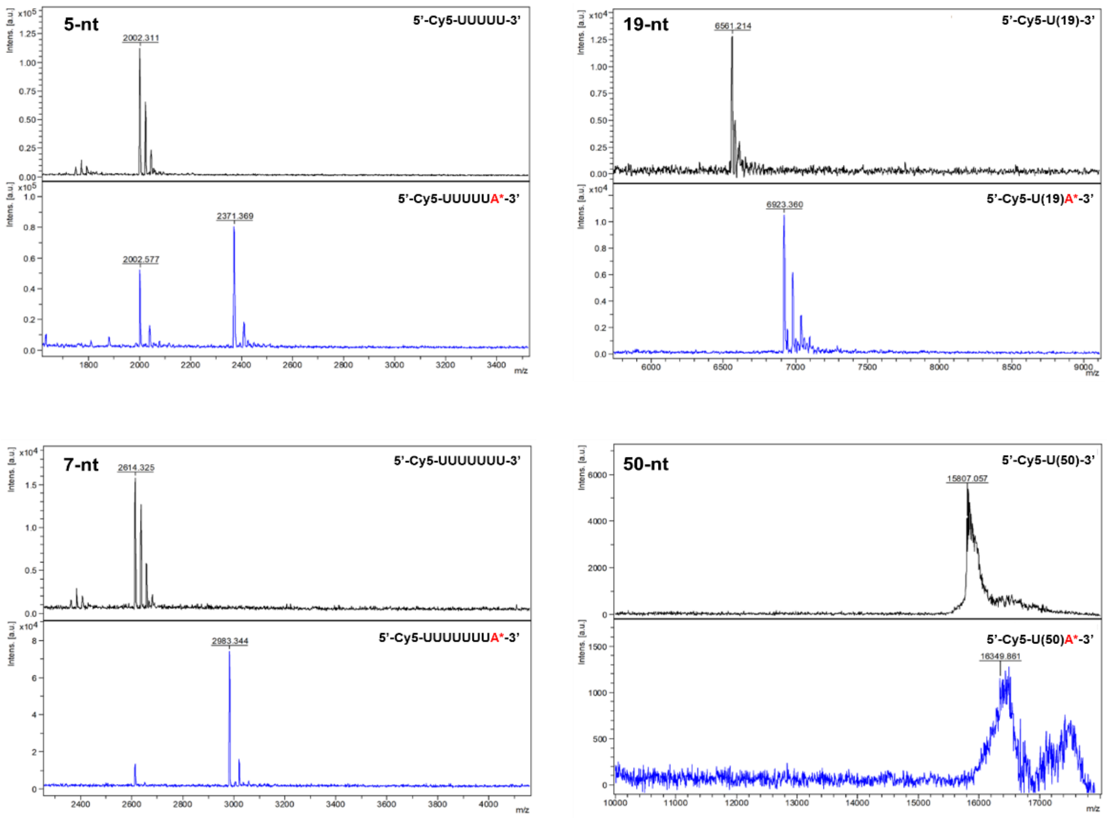


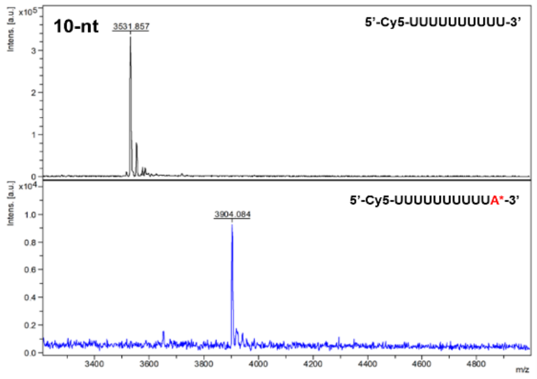


**Observed MALDI-TOF Masses**

| Init. Length | N+0 (m/z) | N+A* (m/z) |
|---|---|---|
| 5-nt | 2002.31 | 2371.36* |
| 7-nt | 2614.32 | 2983.34 |
| 10-nt | 3531.85 | 3904.08 |
| 19-nt | 6561.21 | 6823.36 |
| 50-nt | 15807.05 | 16349.86 |

* Indicates partial conversion to the final product

**Fig. S2A:** An evaluation of Poly(U) Polymerase mutant variant H336R extension activity under standard reaction conditions with a 3'-O-allyl ether ATP reversible terminator in the presence of initiator oligonucleotides of variable length (5-, 7-, 10-, 19-, & 50-nt) using MALDI-TOF mass spectrometry. All initiators were comprised of a poly-U RNA sequence and feature a 5'- Cy5 modification. For each initiator oligonucleotide, mass spectra are shown before (top, black) and after (bottom, blue) enzymatic extension.

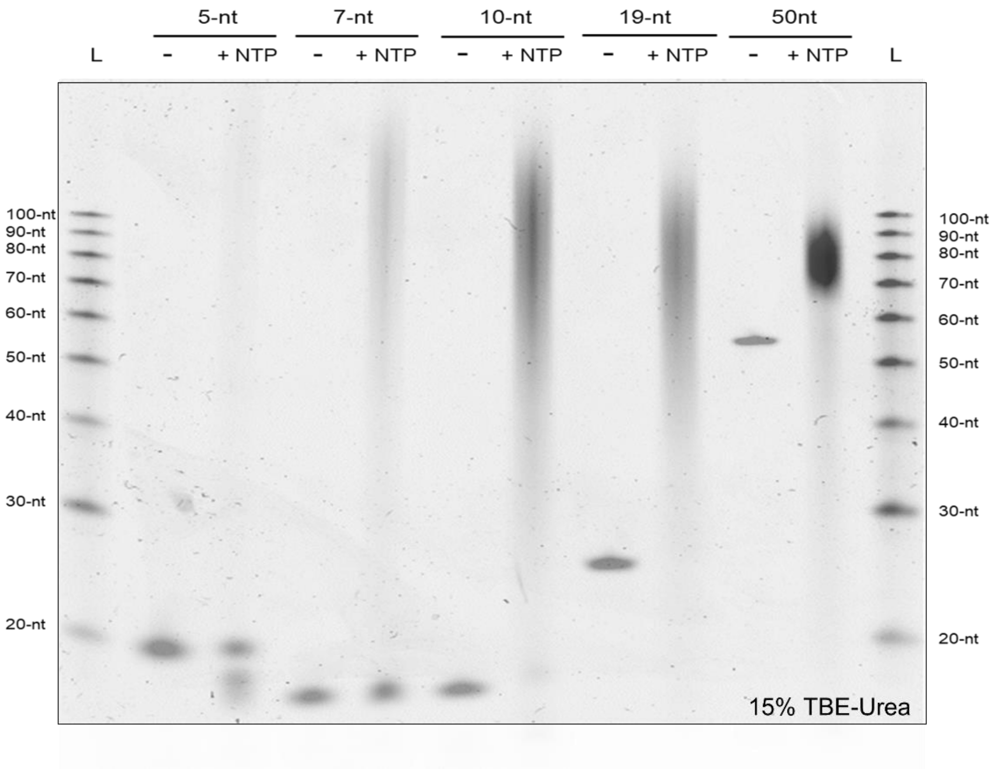


**Fig. S2B**: An evaluation of Poly(U) Polymerase mutant variant H336R extension activity under standard reaction conditions with an equimolar mixture of A, U, G, C natural NTPs in the presence of initiator oligonucleotide of variable length (5-, 7-, 10-, 19-, & 50-nt) using denaturing gel electrophoresis. All initiators were comprised of a poly-U RNA sequence and feature a 5’- Cy5 modification. For each initiator oligonucleotide, two lanes are shown on the gel: the negative (-) contained all reaction components except NTPs while the positive (+) were run with NTPs at a concentration of 1 mM (0.25 mM each base). A 100-nt ssDNA ladder was used to measure the approximate length of uncontrolled polymerization.

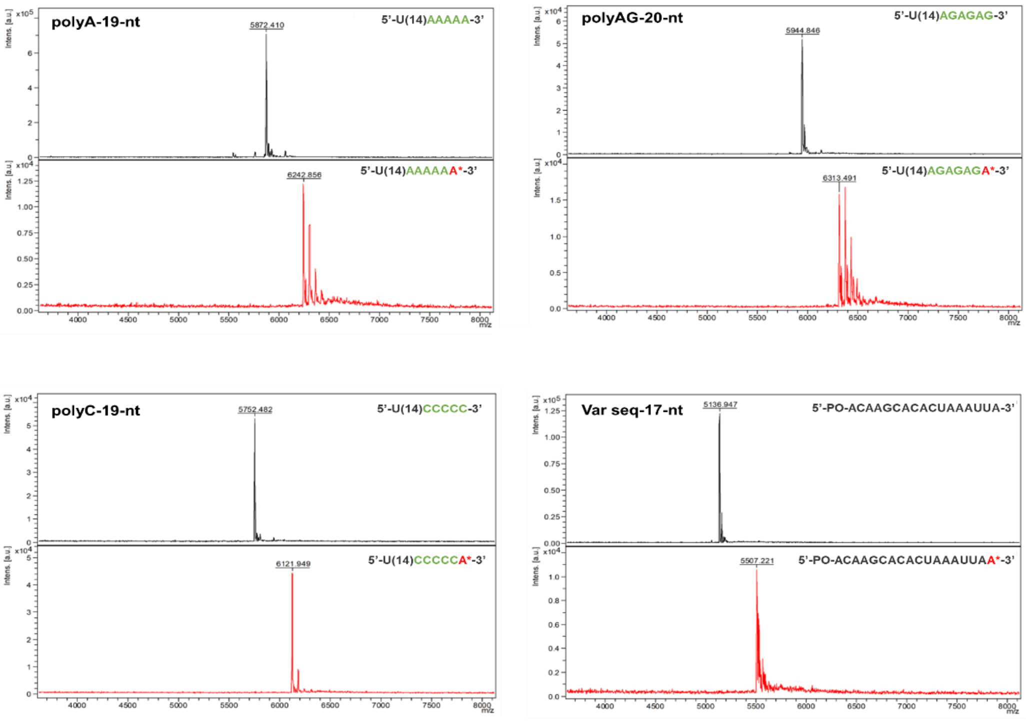


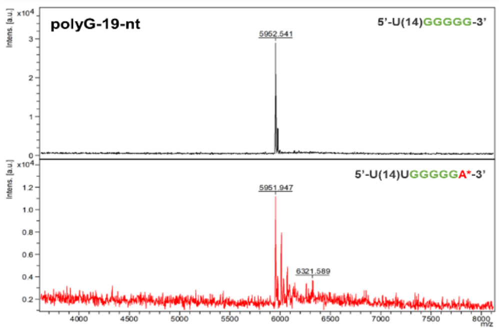


| Observed MALDI-TOF Masses | | |
|---|---|---|
| Init. Comp | N+0 (m/z) | N+A* (m/z) |
| Poly-A | 5872.41 | 6242.85 |
| Poly-C | 5752.48 | 6121.94 |
| Poly-G | 5952.54 | 6321.58* |
| Poly-AG | 5944.84 | 6313.49 |
| Variable Seq | 5136.94 | 5507.22 |

* Indicates partial conversion to the final product

**Fig. S3A**: An evaluation of Poly(U) Polymerase mutant variant H336R extension activity under standard reaction conditions with a 3'-O- allyl ether ATP reversible terminator in the presence of initiator oligonucleotides of variable sequence composition using MALDI-TOF mass spectrometry. Initiators were rationally constructed to ascertain a measurement of enzymatic tolerance for different compositions and combinations of base sequences on the 3'- terminus of the oligonucleotide. The initiators tested featured a stretch of poly-A, -U, -G, -AG RNA bases in addition to an initiator of variable base composition with a 5'-phosphate (5'-PO). The initiators were similar lengths, ranging from 17-nt to 20-nt. For each initiator oligonucleotide, mass spectra are shown before (top, black) and after (bottom, red) enzymatic extension

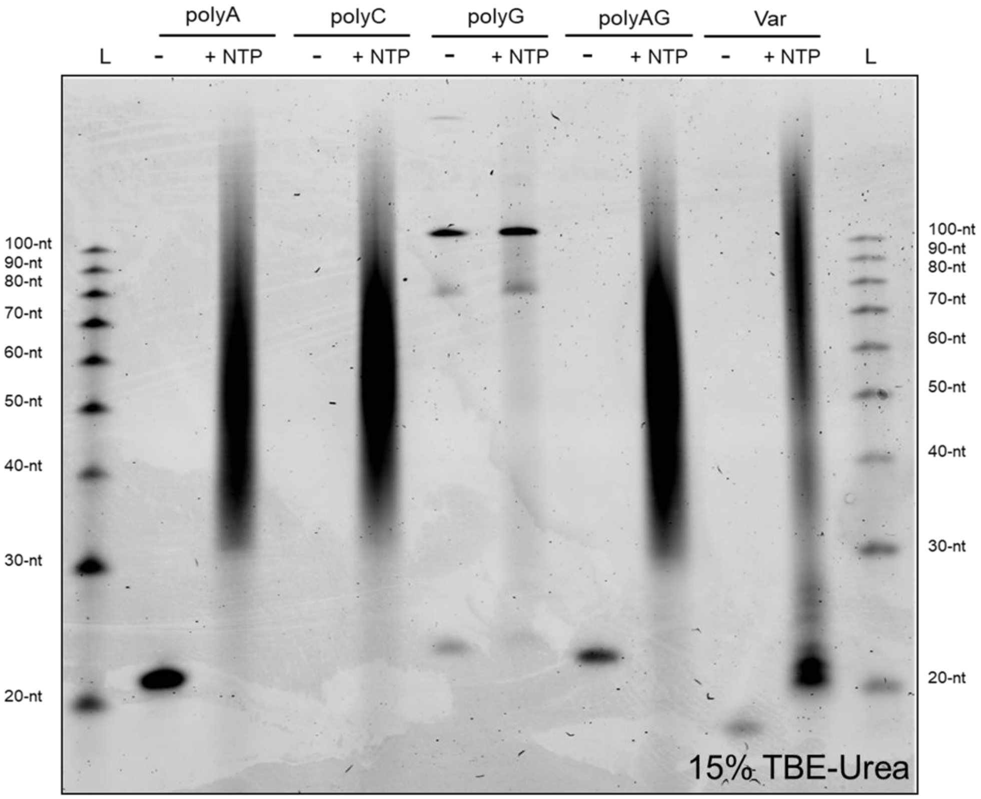


**Fig. S3B**: An evaluation of Poly(U) Polymerase mutant variant H336R extension activity under standard reaction conditions with natural NTPs in the presence of initiator oligonucleotide of variable sequence composition using denaturing gel electrophoresis. The initiators tested featured a stretch of poly-A, -U, -G, -AG RNA bases in addition to an initiator of random sequence with a 5’-PO modification. For each initiator oligonucleotide, two lanes are shown on the gel: the negative (-) contained all reaction components except NTPs while the positive (+) were run with NTPs at a concentration of 1 mM (0.25 mM each base). A 100-nt ssDNA ladder was used to measure the approximate length of uncontrolled polymerization.

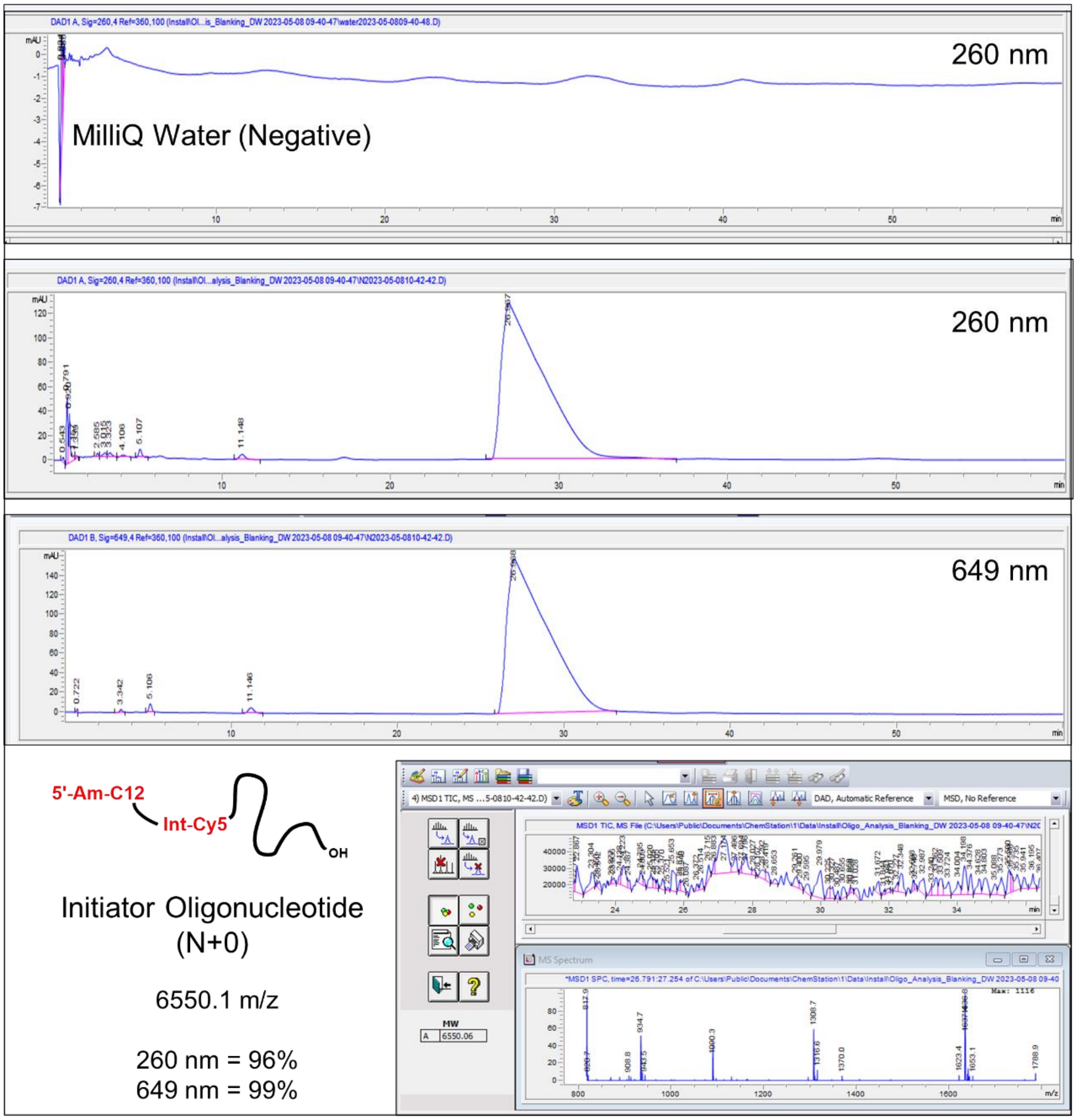


**Fig. S4A**: LC/MS analysis of initiator oligonucleotide starting material used to assess the coupling efficiency and isolated crude purity of controlled extension reactions with the PUP mutant and 3'-O- allyl ether NTP reversible terminator building blocks. The initiator oligonucleotide (N+0) is 19-nt in length and is comprised of a poly-U RNA sequence. The 5'- terminus features a primary amine and C12 linker as well as an internal Cy5 dye. Three HPLC chromatograms are shown: the first is a negative injection of MilliQ water, which all oligonucleotide samples are dissolved in, the second and third are an injection of approximately 2 nmol initiator with the DAD set to monitor 260 nm and 649 nm, respectively. The purity of the initiator was assessed at these wavelengths and found to be 260 nm = 96% and 649 nm = 99%, assuming similar response factors for the impurities as the oligonucleotide. The oligonucleotide mass was determined by deconvolution to be 6550.01 m/z, which corresponds to the expected mass.

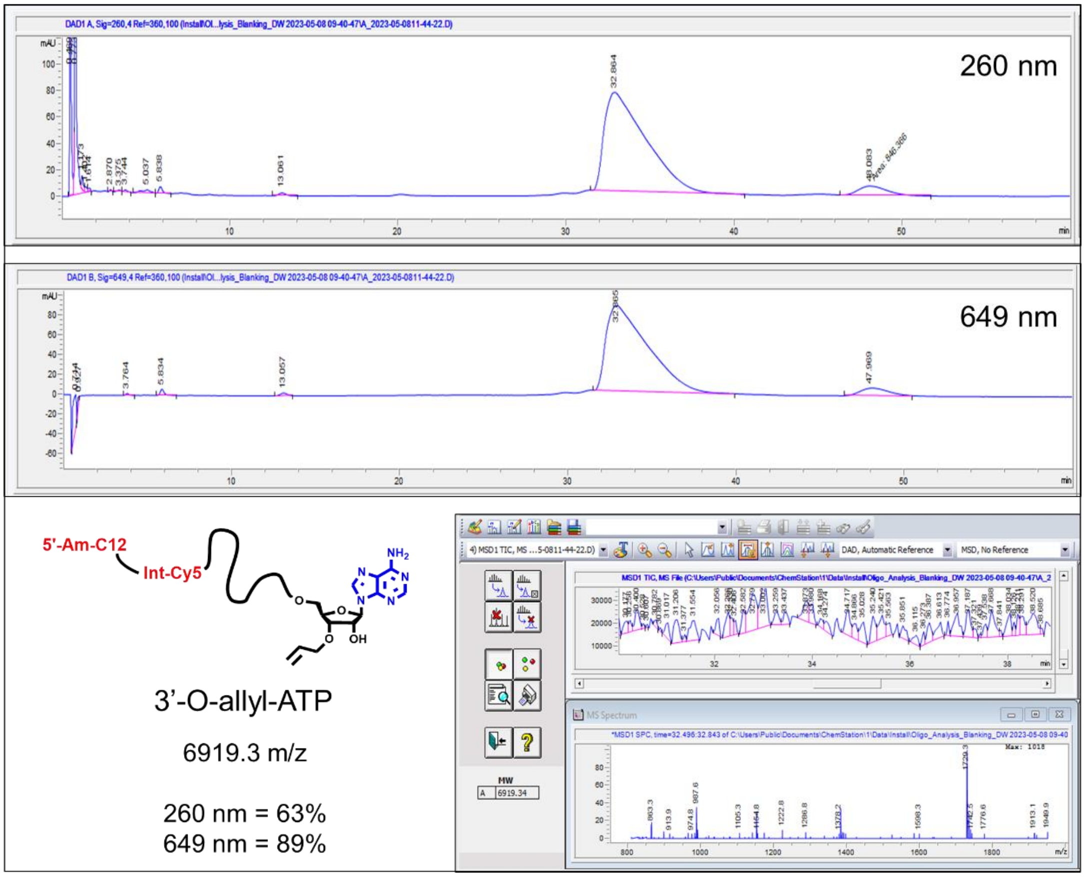


**Fig. S4B:** LC/MS analysis of controlled enzymatic extension using a **3'-O-allyl ether ATP** reversible terminator with the PUP mutant H336R, standard reaction conditions, and Cy5 19-nt initiator. Two HPLC chromatograms are shown that correspond to a single 2 nmol injection of the **N+1*** product at 260 nm and 649 nm. The isolated crude purity was found to be 260 nm = 63% and 649 nm = 89%. The oligonucleotide mass was determined by deconvolution to be 6919.3 m/z, which corresponds to the expected mass of the **N+A*** product. The second peak at 48 min likely corresponds to an **N+AA*** impurity, which is about 10% of the total mass injected.

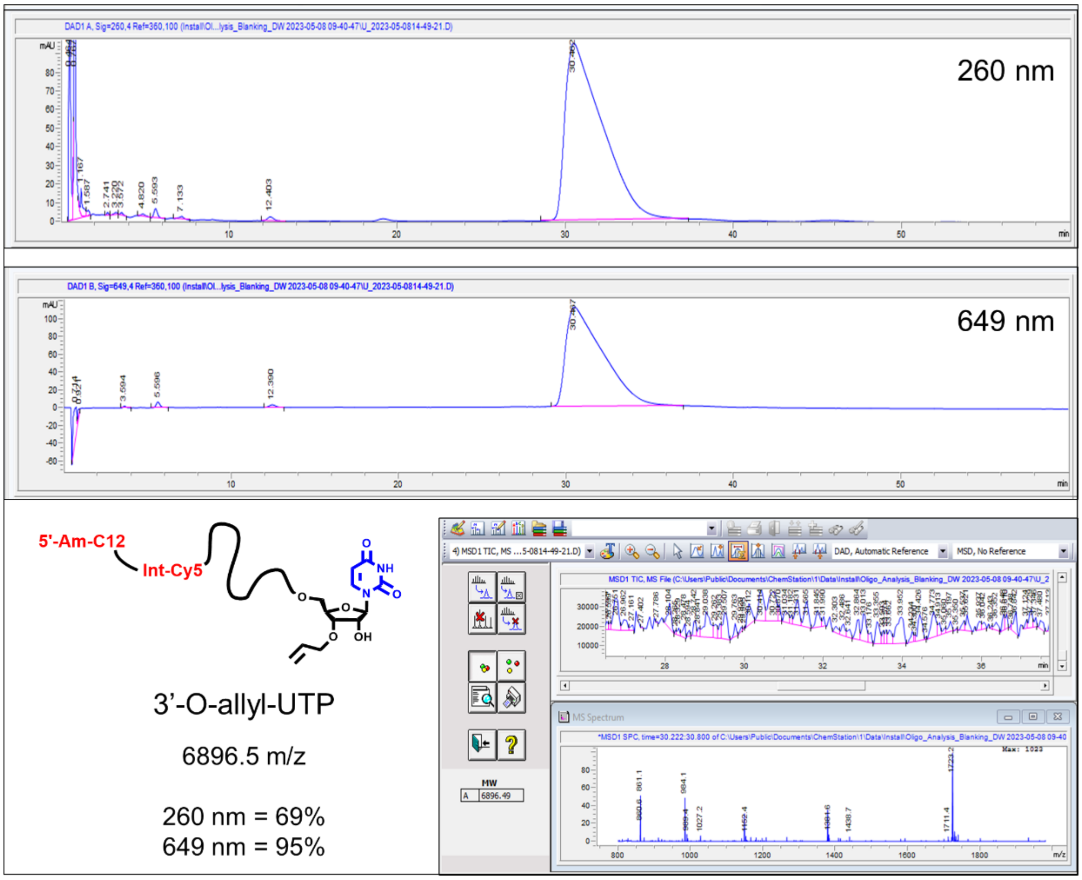


**Fig. S4C:** LC/MS analysis of controlled enzymatic extension using a **3'-O-allyl ether UTP** reversible terminator with the PUP H336R mutant, standard reaction conditions, and Cy5 19-nt initiator. Two HPLC chromatograms are shown that correspond to a single 2 nmol injection of the **N+1*** product at 260 nm and 649 nm. The integration of the main peak was found to be 260 nm = 69% and 649 nm = 95%. The oligonucleotide mass was determined by deconvolution to be 6896.5 m/z, which corresponds to the expected mass of the **N+U*** product.

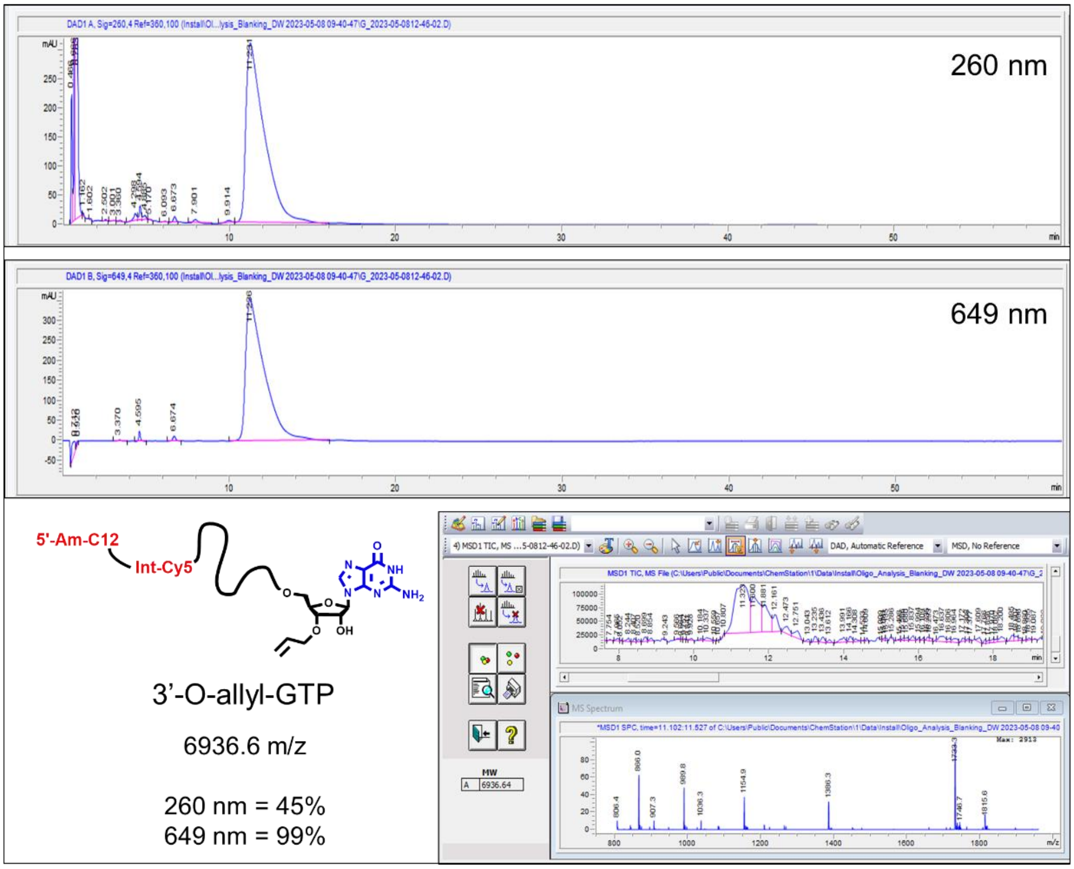


**Fig. S4D:** LC/MS analysis of controlled enzymatic extension using a **3'-O-allyl ether GTP** reversible terminator with the PUP mutant H336R, standard reaction conditions, and Cy5 19-nt initiator. Two HPLC chromatograms are shown that correspond to a single 2 nmol injection of the **N+1*** product at 260 nm and 649 nm. The integration of the main peak was found to be 260 nm = 45% and 649 nm = 99%. The oligonucleotide mass was determined by deconvolution to be 6896.5 m/z, which corresponds to the expected mass of the **N+G*** product.

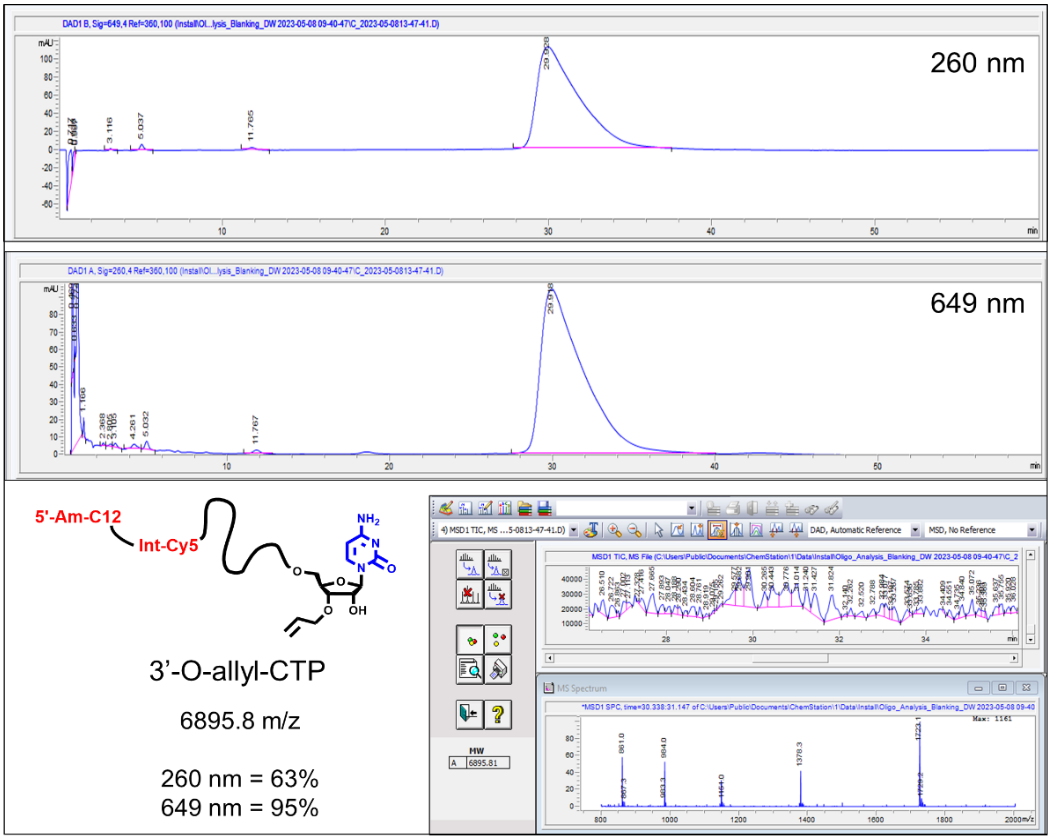


**Fig. S4E:** LC/MS analysis of controlled enzymatic extension using a **3'-O-allyl ether CTP** reversible terminator with the PUP mutant H336R, standard reaction conditions, and Cy5 19-nt initiator. Two HPLC chromatograms are shown that correspond to a single ~2 nmol injection of the **N+1*** product at 260 nm and 649 nm. The integration of the main peak was found to be 260 nm = 63% and 649 nm = 95%. The oligonucleotide mass was determined by deconvolution to be 6895.8 m/z, which corresponds to the expected mass of the **N+C*** product.

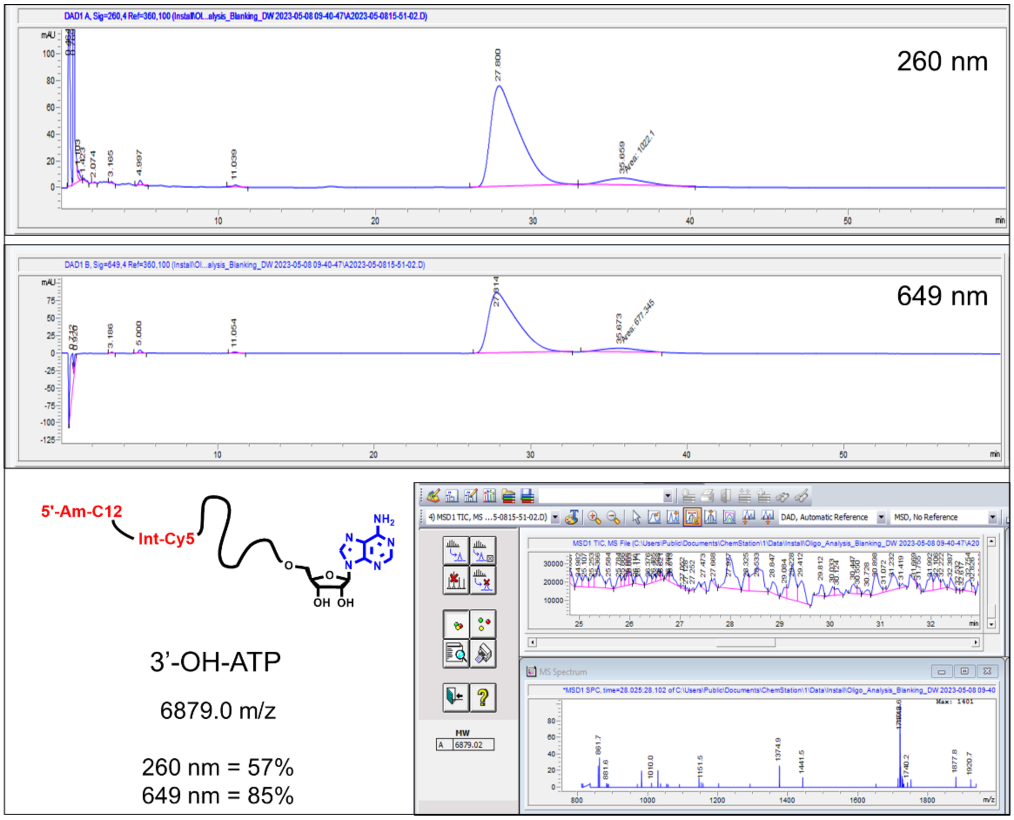


**Fig. S5A:** LC/MS analysis of 3'-O-allyl ether ATP deblocking post-enzymatic extension. Two HPLC chromatograms are shown that correspond to a single 2 nmol injection of the **N+1** product at 260 nm and 649 nm. The integration of the main peak was found to be 260 nm = 57% and 649 nm = 85%. The oligonucleotide mass was determined by deconvolution to be 6879.0 m/z, which corresponds to the expected mass of the **N+A** product. The second peak at 36 min likely corresponds to an **N+AA** impurity, which is about 15% of the total mass injected.

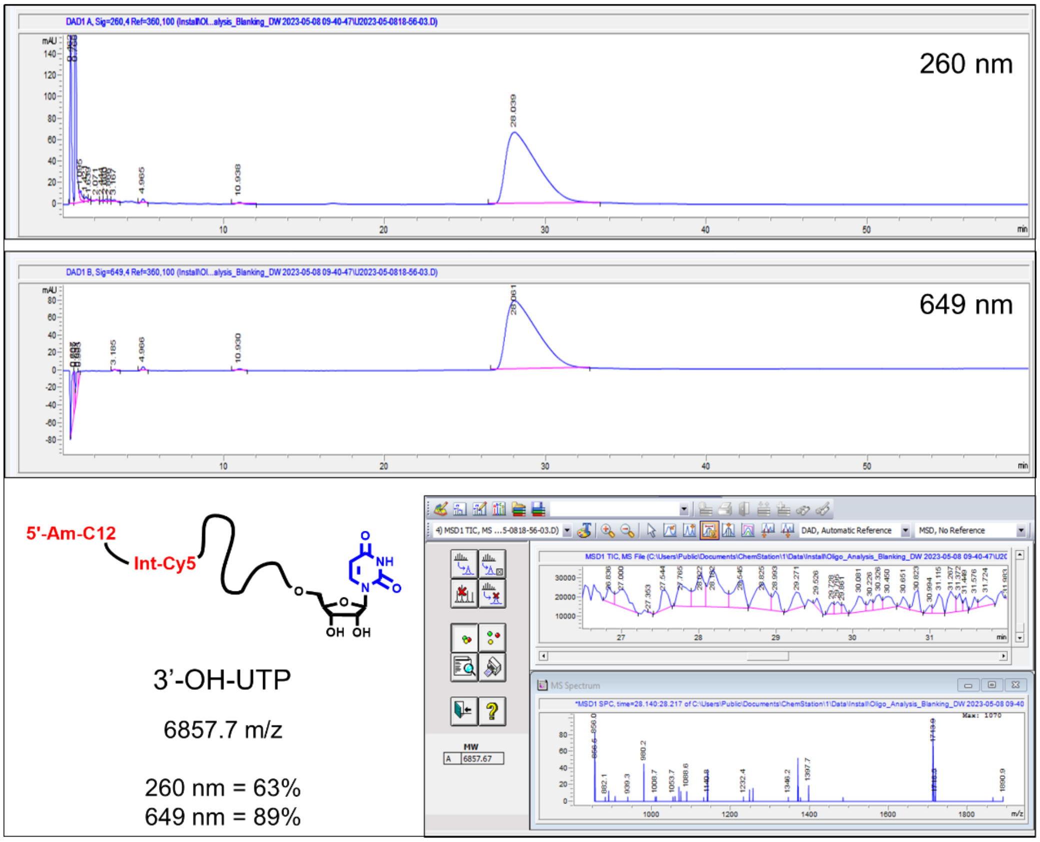


**Fig. S5B:** LC/MS analysis of 3'-O-allyl ether UTP deblocking post-enzymatic extension. Two HPLC chromatograms are shown that correspond to a single 2 nmol injection of the **N+1** product at 260 nm and 649 nm. The integration of the main peak was found to be 260 nm = 63% and 649 nm = 89%. The oligonucleotide mass was determined by deconvolution to be 6857.7 m/z, which corresponds to the expected mass of the **N+U** product.

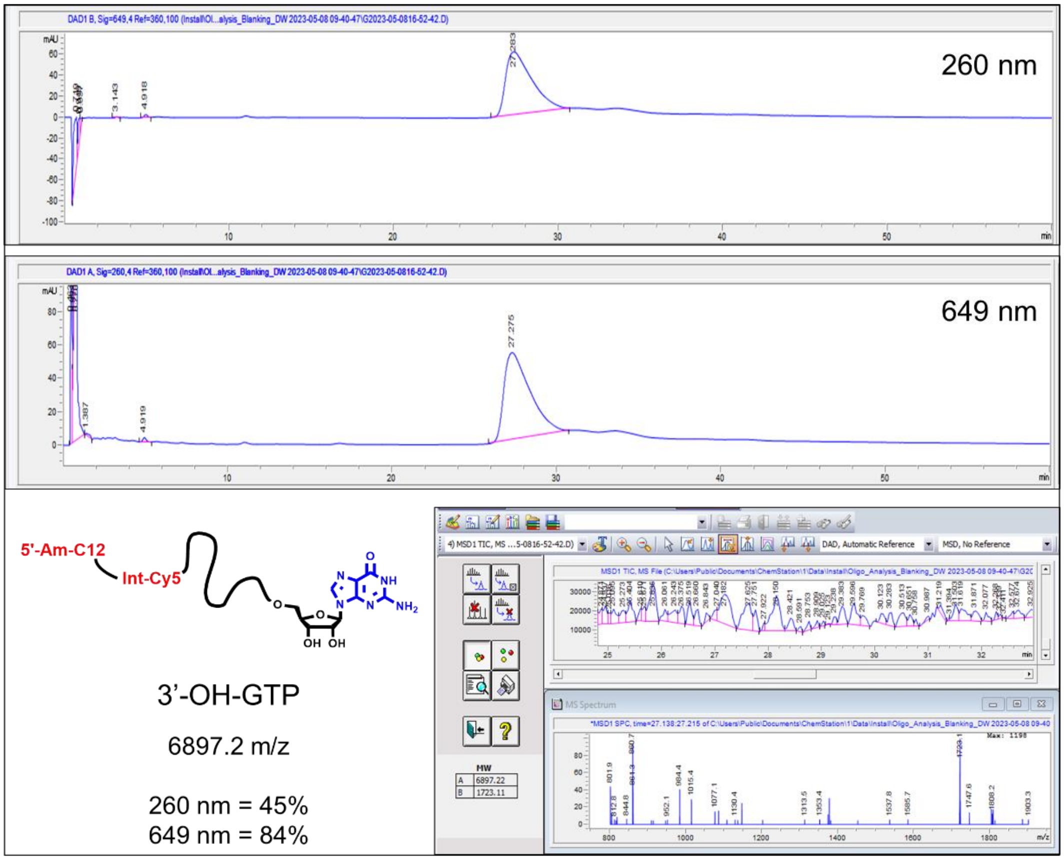


**Fig. S5C:** LC/MS analysis of 3'-O-allyl ether GTP deblocking post-enzymatic extension. Two HPLC chromatograms are shown that correspond to a single 2 nmol injection of the **N+1** product at 260 nm and 649 nm. The integration of the main peak was found to be 260 nm = 45% and 649 nm = 84%. The oligonucleotide mass was determined by deconvolution to be 6897.2 m/z, which corresponds to the expected mass of the **N+G** product.

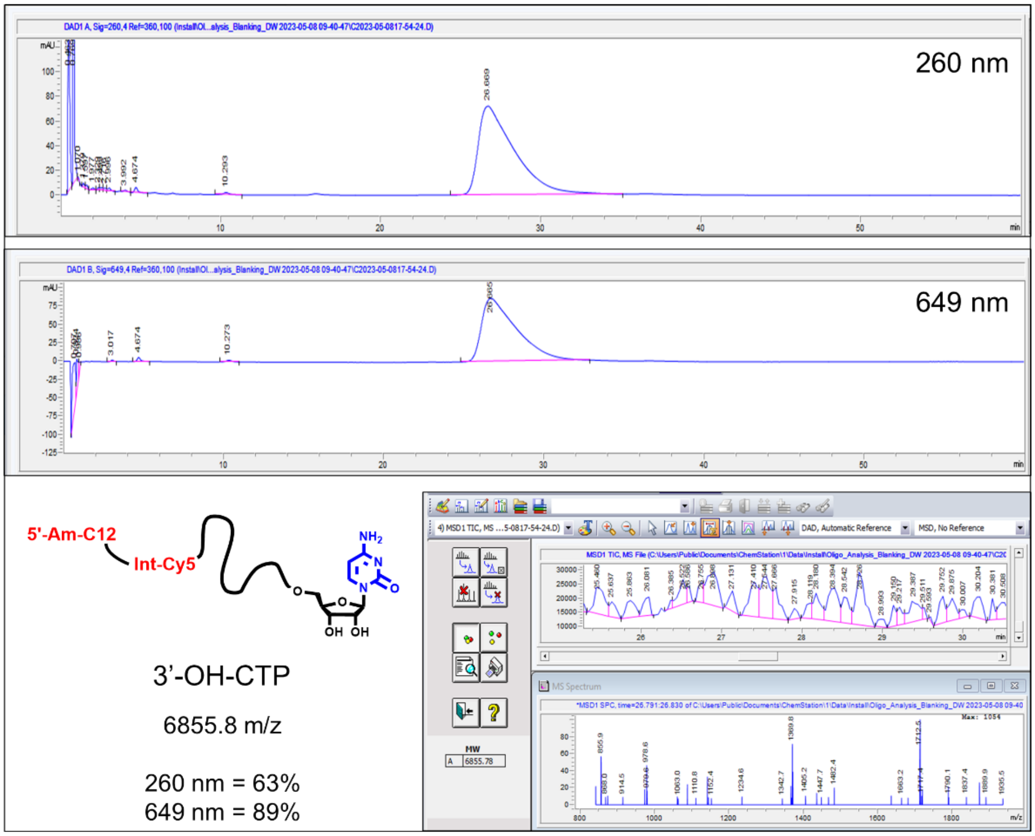


**Fig. S5D:** LC/MS analysis of 3'-O-allyl ether CTP deblocking post-enzymatic extension. Two HPLC chromatograms are shown that correspond to a single ~2 nmol injection of the **N+1** product at 260 nm and 649 nm. The integration of the main peak was found to be 260 nm = 63% and 649 nm = 89%. The oligonucleotide mass was determined by deconvolution to be 6855.8 m/z, which corresponds to the expected mass of the **N+C** product.

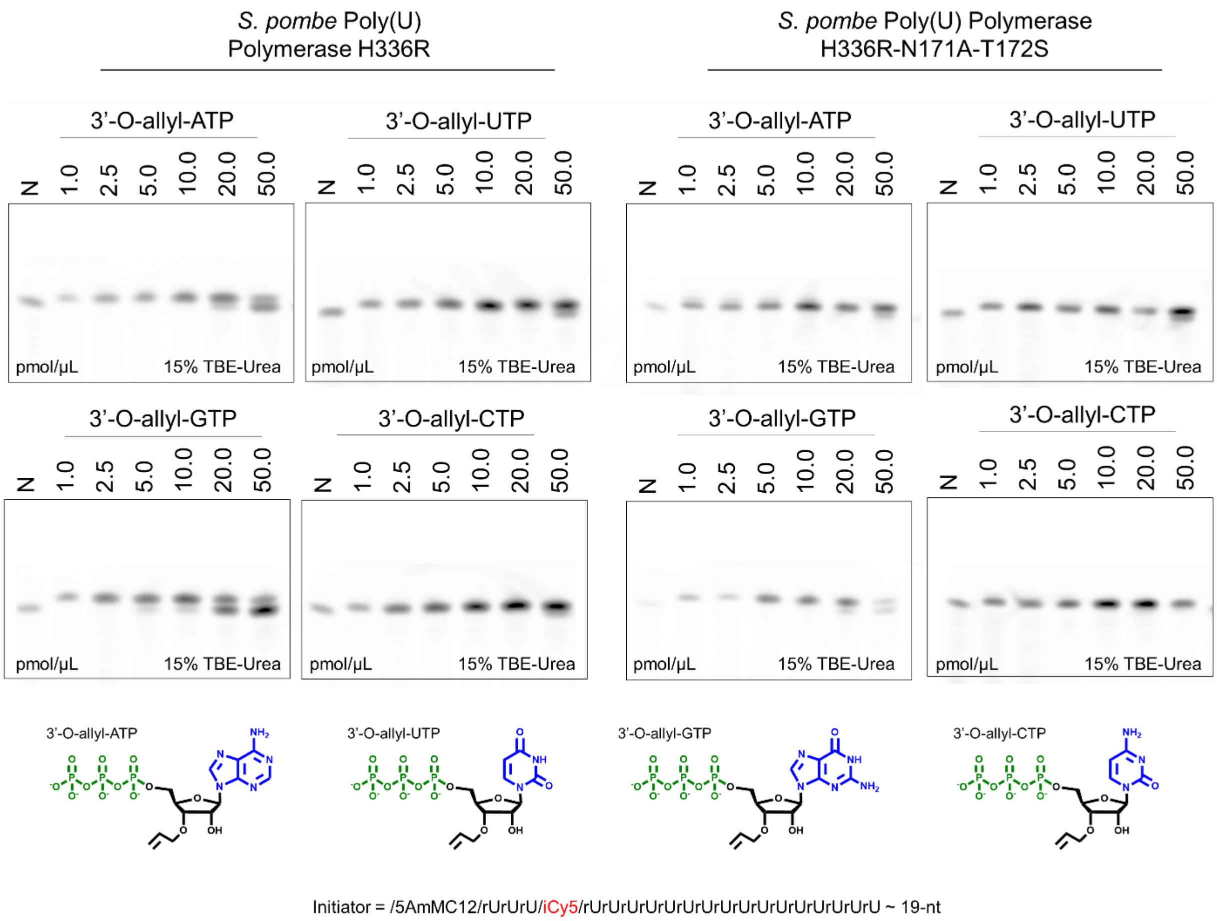


**Fig. S6**: An evaluation of enzymatic processing of the 3'-O-allyl ether NTP set (A, U, G, C) as a function of total initiator oligonucleotide input with Poly(U) Polymerase mutant variants H336R and H336R-N171A-T172S using high resolution gel electrophoresis. Reactions were carried out using standard conditions, a fixed volume of 10 µL and a fixed incubation time of 30 minutes. The concentration of initiator oligonucleotide ranged from 1.0 pmol/µL to 50.0 pmol/µL (a total input of 10 pmol to 500 pmol). Post incubation with the enzyme mutant variants, the efficiency of reversible terminator NTP incorporation was determined using denaturing gel electrophoresis. Control reactions (N) included all components except the NTP.

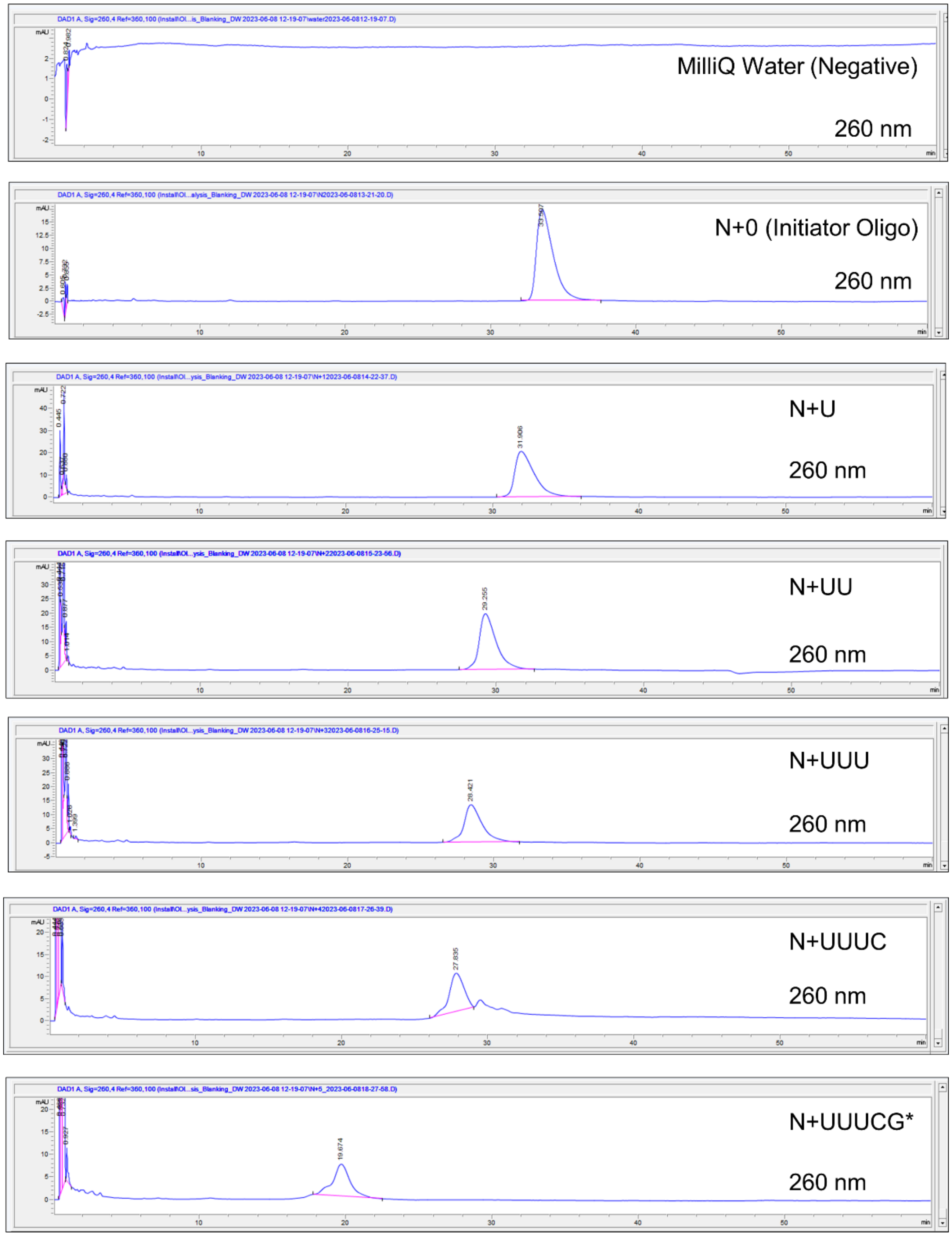


**Fig. S7A:** LC analysis of synthesis intermediates and final product generated when performing the controlled enzymatic synthesis of an **N+5*** RNA oligonucleotide with the sequence N+U-U-U-C-G*. Approximately ≤0.5 nmol of oligonucleotide intermediate and final product was injected onto the HPLC system. The isolated crude purity for each sample was determined by monitoring 260 nm and integrating over the chromatogram. A negative control consisting of MilliQ water was also included, which all oligonucleotide samples were dissolved in.

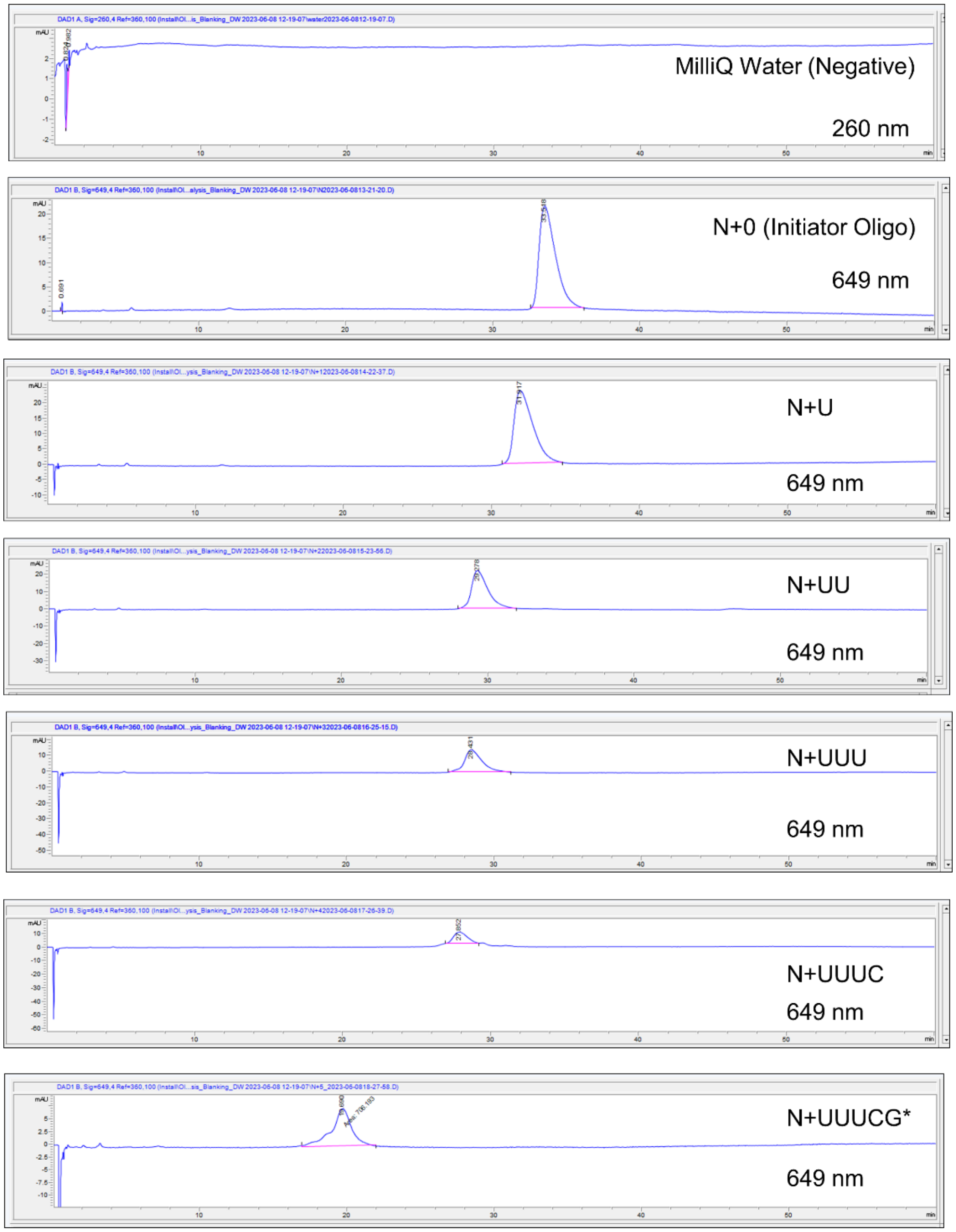


**Fig. S7B:** LC analysis of cycle intermediates and final product generated when performing the controlled enzymatic synthesis of an **N+5*** oligonucleotide with the natural 2'-OH RNA sequence N+U-U-U-C-G*, where N is a 19-nt Cy5 labeled initiator oligonucleotide. Approximately 500 pmol of oligonucleotide intermediate and final product was injected onto the HPLC system. The isolated crude purity for each sample was determined by monitoring 649 nm and integrating over the chromatogram. A negative control consisting of MilliQ water was also included, which all oligonucleotide samples were dissolved in.

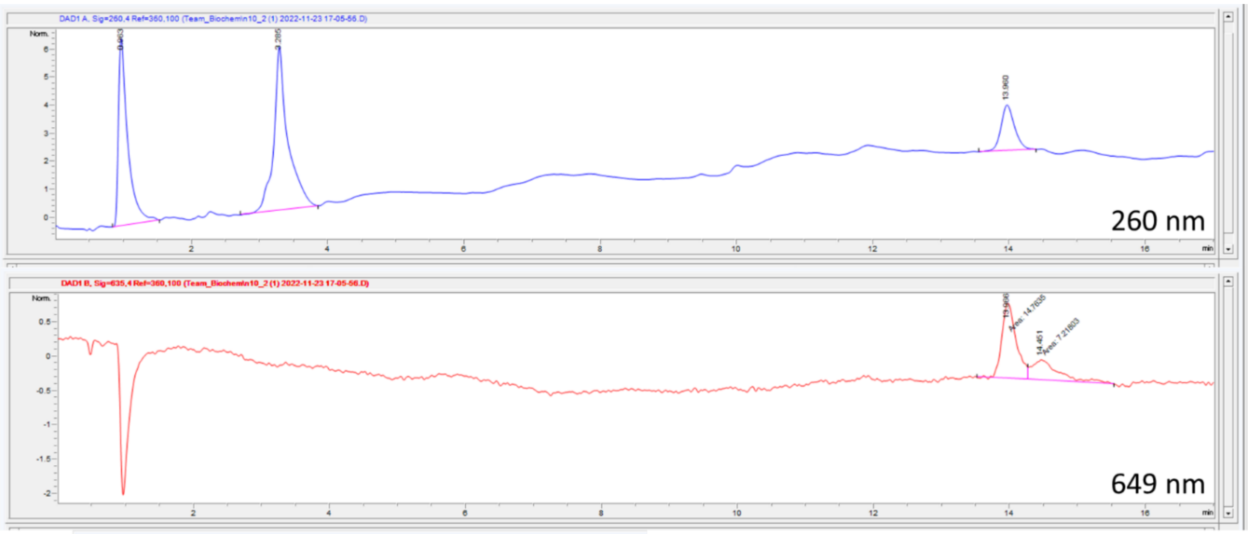


| # | Time | Type | Area | Height | Width | Area% | Symmetry |
|---|---|---|---|---|---|---|---|
| 1 | 0.963 | BB | 57.6 | 6 | 0.1331 | 37.154 | 0.368 |
| 2 | 3.285 | BB | 76.9 | 5.3 | 0.194 | 49.624 | 0.641 |
| 3 | 13.96 | BB | 20.5 | 1.5 | 0.2065 | 13.222 | 0.781 |

260 nm

| # | Time | Type | Area | Height | Width | Area% | Symmetry |
|---|---|---|---|---|---|---|---|
| 1 | 13.966 | MF | 14.8 | 9.9E-1 | 0.2494 | 67.163 | 0.7 |
| 2 | 14.451 | FM | 7.2 | 2.7E-1 | 0.4509 | 32.837 | 0.454 |

649 nm

**Fig. S8.** LC analysis of final product generated by controlled enzymatic synthesis of an **N+10*** oligonucleotide with the natural 2’-OH RNA sequence N+A-C-A-C-C-U-U-A-A-C*, where N is a 19-nt Cy5 labeled initiator oligonucleotide. Approximately 50 pmol of oligonucleotide final product was injected onto the HPLC system. The isolated crude purity (67.1%) was determined by monitoring 649 nm and integrating over the chromatogram. The apparent purity of the product as determined by 260 nm is 13.2%.

.

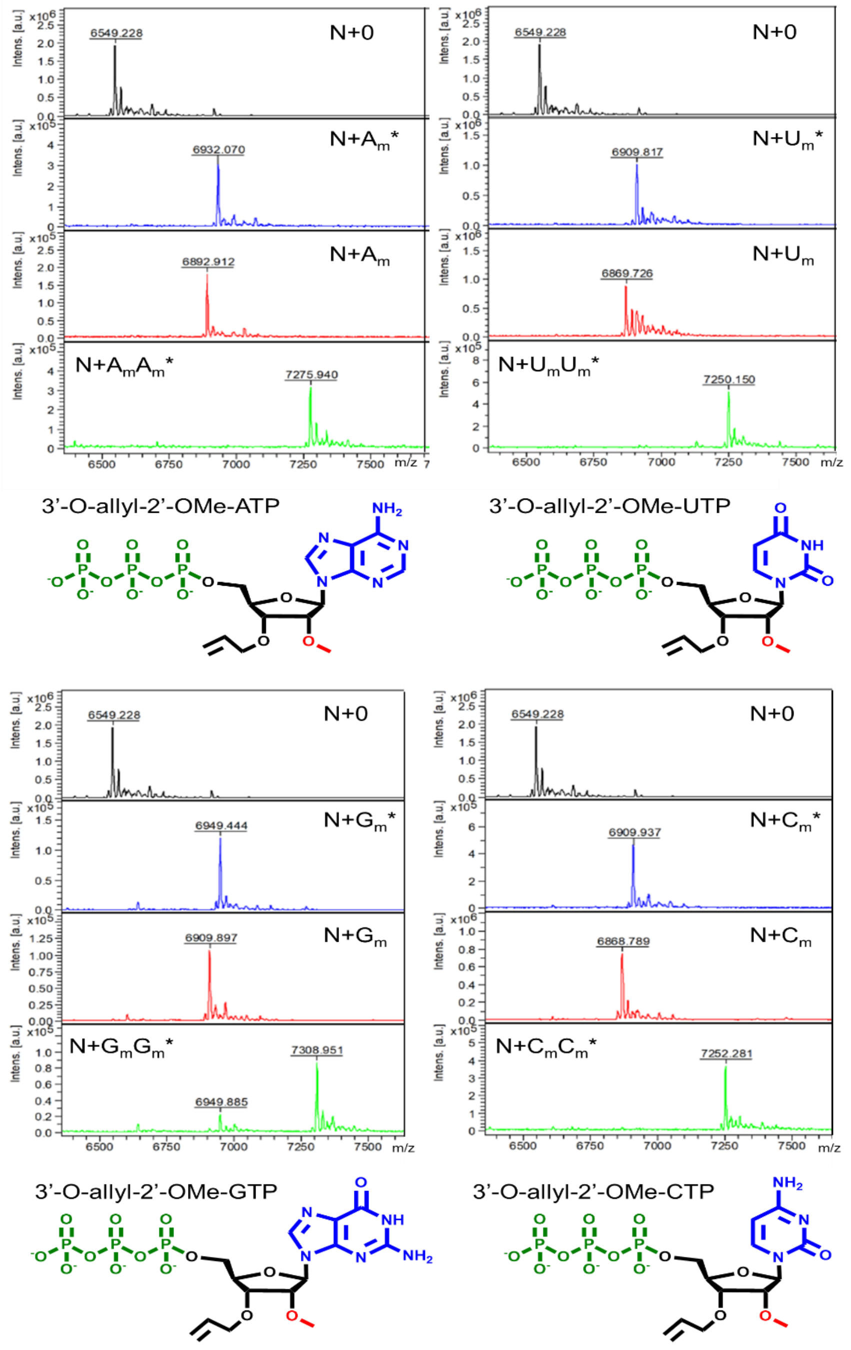


**Fig. S9**: An assessment of controlled, enzymatic oligonucleotide synthesis using the 2'-OMe modified 3'-O-allyl ether reversible terminator NTP set (A, U, G, C) with PUP mutant variant H336R. Extension reactions were carried out using standard reaction conditions using a Cy5 labeled 19-nt initiator oligonucleotide. MALDI-TOF mass spectrometry was used to assess the efficiency of initiator conversion to the **N+1*** product, allyl ether deblocking and then conversion to the **N+2*** product from **N+1**.

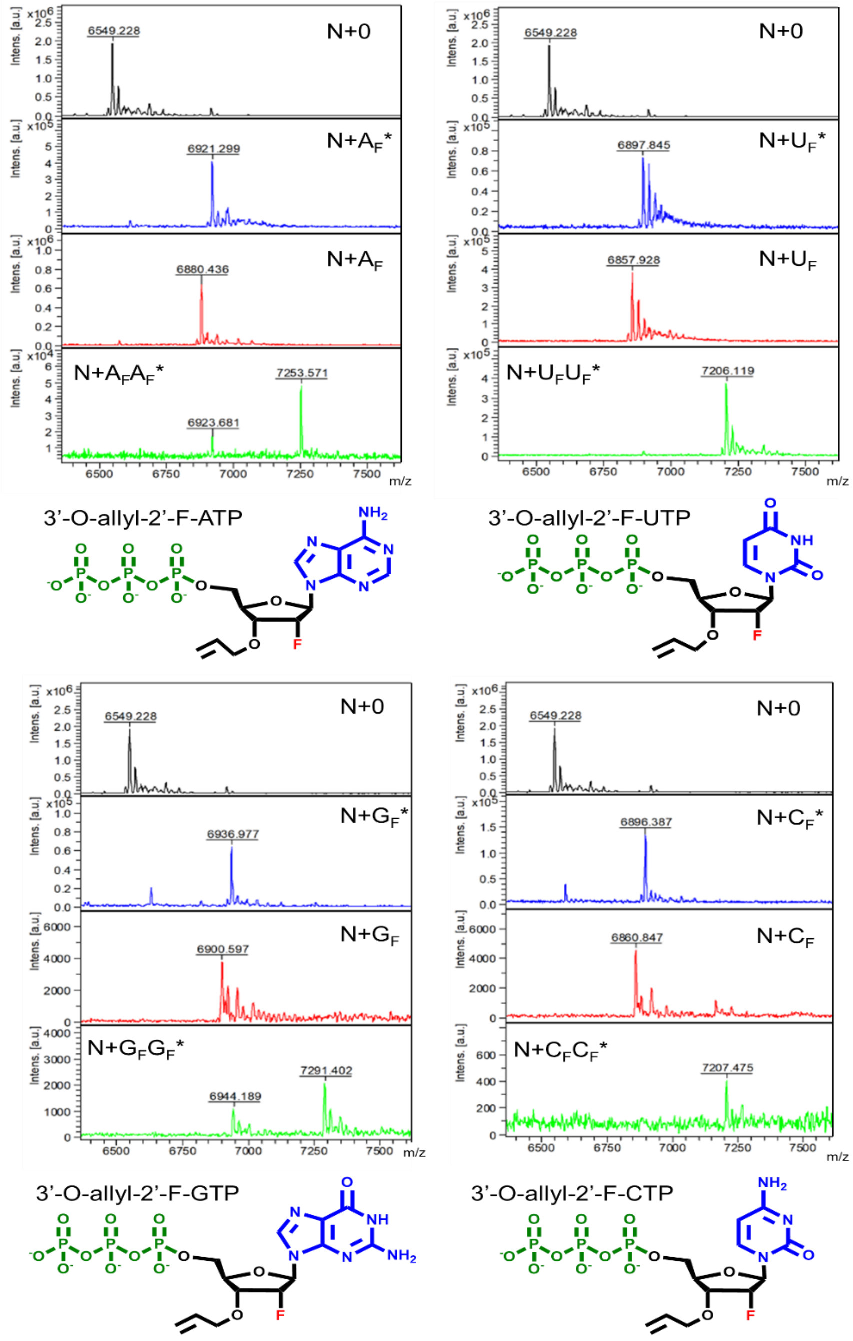


**Fig. S10**: An assessment of controlled, enzymatic oligonucleotide synthesis using the 2'-F modified 3'-O-allyl ether reversible terminator NTP set (A, U, G, C) with PUP mutant variant H336R. Extension reactions were carried out using standard reaction conditions using a Cy5 labeled 19-nt initiator oligonucleotide. MALDI-TOF mass spectrometry was used to assess the efficiency of initiator conversion to the **N+1*** product, allyl ether deblocking and then conversion to the **N+2*** product from **N+1**.

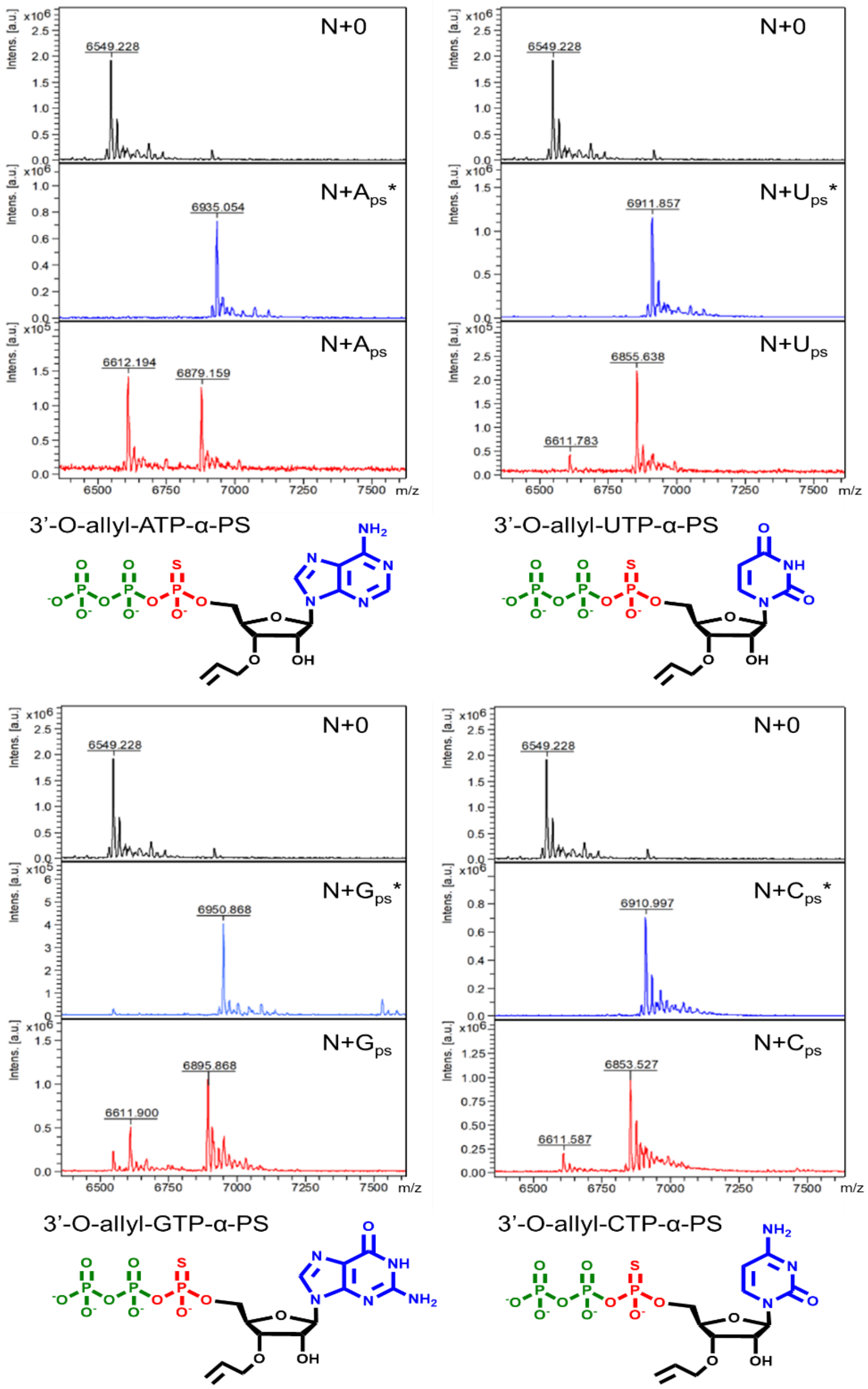


**Fig. S11**: An assessment of controlled, enzymatic oligonucleotide synthesis using the α-phosphorothioate (α-PS) modified 3'-O-allyl ether reversible terminator NTP set (A, U, G, C) with PUP mutant variant H336R. Extension reactions were carried out using standard reaction conditions using a Cy5 labeled 19-nt initiator oligonucleotide. MALDI-TOF mass spectrometry was used to assess the efficiency of initiator conversion to the **N+1*** product and allyl ether deblocking to **N+1**.

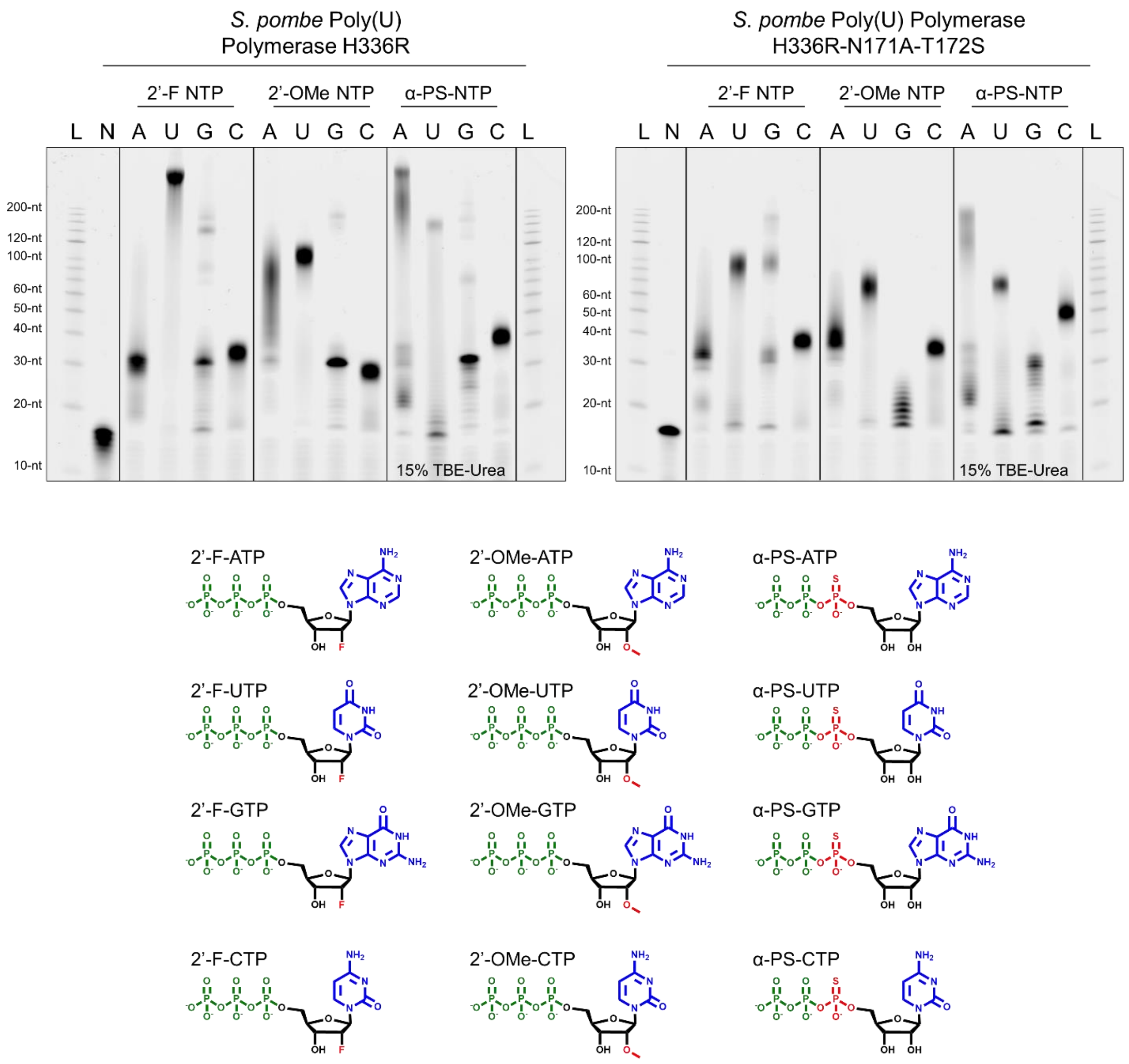


**Fig. S12:** An analysis of Poly(U) Polymerase mutant variant uncontrolled polymerization activity in the presence of unblocked 2'-fluoro (2'-F), 2'-methoxy (2'-OMe) and alpha-phosphorothioate (α-PS). Modified NTPs were incubated with both PUP mutant variants under standard reaction conditions using a Cy5 labeled 19-nt initiator oligonucleotide. The resultant extension products were assessed using denaturing gel electrophoresis using a 200-nt ssDNA ladder to measure the overall length of uncontrolled polymerization.

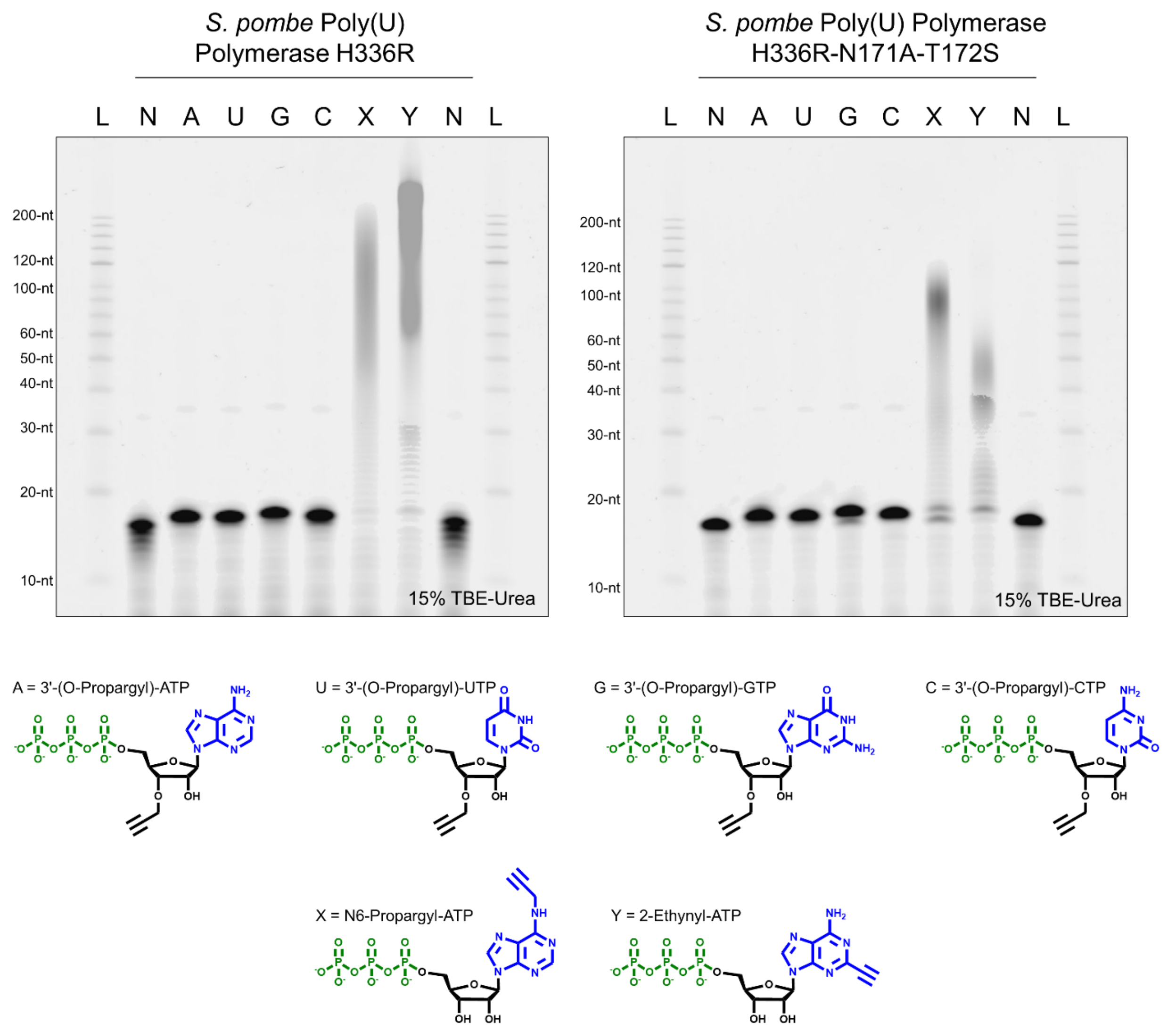


**Fig. S13**: An analysis of Poly(U) Polymerase mutant variant activity in the presence of propargyl modified nucleotides including a set of 3'- propargyl ether NTPs (A, U, G, C) as well as the base modified N6-propargyl-ATP and 2-Ethynyl-ATP (Jena Bioscience). Modified NTPs were incubated with both PUP mutant variants under standard reaction conditions using a Cy5 labeled 19-nt initiator oligonucleotide. Enzymatic activity was assessed using denaturing gel electrophoresis. Control reactions (N) contained all components except NTP. For the bases modified NTPs, a 200-nt ssDNA ladder (L) was used to measure the extent of uncontrolled polymerization.

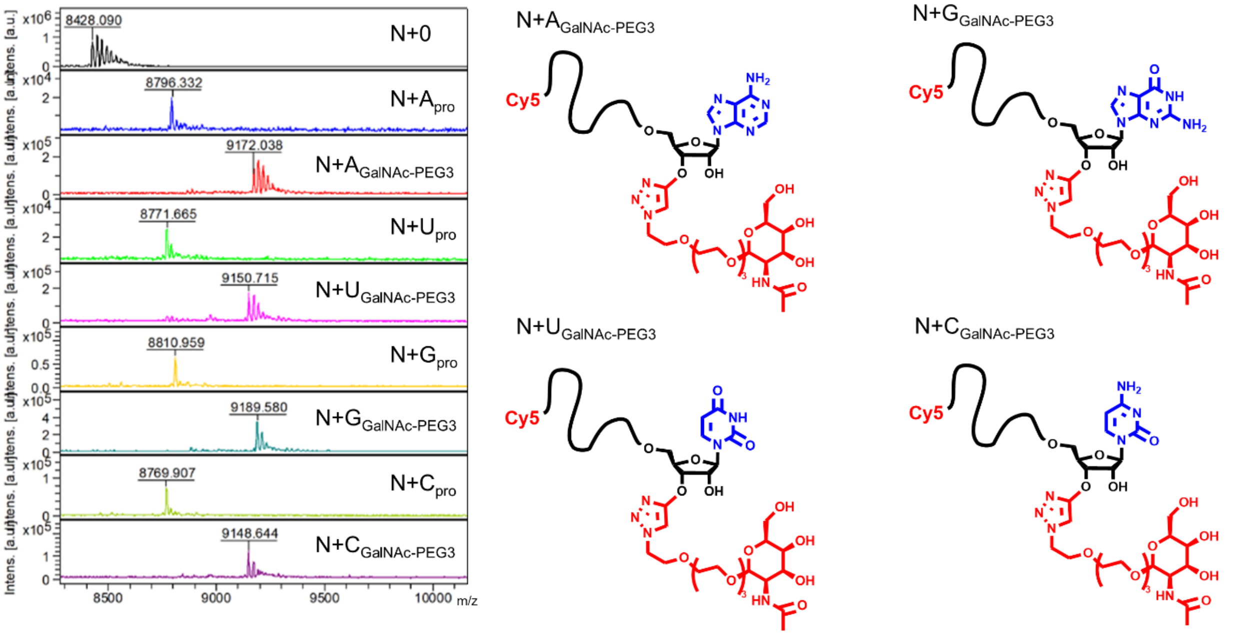


**Fig. S14**: An assessment of oligonucleotide labeling with an α-GalNAc-PEG3-Azide ligand with click chemistry following installation of a 3'- terminal propargyl group with PUP mutant variant H336R under standard reaction conditions and a 25-nt poly-U RNA initiator oligonucleotide modified with a 5'- Cy5. The complete set of 3'- propargyl ether NTPs (A, U, G, C) were evaluated in this activity. MALDI-TOF mass spectrometry was used to verify the conversion of the initiator oligonucleotide to the various **N+1$_{\mathbf{pro}}$** products as well as determine the extent of labeling with α-GalNAc-PEG3 to produce **N+1$_{\mathbf{GalNAc\text{-}PEG3}}$** oligonucleotides.

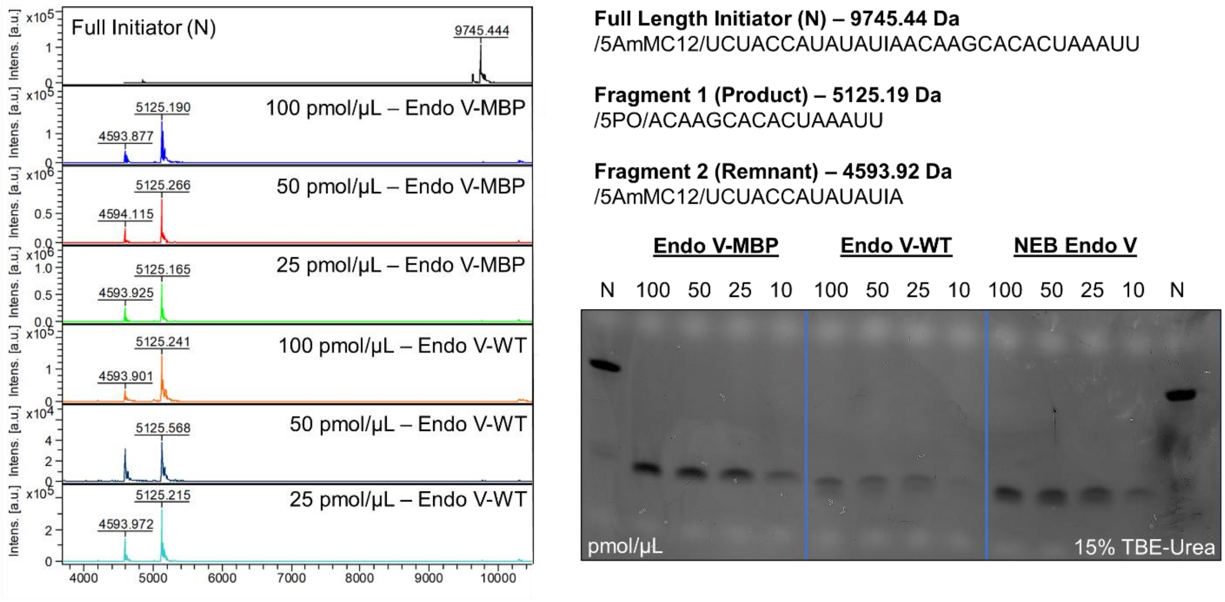


**Fig. S15**: An evaluation initiator oligonucleotide cleavage using Endonuclease V variants (wild-type and a fusion with an N-terminal maltose-binding protein (MBP)) expressed in-house and sourced commercially (NEB). MALDI-TOF mass spectrometry and denaturing gel electrophoresis was used to test cleavage robustness for each Endonuclease V variant. From the full-length initiator (N), it is expected that two fragments are formed: (1) the product at 5125.19 Da and (2) the remnant at 4593.92 Da. These were observed to have been formed when incubated with each Endonuclease V variant under standard cleavage reaction conditions. Similar results were found as the concentration of full-length initiator was increased from 25 pmol/µL to 100 pmol/µL (fixed volume at 10 µL and total input 250 pmol to 1000 pmol). No remaining full length initiator oligonucleotide or erroneous side products were observed in any of the Endonuclease V test cases.

**Variable Sequence – 5137.98 Da**
5'-PO-ACAAGCACACUAAAUUA-3'

**Dephosphorylated Variable Sequence – 5057.16 Da**
5'-OH-ACAAGCACACUAAAUUA-3

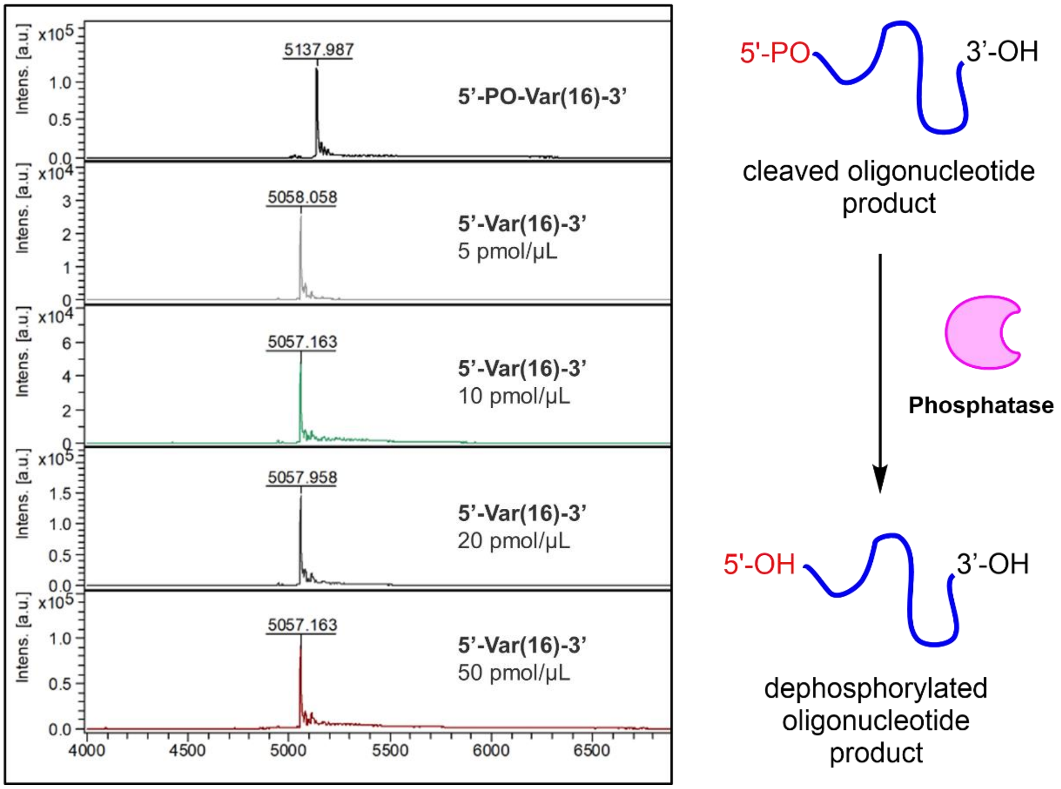


**Fig. S16:** A general scheme for the enzymatic removal of a 5'-phosphate (PO) group with a phosphatase from an oligonucleotide product that was cleaved from its initiator used in template-independent enzymatic synthesis. MALDI-TOF mass spectrometry was used to assess the dephosphorylation activity of Antarctic Phosphatase (NEB) in the presence of an RNA oligonucleotide sequence comprised of variable bases bearing a 5'-phosphate under standard reaction conditions. The expected dephosphorylated product was observed in each test case in which the total 5'-phosphate oligonucleotide concentration was increased from 5 pmol/μL to 50 pmol/μL.

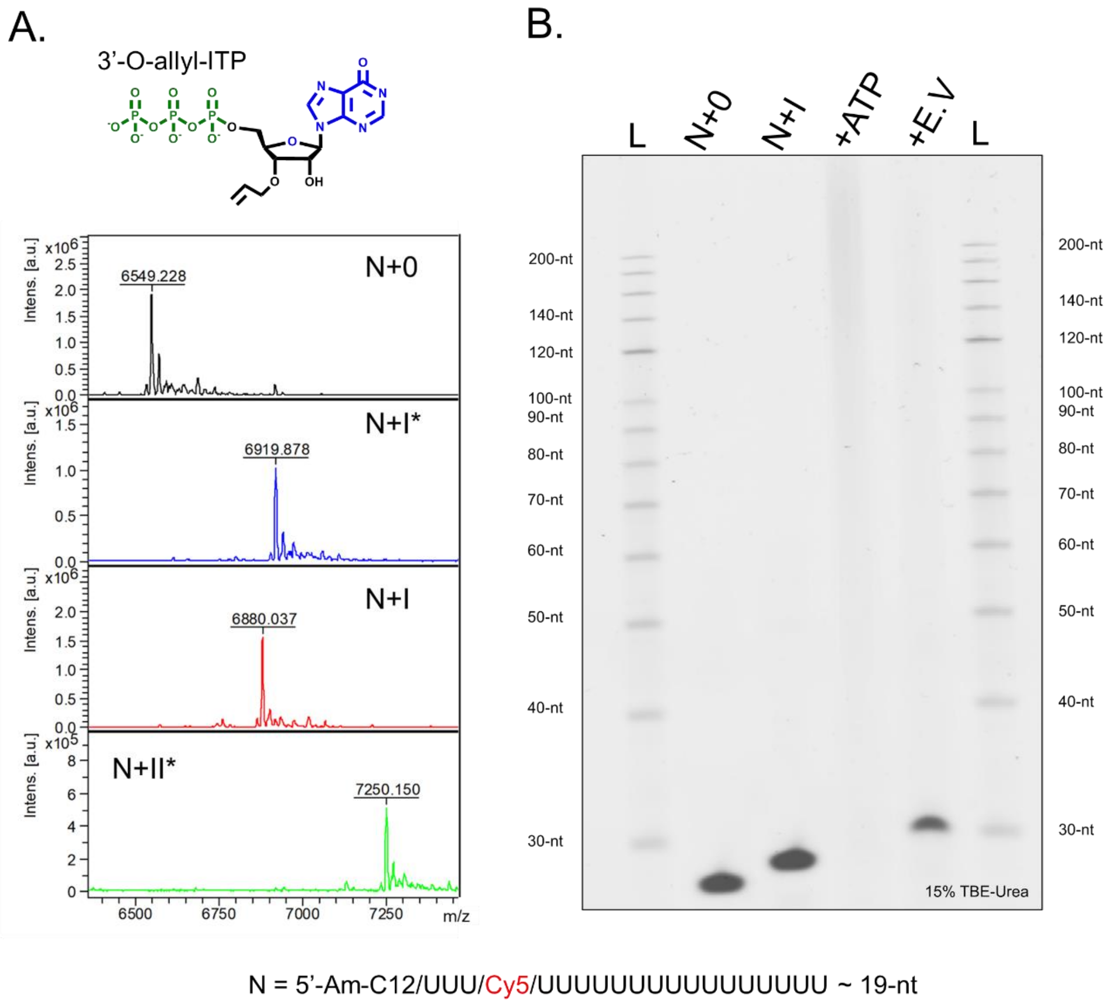


**Fig. S17: (A)** MALDI-TOF analysis of 3’-O- allyl ether ITP reversible terminator during a single cycle of enzymatic synthesis: initial incorporation (N+I*), deblocking (N+I), and generation of an N+2* single base transition product. **(B)** Following uncontrolled polymerization with unblocked ATP, Endonuclease V was used to cleave initiator oligonucleotide from long homopolymer sequence. Gel electrophoresis shows the unextended initiator as a negative control (N), initiator extended and deblocked N+I, which was subsequently used to generate the long +ATP homopolymer sequence that was digested with Endonuclease V to restore initiator oligonucleotide with sequence N+I-A. A 200-nt ssDNA ladder to measure the overall length of uncontrolled polymerization.

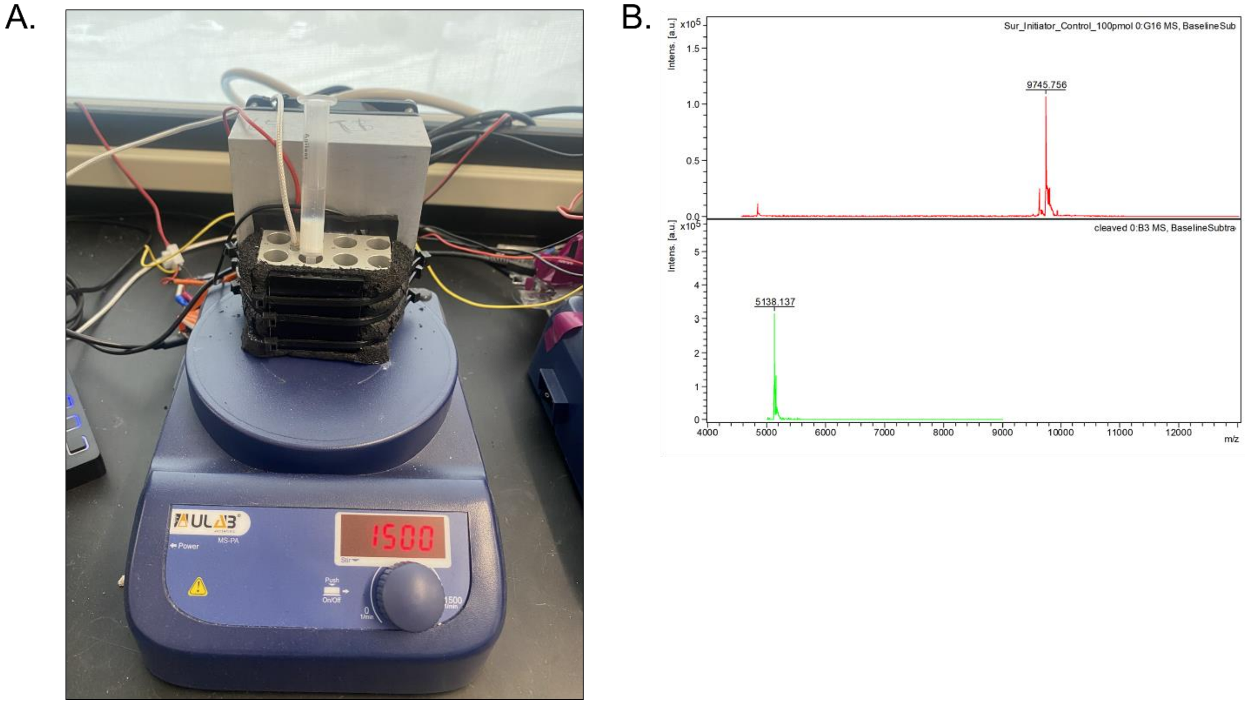


**Fig. S18: (A)** Picture of a stir reactor, which is comprised of a heat block controlled by a Pelletier unit and placed on top of a magnetic stir place. Solid support is placed within a 3 mL solid phase extraction column with a small magnetic flea and the appropriate reaction master mix for either extension, deblocking or Endonuclease V cleavage is added directly. Custom made software controls the heat block temperature, which is regulated by a probe drilled into the block. **(B)** MALDI-TOF analysis of Endonuclease V-mediated cleavage of the full surface initiator oligonucleotide.

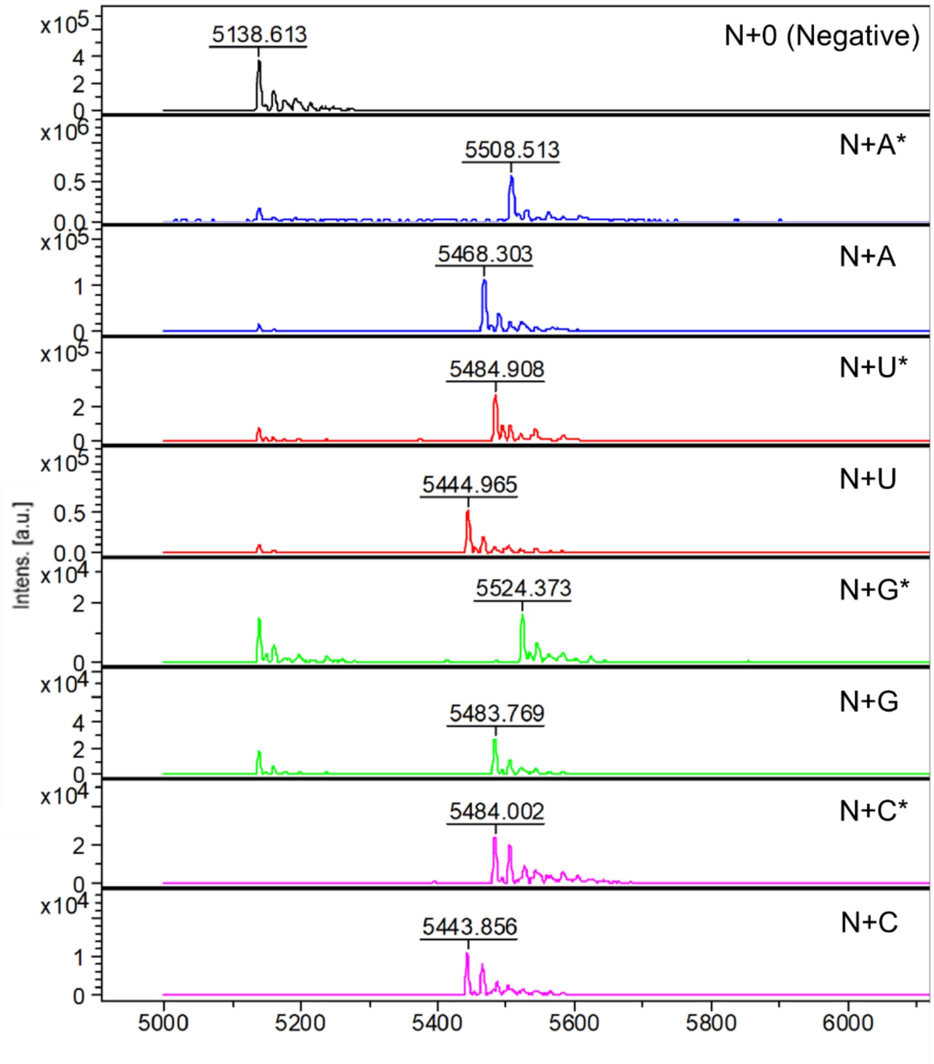


**Fig. S19**: MALDI-TOF analysis of oligonucleotide products cleaved from the CPG solid support via Endonuclease V after a single cycle of enzymatic extension and deblocking with the full 3'-O-allyl ether NTP set. The negative control reaction contained all extension reaction components with the exception of NTP building block and did not undergo deblocking.

**SEQ1: *S. pombe* Poly(U) Polymerase WT**

MNISSAQFIPGVHTVEEIEAEIHKNLHISKSCSYQKVPNSHKEFTKFCYEVYNEIKISDKEF
KEKRAALDTLRLCLKRISPDAELVAFGSLESGLALKNSDMDLCVLMDSRVQSDTIALQF
YEELIAEGFEGKFLQRARIPIIKLTSDTKNGFGASFQCDIGFNNRLAIHNTLLLSSYTKLDA
RLKPMVLLVKHWAKRKQINSPYFGTLSSYGYVLMVLYYLIHVIKPPVFPNLLLSPLKQE
KIVDGFDVGFDDKLEDIPPSQNYSSLGSLLHGFFRFYAYKFEPREKVVTFRRPDGYLTKQ
EKGWTSATEHTGSADQIIKDRYILAIEDPFEISHNVGRTVSSSGLYRIRGEFMAASRLLNS
RSYPIPYDSLFEEAPIPPRRQKKTDEQSNKKLLNETDGDNSE

**SEQ2: *S. pombe* Poly(U) Polymerase H336R**

MNISSAQFIPGVHTVEEIEAEIHKNLHISKSCSYQKVPNSHKEFTKFCYEVYNEIKISDKEF
KEKRAALDTLRLCLKRISPDAELVAFGSLESGLALKNSDMDLCVLMDSRVQSDTIALQF
YEELIAEGFEGKFLQRARIPIIKLTSDTKNGFGASFQCDIGFNNRLAIHNTLLLSSYTKLDA
RLKPMVLLVKHWAKRKQINSPYFGTLSSYGYVLMVLYYLIHVIKPPVFPNLLLSPLKQE
KIVDGFDVGFDDKLEDIPPSQNYSSLGSLLHGFFRFYAYKFEPREKVVTFRRPDGYLTKQ
EKGWTSATEHTGSADQIIKDRYILAIEDPFEIS**R**NVGRTVSSSGLYRIRGEFMAASRLLNS
RSYPIPYDSLFEEAPIPPRRQKKTDEQSNKKLLNETDGDNSE

**SEQ3: *S. pombe* Poly(U) Polymerase H336R-N171A-T172S**

MNISSAQFIPGVHTVEEIEAEIHKNLHISKSCSYQKVPNSHKEFTKFCYEVYNEIKISDKEF
KEKRAALDTLRLCLKRISPDAELVAFGSLESGLALKNSDMDLCVLMDSRVQSDTIALQF
YEELIAEGFEGKFLQRARIPIIKLTSDTKNGFGASFQCDIGFNNRLAIH**AS**LLLSSYTKLDA
RLKPMVLLVKHWAKRKQINSPYFGTLSSYGYVLMVLYYLIHVIKPPVFPNLLLSPLKQE
KIVDGFDVGFDDKLEDIPPSQNYSSLGSLLHGFFRFYAYKFEPREKVVTFRRPDGYLTKQ
EKGWTSATEHTGSADQIIKDRYILAIEDPFEIS**R**NVGRTVSSSGLYRIRGEFMAASRLLNS
RSYPIPYDSLFEEAPIPPRRQKKTDEQSNKKLLNETDGDNSE

**SEQ4:** ***E. coli*** **Endonuclease V WT**

MDLASLRAQQIELASSVIREDRLDKDPPDLIAGADVGFEQGGEVTRAAMVLLKYPSLEL
VEYKVARIATTMPYIPGFLSFREYPALLAAWEMLSQKPDLVFVDGHGISHPRRLGVASH
FGLLVDVPTIGVAKKRLCGKFEPLSSEPGALAPLMDKGEQLAWVWRSKARCNPLFIATG
HRVSVDSALAWVQRCMKGYRLPEPTRWADAVASERPAFVRYTANQP

**SEQ5:** ***E. coli*** **Endonuclease V WT – N-Term. MBP Fusion**

MDLASLRAQQIELASSVIREDRLDKDPPDLIAGADVGFEQGGEVTRAAMVLLKYPSLEL
VEYKVARIATTMPYIPGFLSFREYPALLAAWEMLSQKPDLVFVDGHGISHPRRLGVASH
FGLLVDVPTIGVAKKRLCGKFEPLSSEPGALAPLMDKGEQLAWVWRSKARCNPLFIATG
HRVSVDSALAWVQRCMKGYRLPEPTRWADAVASERPAFVRYTANQPMKIEEGKLVIW
INGDKGYNGLAEVGKKFEKDTGIKVTVEHPDKLEEKFPQVAATGDGPDIIFWAHDRFGG
YAQSGLLAEITPDKAFQDKLYPFTWDAVRYNGKLIAYPIAVEALSLIYNKDLLPNPPKT
WEEIPALDKELKAKGKSALMFNLQEPYFTWPLIAADGGYAFKYENGKYDIKDVGVDNA
GAKAGLTFLVDLIKNKHMNADTDYSIAEAAFNKGETAMTINGPWAWSNIDTSKVNYGV
TVLPTFKGQPSKPFVGVLSAGINAASPNKELAKEFLENYLLTDEGLEAVNKDKPLGAVA
LKSYEEELAKDPRIAATMENAQKGEIMPNIPQMSAFWYAVRTAVINAASGRQTVT

**Preparation of 3’-O- allyl ether NTPs (A, U, G, C)**

**Scheme I. 3’-O- allyl ether ATP**

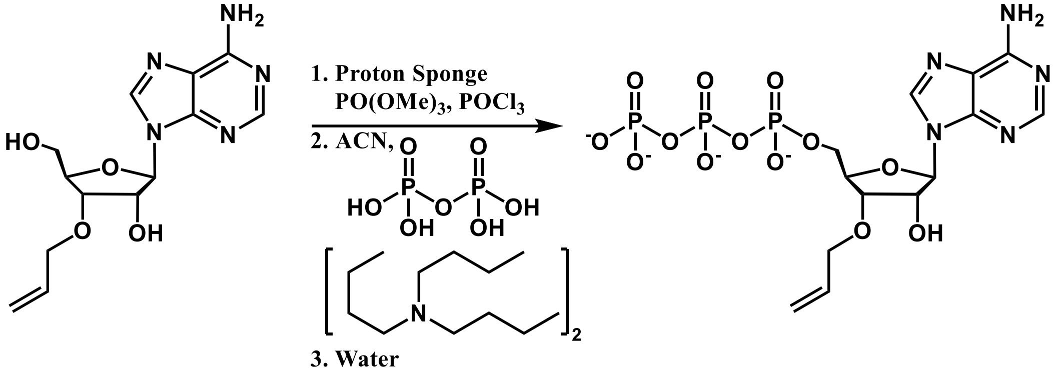


The mixture of nucleoside (molecular weight: 307.3 g/mol, 1.04 g , 3.38 mmol) and proton sponge (molecular weight: 214.31, 2.03 g, 9.47 mmol) were prepared as described in the general procedure. This mixture was dissolved in trimethoxy phosphate (25.0 mL) and was cooled to -5 $^{o}$C. This was followed by the slow addition of phosphoryl oxychloride (molecular weight: 153.32, density: 1.64 g/cm$^3$, 0.35 mL, 3.7 mmol). After 3 minutes, another portion of phosphoryl oxychloride (molecular weight: 153.32, density: 1.64 g/cm$^3$, 0.1 mL, 1.1 mmol) was added. After stirring for 10 minutes a pre-chilled mixture of tributyl ammonium pyrophosphate (Molecular Weight: 548.68, 7.0 g, 12.8 mmol), acetonitrile (55 mL), and tributyl amine (12 mL) was quickly added to the reaction. This was allowed to stir for two hours and slowly warm to room temperature. The reaction was quenched by the addition of water (*circa*. 150 mL) and the reaction was worked up, isolated and purified according to the general procedure (yield 33%, formula weight for the tetra-triethylammonium salt: 951.5 g/mol, 1.65 g, 1.11 mmol). $^1$H NMR (400 MHz, $D_2O$) δ 8.43 (s, 1H), 8.15 (s, 1H), 6.04 (d, *J* = 6 Hz, 1H), 5.96 (m, *J* = 17, 11 Hz, 1H), 5.37 (dd, *J* = 17, 2 Hz, 1H), 5.23 (dd, *J* = 11, 2 Hz, 1H), 4.80 (t, *J* = 6 Hz, 1H), 4.42 (t, *J* = 3 Hz, 1H), 4.30 (dd, *J* = 6, 3 Hz, 1H), 4.19 (d, *J* = 6 Hz, 2H), 4.15 (t, *J* = 3 Hz, 2H), 3.09 (q, 16 H), 1.18 (t, 24H). $^{31}$P NMR (400 MHz, $D_2O$) δ – 10.6 (d, *J* = 20, 1H), - 11.5 (d, *J* = 20, 1H), - 23.1 (t, *J* = 20). ESI-MS: calculated for $[C_{13}H_{21}N_5O_{13}P_3]^+$ = 548.0343, found: 548.0347.

**Scheme II. 3'-O- allyl ether UTP**

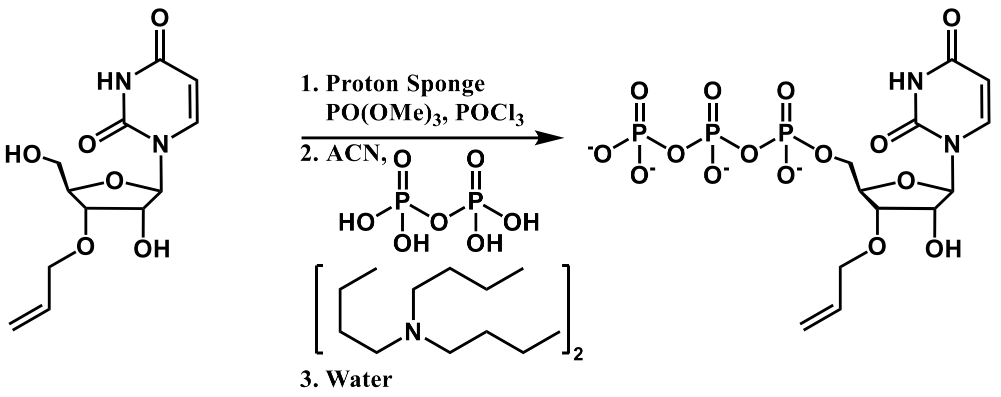


The mixture of nucleoside (molecular weight: 284.10 g/mol, 0.57 mg, 2.00 mmol) and proton sponge (molecular weight: 214.31, 1.00 g, 4.67 mmol) were prepared as described in the general procedure. This mixture was dissolved in trimethoxy phosphate (9.0 mL) and was cooled to -5 $^oC$. This was followed by the slow addition of phosphoryl oxychloride (molecular weight: 153.32, density: 1.64 g/cm$^3$, 0.2 mL, 2.1 mmol). After 3 minutes, another portion of phosphoryl oxychloride (molecular weight: 153.32, density: 1.64 g/cm$^3$, 0.1 mL, 1.1 mmol) was added. After stirring for 10 minutes a pre-chilled mixture of tributyl ammonium pyrophosphate (Molecular Weight: 548.68, 3.7 g, 6.7 mmol), acetonitrile (32 mL), and tributyl amine (6 mL) was quickly added to the reaction. This was allowed to stir for two hours and slowly warmed to room temperature. The reaction was quenched by the addition of water (*circa*. 200 mL) and the reaction was worked up, isolated, and purified according to the general procedure (yield 29 %, formula weight for the tetra-triethylammonium salt: 928.5 g/mol, 0.54 g, 0.58 mmol). $^1$H NMR (400 MHz, $D_2O$) δ 7.88 (d, *J* = 8 Hz, 1H), 5.93 (m, 3 H), 5.32 (dd, *J* = 17, 2 Hz, 1H), 5.22 (dd, *J* = 11, 2 Hz, 1H), 4.40 (t, *J* = 6 Hz, 1H), 4.32 (m , *J* = 3 Hz, 1H), 4.15 (m, *J* = 3 Hz, 3H), 3.12 (q, 19 H), 1.20 (t, 29 H). $^{31}$P NMR (400 MHz, $D_2O$) δ -10.8 (d, *J* = 20, 1H), -11.6 (d, *J* = 20, 1H), -23.3 (t, *J* = 20, 1H). ESI-MS: calculated for $[C_{12}H_{18}N_2O_{15}P_3]^-$ = 522.9926 , found: 522.9928.

**Scheme III. 3'-O- allyl ether GTP**

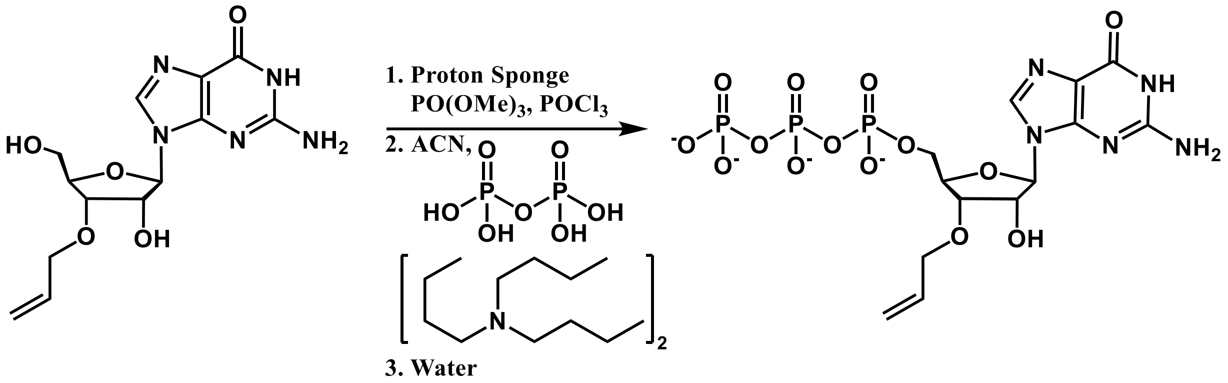

The mixture of nucleoside (molecular weight: 323.12 g/mol, 1.1 g, 3.40 mmol) and proton sponge (molecular weight: 214.31, 2.21 g, 10.3 mmol) were prepared as described in the general procedure. This mixture was dissolved in trimethoxy phosphate (25.0 mL) and was cooled to -5 $^oC$. This was followed by the slow addition of phosphoryl oxychloride (molecular weight: 153.32, density: 1.64 g/cm$^3$, 0.35 mL, 3.7 mmol). After 3 minutes, another portion of phosphoryl oxychloride (molecular weight: 153.32, density: 1.64 g/cm$^3$, 0.1 mL, 1.1 mmol) was added. After stirring for 10 minutes a pre-chilled mixture of tributyl ammonium pyrophosphate (molecular weight: 548.68, 7.0 g, 12.8 mmol), acetonitrile (55 mL), and tributyl amine (12 mL) and was quickly added to the reaction. This was allowed to stir for two hours and slowly warmed to room temperature. The reaction was quenched by the addition of water (*circa*. 200 mL) and the reaction was worked up, isolated, and purified according to the general procedure (yield 40 %, formula weight for the tetra-triethylammonium salt: 967.5 g/mol, 1.33 g, 1.37 mmol). $^1H$ NMR (400 MHz, $D_2O$) δ 8.03 (s, 1H), 5.96 (ddt, *J* = 6 Hz, 1H), 5.84 (d, *J* = 7 Hz, 1H), 5.35 (dd, *J* = 17 Hz, 1H), 5.23 (dd, *J* = 11 Hz, 1H), 4.84 (dd, *J* = 7 Hz, 1H), 4.39 (m, *J* = 3 Hz, 1H), 4.29 (dd, *J* = 5, 3 Hz, 1H), 4.19 (d, *J* = 6 Hz, 2H), 4.15 (m, *J* = 6 Hz, 2H), 3.11 (q, 19H), 1.19 (t, 29H). $^{31}P$ NMR (400 MHz, $D_2O$) δ -10.6 (d, *J* = 20 Hz, 1H), -11.5 (d, *J* = 20 Hz, 1H), -23.1 (t, *J* = 20 Hz, 1H). ESI-MS: calculated for $[C_{13}H_{19}N_5O_{14}P_3]^-$ = 562.0147, found: 562.0150.

**Scheme IV. 3’-O- allyl ether CTP:**

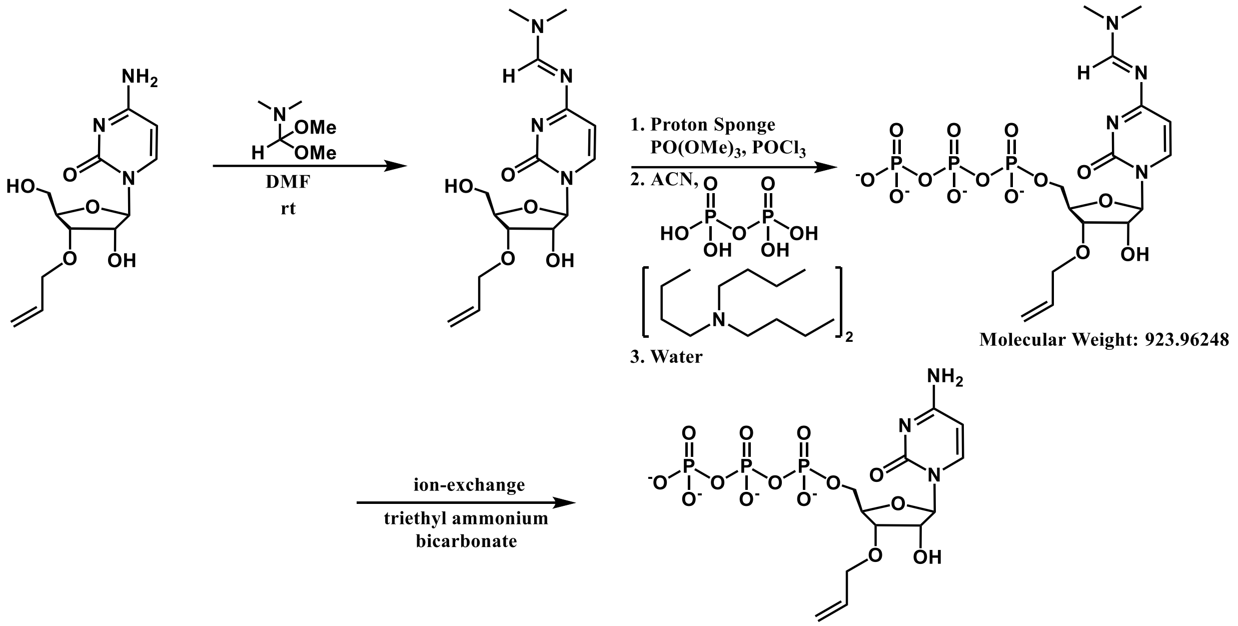


The nucleoside (molecular weight: 283.3 g/mol, 0.22 g, 0.78 mmol) was dissolved in a mixture of DMF and 1,1-dimethoxytrimethylamine (*circ*. 6:1, 2.1 mL) and allowed to stir for 36 hours. The crude product was concentrated *in vacuo*. To the resultant solid was added dry proton sponge (molecular weight: 214.3, 10.45 g, 2.1 mmol) and the mixture was dried on the lyophilizer overnight in the reaction flask. Under a blanket of argon, the mixture was dissolved in trimethoxy phosphate (5.0 mL). This solution was cooled to -5°C and phosphoryl oxychloride was added (0.07 mL, 0.75 mmol) after 3 minutes another portion of phosphoryl oxychloride (0.03 mL, 0.32 mmol) was added. This solution was allowed to stir in the cold bath for about 20 minutes. After which time, pre-chilled tributyl ammonium pyrophosphate (1.4 g, 2.6 mmol) and tributyl amine (2.4 mL) in acetonitrile (11 mL) was added in one portion. This was allowed to stir for two hours and slowly warm to room temperature. The reaction was quenched by the addition of water and the reaction was worked up, isolated, and purified according to the general procedure (yield 39 %, formula weight for the tetra-triethylammonium salt: 927.5 g/mol, 283.0 mg, 0.31 mmol). $^{1}$H NMR (400 MHz, $D_2O$) δ 7.90 (d, *J* = 8 Hz, 1H), 6.09 (d, *J* = 8 Hz, 1H), 5.93 (m, *J* = 6, 5 Hz, 2H), 5.32 (dd, *J* = 17, 2 Hz, 1H), 5.22 (dd, *J* = 10, 2 Hz, 1H), 4.36 (t, *J* = 5 Hz, 1H), 4.30 (dt, *J* = 3 Hz, 1H), 4.22 (ddd, *J* = 3 Hz, 1H), 4.14 (m, 2H), 3.13 (q, 11H), 1.21 (t, 18H). $^{31}$P NMR (400 MHz, $D_2O$) δ

-10.42 (d, *J* = 20 Hz, 1H), -11.47 (d, *J* = 20 Hz, 1H), -23.12 (t, *J* = 20 Hz, 1H). ESI-MS: calculated for $[C_{12}H_{19}N_3O_{14}P_3]^-$ = 522.0085, found: 522.0089.

**3’-O- allyl ether ATP – Proton NMR**

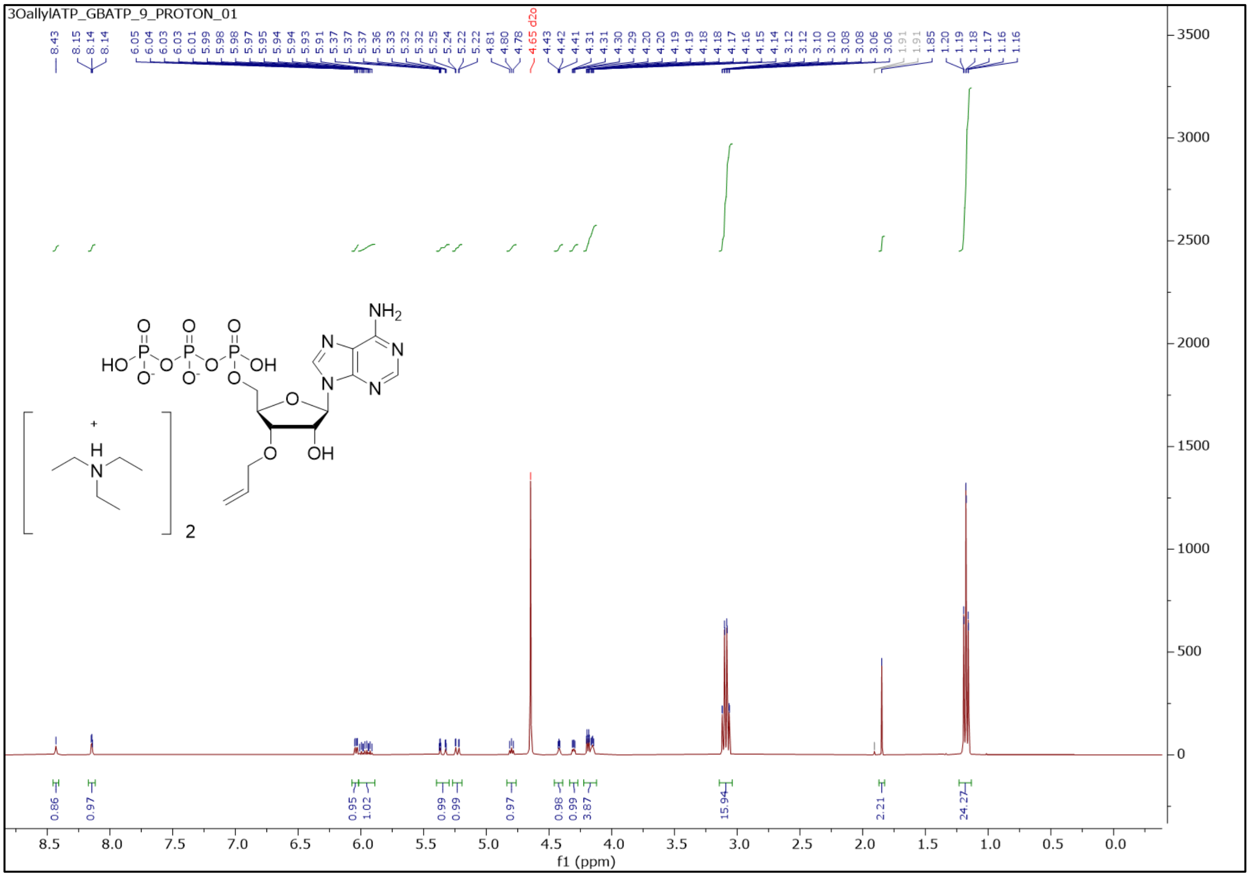

#### 3'-O- allyl ether ATP – Phosphorous NMR

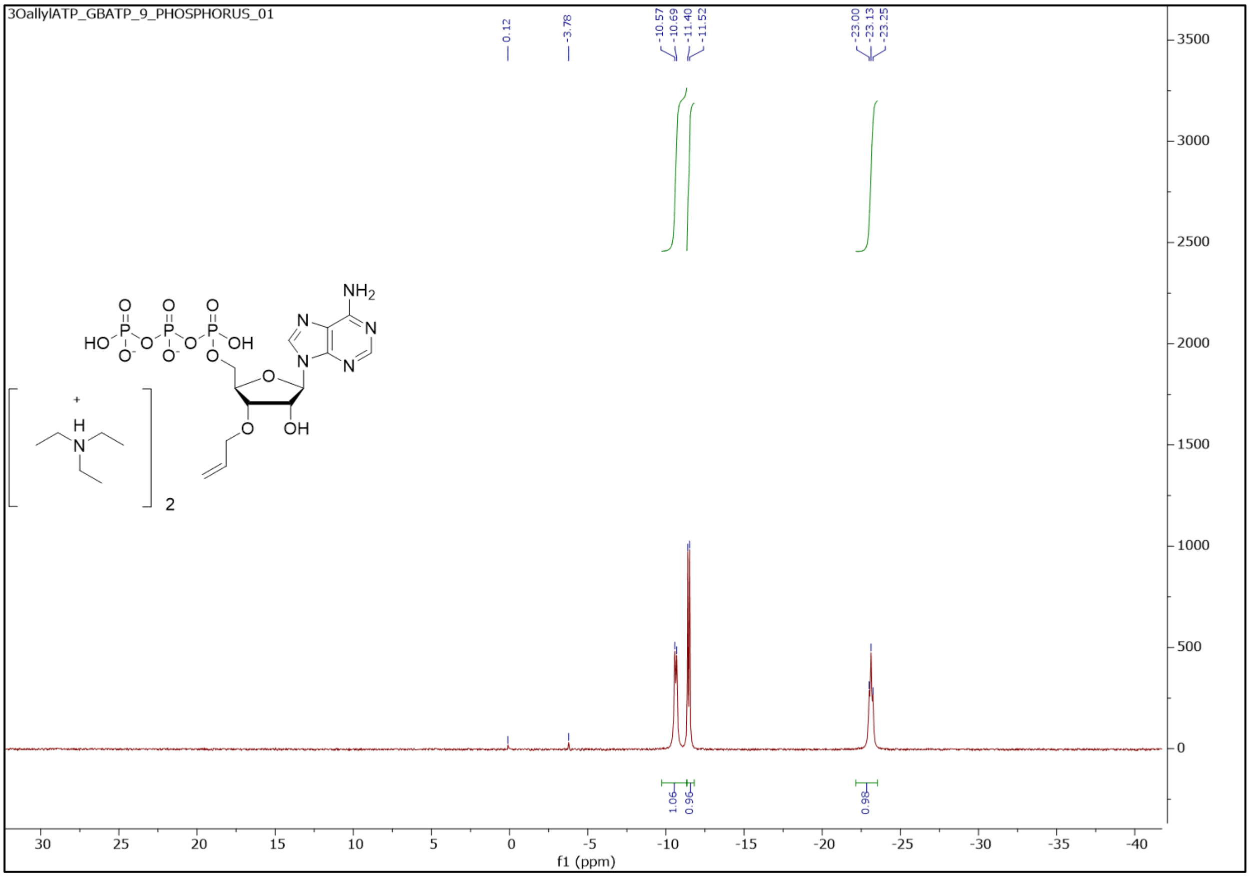

**3’-O- allyl ether UTP – Proton NMR**

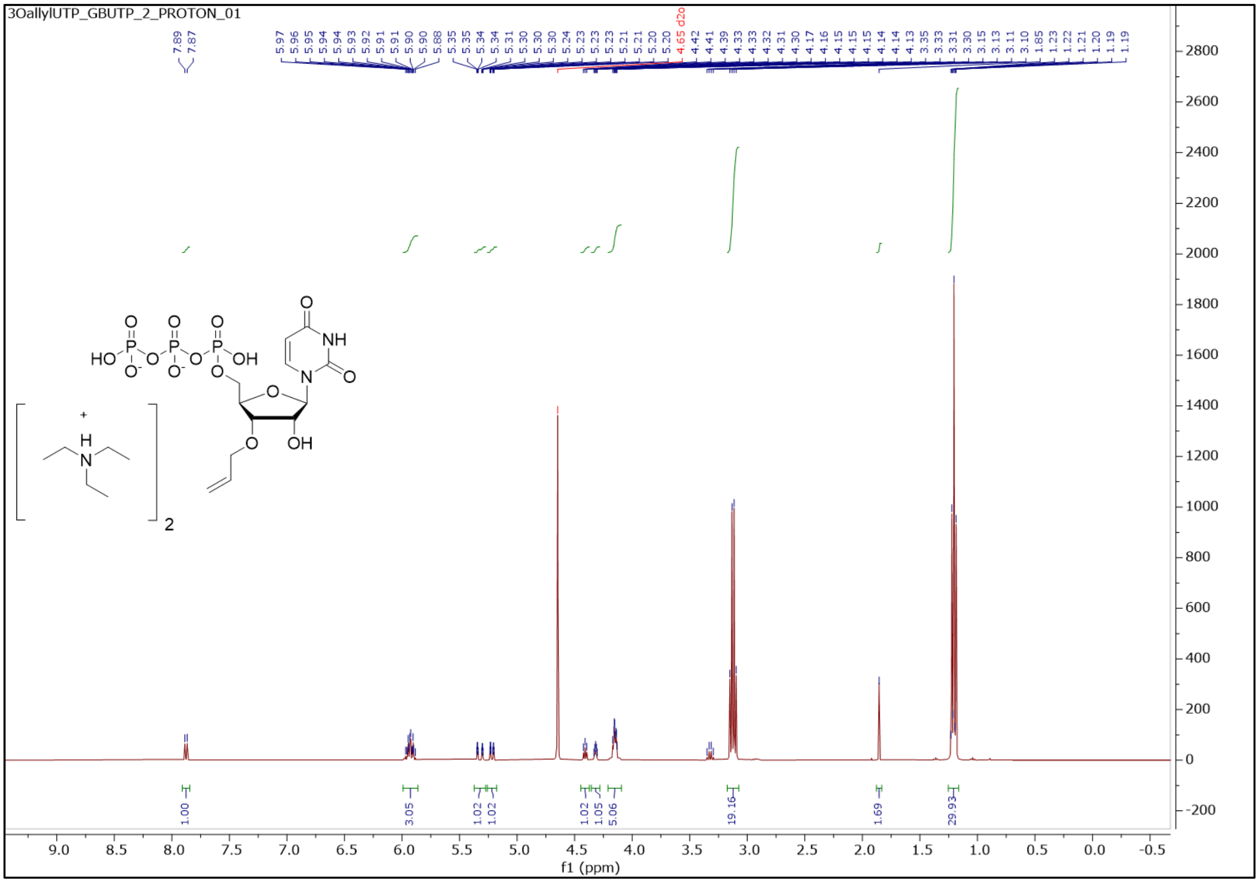

**3’-O- allyl ether UTP – Phosphorous NMR**

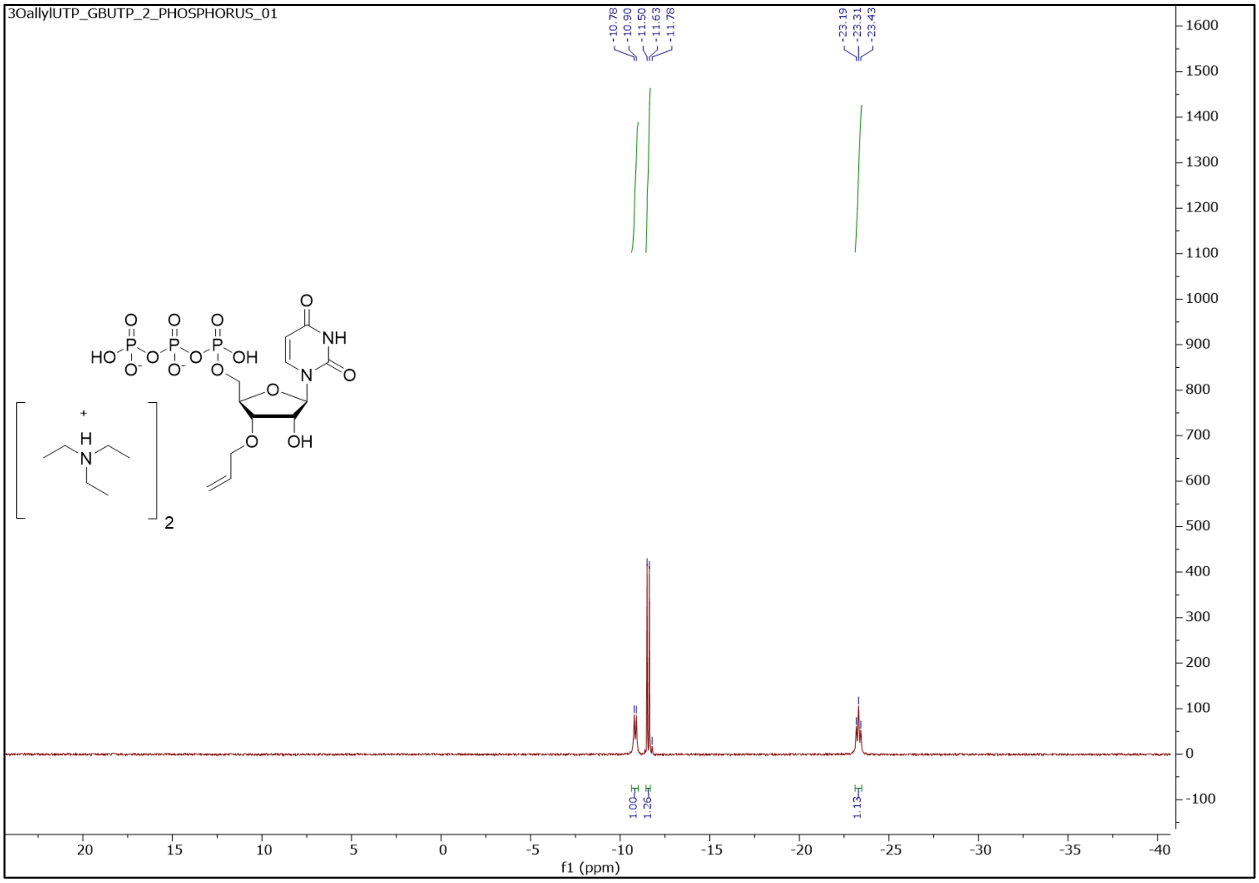

**3'-O- allyl ether GTP – Proton NMR**

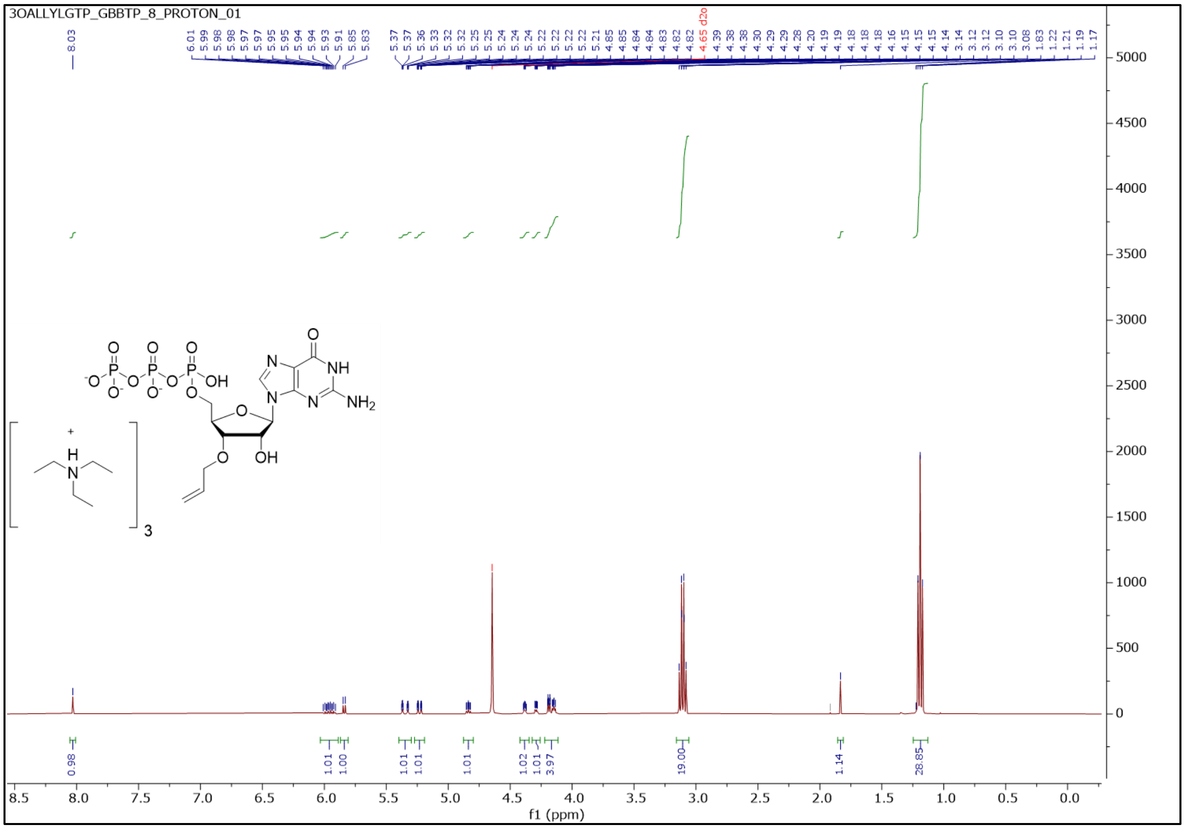

**3’-O- allyl ether GTP – Phosphorous NMR**

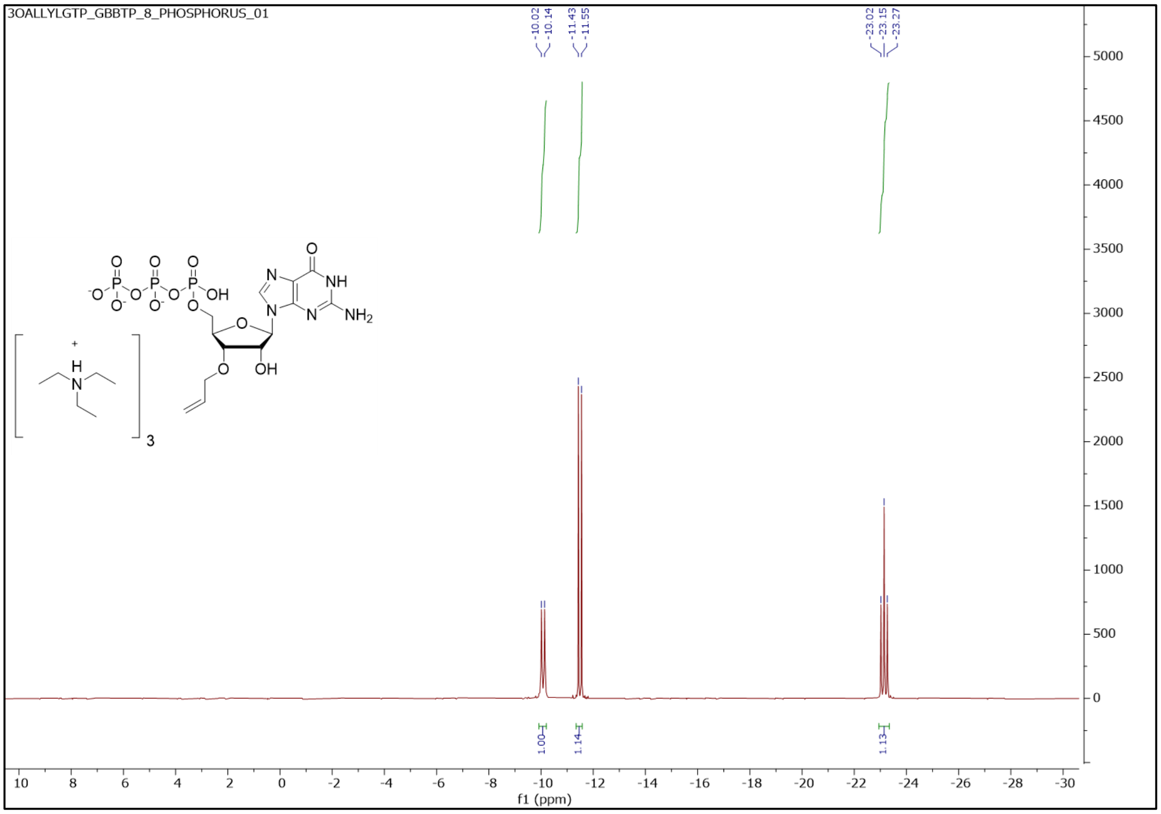

**3’-O- allyl ether CTP – Proton NMR**

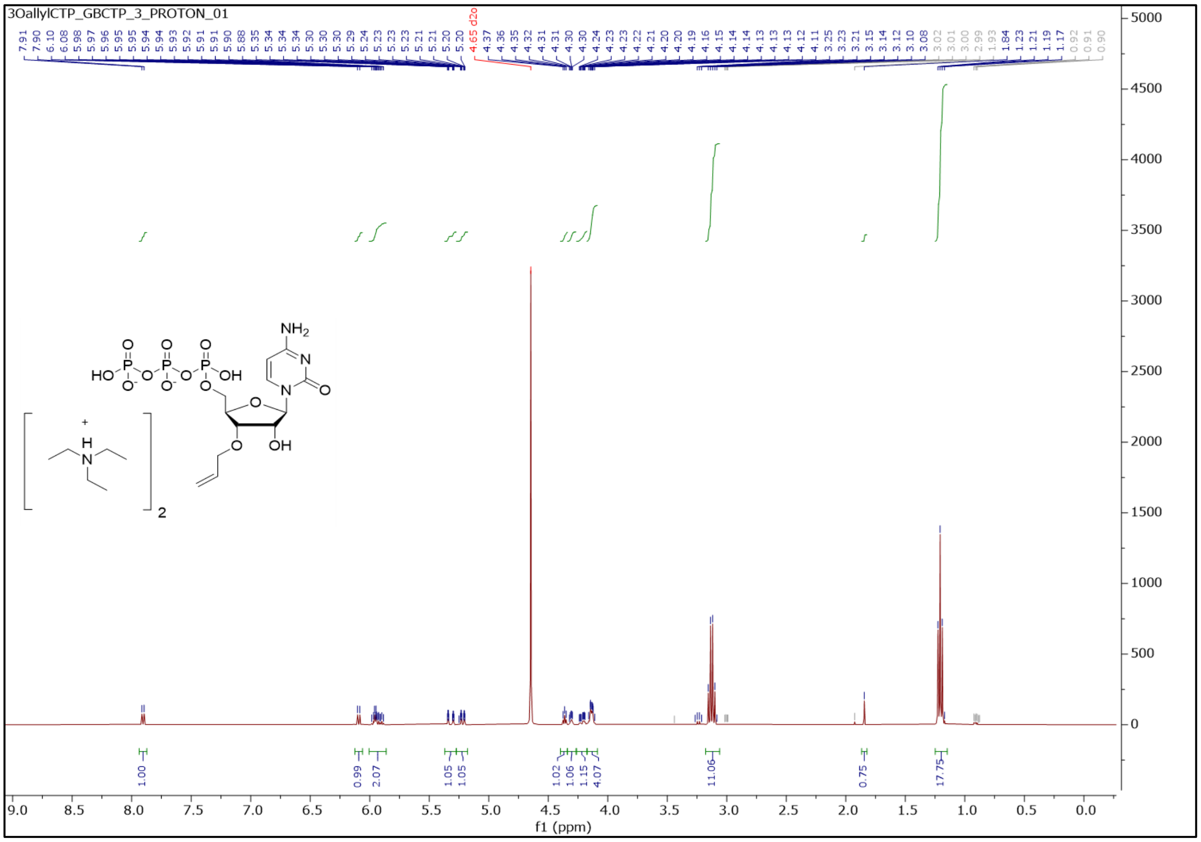

#### 3’-O- allyl ether CTP – Phosphorous NMR

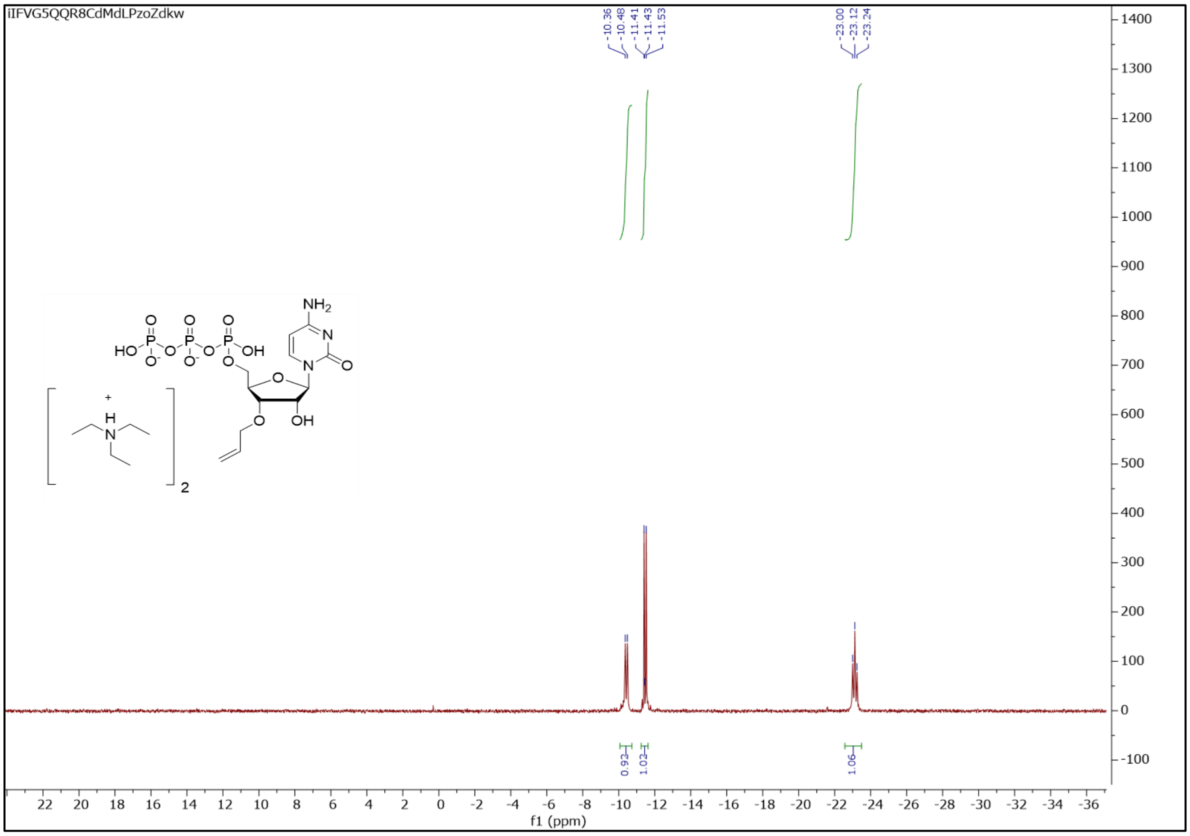

**References**

50. A. R. Kore, M. Shanmugasundaram, A. Senthilvelan, B. Srinivasan, An improved protection-free one-pot chemical synthesis of 2’-deoxynucleoside-5’-triphosphates. *Nucleosides Nucleotides Nucleic Acids*. **31**, 423–431 (2012).
